# Reproducible Research: Computational Design of Personalized Clinical Treatments for Walking Impairments Caused by Knee Osteoarthritis and Stroke

**DOI:** 10.64898/2026.03.02.709099

**Authors:** Robert M. Salati, Geng Li, Spencer Williams, Benjamin J. Fregly

## Abstract

**Background:** Personalized computational neuromusculoskeletal models have great potential for optimizing the design of clinical treatments for movement impairments. While many software tools address specific parts of the model personalization and treatment optimization processes, they typically require significant programming experience to use and do not cover the full breadth of these two processes. Furthermore, published neuromusculoskeletal modeling studies typically do not provide all the methodological details needed for others to reproduce the work. Consequently, researchers seeking to develop skills in the model personalization and treatment optimization processes face a steep learning curve due to the lack of detailed training materials that demonstrate both processes for real-life clinical problems using real-life subject movement data.

**Methods:** This article presents detailed training tutorials for the model personalization and treatment optimization processes using two real-life clinical problems and the Neuromusculoskeletal Modeling (NMSM) Pipeline. The first clinical problem involves the design of personalized gait modifications and high tibial osteotomy surgery for an individual with bilateral medial knee osteoarthritis, where the goal is to reduce the peak adduction moment in both knees to a specified target level. The second clinical problem involves the design of a synergy-based functional electrical stimulation prescription for an individual post-stroke with impaired walking function, where the goal is to equalize the propulsive and braking impulses between the two legs. Both tutorials were implemented as course projects given to novice users in a combined undergraduate/graduate mechanical engineering course.

**Results:** Both tutorials produced personalized neuromusculoskeletal models and associated dynamically consistent tracking optimizations that closely reproduced subject-specific experimental joint angles, joint moments, ground reaction forces and moments, and (if applicable) muscle activations measured during walking. Subsequent design optimizations predicted personalized treatments that achieved target values of peak knee adduction moments or propulsive and braking impulses.

**Conclusions:** The detailed step-by-step tutorials presented with this article are the first to walk users through the entire process of creating personalized neuromusculoskeletal models and then using them to design personalized treatments for two subjects with two different clinical problems. These tutorials can be used to introduce new users to the NMSM Pipeline and as projects in neuromusculoskeletal modeling courses.

## Introduction

Neuromusculoskeletal (NMS) modeling has great potential for informing the clinical treatment design process. For NMS models to be used effectively to design treatments for movement impairments, they need to be personalized to each individual, where the aspects of an individual’s model that require personalization will depend on the clinical problem being addressed (1). For some clinical problems, personalized skeletal geometry developed from the individual’s imaging data may be necessary (2–11). Several software tools exist that can personalize skeletal geometry either with the use of imaging data (e.g., NMS Builder (12), The MAP Client (13–15), OpenSim Creator (16), Bone Deformation Tool (17), STAPLE (18), the workflow presented in (19)) and without the use of imaging data (e.g., The MAP Client (13), Torsion Tool (20)). For other clinical problems, personalized joint structure, muscle-tendon, neural control, and foot-ground contact model components calibrated using the individual’s movement, force, and electromyographic (EMG) data may be necessary (3,21–28). Several tools exist that can personalize some, but not all, of these model components (e.g., AddBiomechanics (29), CEINMS (30), CEINMS-RT (31), SimCP (32)). Consequently, creation of a personalized neuromusculoskeletal model possessing these personalized model components is either not possible or challenging due to the need to use a combination of tools.

Once individual-specific NMS models are available, they can be used to predict computationally how the individual will function following a planned clinical intervention (3,21,23,33). This computational process, termed “predictive simulation,” is itself challenging, historically requiring specific skills in computational simulation and numerical methods to perform. To date, most predictive simulations of gait using NMS models have been performed using in-house software (21,34–42), but such tools are difficult for other researchers to use for their own research purposes. Consequently, several research groups have recently developed software tools that make the predictive simulation process easier to learn, facilitating replication of research performed by others. OpenSim Moco (43–45) has made predictive simulation much more accessible within the OpenSim simulation environment, while other software tools facilitate generation of predictive simulations either partially or completely outside of the OpenSim environment (e.g., BioMAC-Sim-Toolbox (46), SCONE (47), OpenSim-MATLAB Optimal Control (48), MyoSuite (49), GaitDynamics (50)). Each of these tools can be used to generate complex simulations that predict the functional outcome of a planned treatment, but none of them were designed to interface directly with model personalization software. For this reason, nearly all published predictive simulations of human movement have utilized scaled generic models that predict generic movements rather than personalized models that predict individual subject movements.

For model personalization and predictive simulation alike, training new researchers to use these valuable software tools poses a significant challenge. If a new researcher wants to generate predictive simulations of an individual’s post-treatment movement function using a personalized NMS model of the individual (henceforth called “treatment optimization”), they will need to learn a combination of tools and write code that interfaces the tools with each other. Most tools mentioned thus far have well-developed introductory tutorials that teach new users about basic and some advanced functionality (51–56). However, most of these tutorials are not “real world” but rather work through smaller problems designed to teach specific isolated functionality. The tutorials also never work through the entire clinical treatment design process starting with model personalization and ending with treatment optimization. Due to their reduced scope, tutorials typically fail to communicate the nuances of tool use, which are critical for determining if and how a particular tool could be used to address a particular research goal.

To learn how to apply these tools to real-life clinical problems, researchers need to study published research articles. Unfortunately, reproducing published research is often challenging. On the one hand, there has been significant community effort to share models and data through web-based repositories such as SimTK (57) or AddBiomechanics (58). On the other hand, published articles do not include all implementation details needed for readers to go from initial models and data to final models and results, thereby preventing other researchers from reproducing and expanding upon the work. This state of affairs is understandable given the challenges of recording all steps and implementation details so that other researchers can reproduce them. This situation has resulted in a “reproducibility crisis” that significantly hinders progress of the field (59). If new researchers want to learn how to use existing software tools to create personalized predictive simulations that can be used for clinical treatment design, they must be resourceful, tenacious, and technically adept to assemble the necessary knowledge.

A major reason for this lack of “real-life” training materials for model personalization and treatment optimization is that existing software tools do not cover the entire process, so at best, tool developers can create tutorials that cover only isolated areas. To create real-life training materials that teach users the entire model personalization and treatment optimization process, we need a toolset that covers the full spectrum of these two processes. This problem has been addressed by the recently published Neuromusculoskeletal Modeling (NMSM) Pipeline (60). The NMSM Pipeline is a MATLAB-based open-source software package with integrated Model Personalization and Treatment Optimization toolsets for OpenSim (44,45). All tools within each toolset were designed to be used sequentially, with Model Personalization tools naturally interfacing with Treatment Optimization tools with minimal effort from the user. The inputs to the NMSM Pipeline are a scaled generic OpenSim model with or without personalized skeletal geometry (as appropriate) and individual subject movement data. Users can then personalize relevant joint structure, muscle-tendon, neural control, and foot-ground contact (as appropriate) model components for an individual subject and then use the subject’s personalized model in the treatment optimization process. Users can perform the entire model personalization and treatment optimization process without writing a single line of code. Instead, they create all necessary tasks using Extensible Markup Language (XML) settings files, created either in an OpenSim graphical user interface or by manual editing of XML files, which significantly lowers the barrier to entry for these high-end NMS modeling and simulation capabilities. This low barrier to entry can significantly increase the number of potential users who can explore computational treatment design using personalized NMS models.

Although the NMSM Pipeline makes treatment design with predictive simulations easier than it was previously, the tools still require significant training to learn. Basic tutorials for the NMSM Pipeline have already been made available (61), but like other musculoskeletal modeling software packages, these tutorials do not capture the nuances and details that should be understood to use the tools effectively for addressing real-life clinical problems. There is still a need for detailed tutorials that work through the entire model personalization to treatment optimization process for real-life clinical problems. Because the NMSM Pipeline makes model personalization and treatment optimization accessible and connected by a single workflow, it is now possible to design such in-depth tutorials. These tutorials can focus not only on what users are doing and why they are doing it, but also on how the final results are obtained.

This paper presents in-depth training tutorials for the NMSM Pipeline that guide users through real-life research to design treatments for two clinical conditions that impair walking function. These tutorials are the first to take users through the entire computational treatment design process, starting with model personalization and ending with treatment optimization using the personalized model. The first tutorial explores personalized joint and foot-ground contact modeling for a single subject with bilateral medial knee osteoarthritis (OA). The tutorial guides users through the model personalization and treatment optimization processes without muscles or neural control to design either a walking modification or high tibial osteotomy (HTO) surgery to reduce the peak adduction moment in both knees. The second tutorial explores personalized muscle-tendon and neural control modeling for a single subject with stroke. The tutorial guides users through the model personalization and treatment optimization processes including muscles and neural control to design a synergy-based functional electrical stimulation (FES) prescription to correct gait propulsion asymmetries. The two tutorials serve as complementary guides to using the NMSM Pipeline for research and give extra focus to the minute details that often affect the quality of predictive simulation results. The tutorials were deployed as two projects in a combined undergraduate/graduate course to demonstrate their effectiveness at teaching personalized neuromusculoskeletal modeling and treatment optimization to novice users. The NMSM Pipeline has potential applicability to a subset of clinical problems and patients, and ideally these tutorials will help researchers decide if the NMSM Pipeline might be applicable to their research problems.

Both tutorial documents are available as supplemental materials to this article (see Additional File 1.pdf and Additional File 2.pdf) and are also available at https://nmsm.rice.edu. Readers are encouraged to download the tutorial documents and read them in parallel with this article.

## Methods

### 1. Tutorial Overviews

The tutorials presented in this paper walk users through the model personalization and treatment optimization processes to design personalized treatments for two clinical conditions using the NMSM Pipeline and OpenSim (44,45). Each tutorial focuses on designing a clinical treatment for a single subject with gait dysfunction. The first tutorial explores skeletal modeling for one subject with bilateral medial knee osteoarthritis (OA) (male, age 47 years, height 1.7 m, mass 72.8 kg), while the second tutorial explores neuromusculoskeletal modeling for one subject who suffered a stroke (male, age 79 years, height 1.7 m, mass 80.5 kg; right-sided hemiparesis, LE Fugl-Meyer Motor Assessment 32/34 pts).

The experimental walking data used to develop both tutorials were collected from the two subjects under similar conditions. Ground reaction data were collected using a split-belt instrumented treadmill (Bertec Corp., Columbus, OH, USA) while the subjects walked at a self-selected speed (1.2 m/s for the subject with knee OA) or fastest comfortable speed (0.8 m/s for the subject with a stroke). Surface marker motion data were collected using a video-based motion capture system (Vicon Corp., Oxford, UK) with a full-body marker set that included three markers on the pelvis and each thigh, shank, and foot. In addition, for the subject who had a stroke, 16 channels of surface and fine wire EMG data were collected from each leg using two EMG systems (Motion Lab Systems, Baton Rouge, LA, USA). Informed consent was obtained from both subjects, and experimental data collection with subsequent modeling was approved by the institutional review board of the University of Florida. Gait data collected from these two subjects has been described in previous publications (23,60,62,63)

#### a. Clinical Problems

The first tutorial involves designing a walking modification or HTO surgical plan to treat an individual with bilateral medial knee OA. The peak knee adduction moment (KAM) during stance phase has been proposed as a surrogate measure for medial knee contact force, with a value less than 2.5% bodyweight x height (BWxHT) corresponding to the best long-term outcome for individuals with medial knee OA (64–66). For this subject, the target value for the peak KAM is 30.3 Nm. The peak KAM is derived entirely from external forces and skeletal kinematics and thus is a convenient measure for physics-based predictive simulations that do not calculate internal knee contact forces.

Two potential treatments for medial knee OA that can reduce the peak KAM are high tibial osteotomy (HTO) surgery and medial thrust gait (MTG) modifications (64,66). For the first treatment, an optimal HTO surgery is designed. This surgery involves cutting the subject’s proximal tibia and angling the distal end outward to turn a varus (bow-legged) knee into a valgus (knock-kneed) knee. The surgery is approximated in the OpenSim model by changing the knee adduction angle without altering the knee functional axis. The effect of the surgery on the resulting knee adduction moment during walking is estimated by performing a predictive simulation with the altered OpenSim model. The cost function minimizes changes relative to a baseline in the lower body joint moments controlling the motion. For the second treatment, optimal MTG modifications similar to (21) are designed. These gait modifications involve alterations to pelvic obliquity and lower body joint angles such that the knee moves closer to the midline of the body. No changes to the subject’s OpenSim model are required. The effect of the gait modifications on the resulting knee adduction moment during walking is estimated by performing a predictive simulation with the original OpenSim model. The cost function minimizes the knee adduction moment and changes relative to a baseline in the lower body joint moments controlling the motion, while path constraints maintain the same foot motion with respect to ground. For both potential treatments, all unlocked coordinates in the OpenSim model are treated as states that are changeable by the predictive simulations, and the treatments are designed for both knees since the subject has bilateral knee OA.

The second tutorial involves designing a synergy-based FES prescription to treat a person with right-sided hemiparesis caused by a stroke. The subject is high functioning but has an asymmetry in his propulsive and braking impulses between his legs, which leads to an inefficient gait pattern. Synergy-based FES is a novel treatment for stroke that electrically stimulates muscles based on the subject’s neural control characteristics (67–69). Muscle synergies can be calculated that decompose a subject’s muscle EMG data into a lower dimensional set of time-varying synergy activations (representing activation timing) and time-invariant synergy vector weights (representing activation coordination) (70–73). By comparing a subject’s synergy activations and vectors with those obtained from healthy subjects, researchers have identified a subject’s “impaired” synergies and then stimulated the subject’s paretic leg muscles with patterns consistent with corresponding healthy subject synergies (67,68).

For the subject in this tutorial, the calculated muscle synergies shared synergy vectors between the two legs while also making the subject’s personalized model walk the same way the subject did. Calculating bilateral muscle synergies in this unique way allowed direct comparison between the subject’s right and left leg synergy activations. This comparison revealed asymmetries in the subject’s synergy activation amplitudes but not shapes, allowing corresponding paretic/non-paretic pairs of synergy activations to be placed into one of three categories based on relative amplitudes: healthy (paretic ≈ non-paretic), impaired-compensatory (paretic/impaired < non-paretic/compensatory), and compensatory (paretic > non-paretic). We hypothesize that correcting the asymmetry in synergy activations will also correct the asymmetry in propulsive and braking impulses. The second tutorial will model a synergy-based FES protocol by scaling paired synergy activations so that they have equal amplitudes between legs. Cost terms will be used to adjust the propulsive and braking impulses so that these quantities are symmetrical as well.

#### b. Tool Descriptions

The primary software tools used in the tutorials are the NMSM Pipeline’s Model Personalization and Treatment Optimization toolsets (60). The NMSM Pipeline is MATLAB-based software that uses OpenSim models through OpenSim’s MATLAB application programming interface (API). The tutorials use software that is either open-source, widely available in an academic setting, or proprietary but inexpensive. MATLAB’s optimization toolbox, parallel computing toolbox, statistics and machine learning toolbox, curve fitting toolbox, symbolic math toolbox, and signal processing toolbox are required to use the NMSM Pipeline. The NMSM Pipeline’s Treatment Optimization toolset requires the proprietary GPOPS-II direct collocation optimal control solver (74). GPOPS-II currently costs $165 per year for a single user academic license and $825 per year for a department-wide academic license. The tutorials and results presented in this manuscript were obtained using the NMSM Pipeline Version 1.4.0, OpenSim Version 4.5, MATLAB Version 2024a, and GPOPS-II Version 2.5.

The NMSM Pipeline’s Model Personalization toolset consists of four tools: Joint Model Personalization, Ground Contact Model Personalization, Muscle-tendon Model Personalization, and Neural Control Model Personalization. All Model Personalization tools personalize the model by adjusting relevant parameters to track experimental data. Joint Model Personalization finds subject-specific joint parameter values (joint positions/orientations in body segments, body scale factors, marker locations on body segments) that allow the personalized model to reproduce experimental marker motions as closely as possible. Joint Model Personalization is important for obtaining accurate inverse dynamics joint loads, which can have large down-stream effects for predictive simulations (75–79). Ground Contact Model Personalization finds subject-specific foot-ground contact model parameter values (stiffness, damping, and friction parameters) that allow the personalized model to reproduce experimental ground reaction forces *and* moments as closely as possible. A personalized contact model is important to accurately predict ground reactions when a predictive simulation significantly changes foot kinematics (80). Muscle-tendon Model Personalization finds subject-specific EMG-driven model parameter values (EMG scale factors, activation dynamics parameters, Hill-type muscle-tendon model parameters) that allow the personalized model to reproduce experimental inverse dynamics joint moments as closely as possible (3,22,23). It can also estimate missing EMG signals using synergy extrapolation (63). Muscle-tendon Model Personalization has been shown to yield accurate amplitude and shape estimates for muscle activations, yielding more accurate muscle force estimates (22,81–87). Neural Control Model Personalization finds subject-specific muscle synergy controls that simultaneously reproduce muscle activations from Muscle-tendon Model Personalization and experimental inverse dynamics joint moments as closely as possible (88). Neural Control Model Personalization fits “functional” synergies that are able to reproduce the desired joint moments, which cannot be done by matching only muscle activations (33,88,89).

The NMSM Pipeline’s Treatment Optimization toolset consists of three tools: Tracking Optimization, Verification Optimization, and Design Optimization. Tracking Optimization generates a “starting point” dynamic simulation that closely reproduces all pre-treatment movement data available from the subject and is dynamically consistent. Achieving a dynamically consistent solution is an important step towards ensuring that the simulated treatment is physically realistic for the subject. Verification Optimization generates a “sanity check” predictive simulation that accurately reproduces the motion and external forces found by Tracking Optimization without tracking them explicitly, where the cost function closely tracks the controls found by Tracking Optimization. Verification Optimization is an important troubleshooting step because it isolates issues with optimal control problem formulation that may hinder Design Optimization. Design Optimization generates a predictive simulation of the patient’s post-treatment movement function when either a) a pre-defined treatment design is to be evaluated based on how well it achieves a specified treatment goal, or b) an optimal treatment design is to be found that achieves a specified treatment goal. Design Optimization adjusts the model controls and states produced by Verification Optimization and is the final step in the computational treatment design process. For more detail regarding NMSM Pipeline tools, readers are encouraged to read “The Neuromusculoskeletal Modeling Pipeline: MATLAB-based model personalization and treatment optimization functionality for OpenSim” by Hammond *et al.* (60).

### 2. Tutorial Descriptions

The knee OA tutorial focuses on skeletal modeling, and the stroke tutorial introduces muscles and neural control into the models. Each tutorial has four modules that focus on a different tool in the NMSM Pipeline and includes instructions on the relevant OpenSim tools (Table 1). The first two modules of each tutorial focus on the Model Personalization toolset, and the last two modules focus on the Treatment Optimization toolset. OpenSim tools are used throughout the tutorials as relevant to the tutorial material. Each module is designed to take approximately one week to complete, and so some steps are pre-done for users in the tutorial distribution to lower workload and focus on the primary tasks of each tutorial. Detailed instructions are given that are designed to “hold the user’s hand” through the entire modeling process to facilitate active learning and focus on the primary learning objectives of each tutorial. At the end of each module, users are prompted to gather deliverables that are designed to guide them through critical evaluation of their results. These deliverables include presentation of raw results, analysis and interpretation of the results, and explanation of the tools being used.

**Table 1:** List of NMSM Pipeline and OpenSim tools that are used in each tutorial. For tools labeled Performed, users specifically work through that tool in the tutorial. For tools labeled Provided, pre-generated results are provided to users.

| Toolset | Tool | Tutorial 1 | Tutorial 2 |
| --- | --- | --- | --- |
| NMSM Pipeline<br>Model<br>Personalization | Joint Model Personalization | Performed | Provided |
|  | Ground Contact Model Personalization | Performed | Provided |
|  | Muscle-tendon Model Personalization | N/A | Performed |
|  | Neural Control Model Personalization | N/A | Performed |
| NMSM Pipeline<br>Treatment<br>Optimization | Tracking Optimization | Performed | Performed |
|  | Verification Optimization | Performed | Performed |
|  | Design Optimization | Performed | Performed |
| OpenSim | Scale Model | Performed | Provided |
|  | Inverse Kinematics | Performed | Provided |
|  | Inverse Dynamics | Performed | Provided |
|  | Muscle Analysis | N/A | Performed |

#### a. Tutorial 1 Layout

The first tutorial guides users through using the NMSM Pipeline computational treatment design process for a situation where modeling of muscles and neural control may not be needed and skeletal modeling is sufficient. The tutorial uses the NMSM Pipeline’s Joint Model Personalization, Ground Contact Model Personalization, and torque-driven Treatment Optimization tools, as well as OpenSim’s Scale Model and Inverse Kinematics tools to design a walking modification or HTO surgery for a single subject with bilateral knee OA. The tutorial consists of four modules, with two Model Personalization modules and two Treatment Optimization modules. The inputs and outputs of all modules are presented in Figure 1. The tutorial document can be found in Additional File 1.pdf (Supplementary Material A - Tutorial 1).

**Figure 1:**
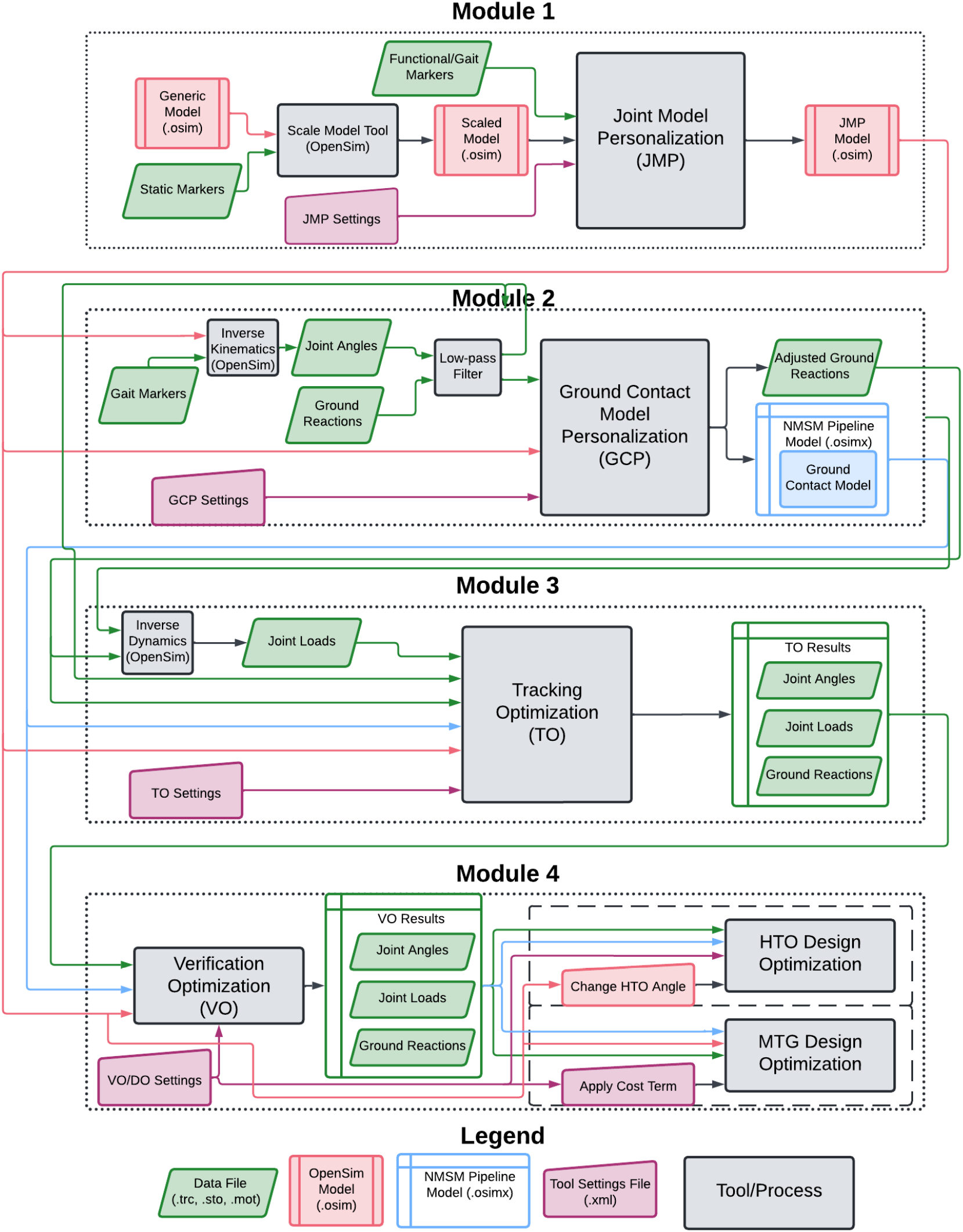
Flowchart showing the inputs and outputs of each module in Tutorial 1. Green elements indicate data files, red elements indicate OpenSim models, blue elements indicate NMSM Pipeline models, and purple elements indicate NMSM Pipeline settings files.

The first module teaches users how to perform the Joint Model Personalization process. This process involves using the NMSM Pipeline’s Joint Model Personalization tool, and OpenSim’s Scale Model and Inverse Kinematics tools. The first task of the module guides users through a unique model scaling process that ensures that the foot markers are placed correctly on the feet to help with the subsequent Ground Contact Model Personalization runs. The second task of the module guides users through using the Joint Model Personalization tool to personalize lower extremity functional axes both sequentially in multiple optimizations (i.e., personalize one joint at a time) and simultaneously in one optimization (i.e., personalize all joints at once). The deliverables for this task include an analysis of inverse kinematics marker errors for both Joint Model Personalization problem formulations and an analysis of which Joint Model Personalization problem formulation is optimal for this subject.

The second module teaches users how to perform the Ground Contact Model Personalization process. This process involves calibrating the subject’s foot-ground contact model using the NMSM Pipeline’s Ground Contact Model Personalization tool and OpenSim’s Inverse Kinematics tool. The first task of the module guides users through using the Inverse Kinematics tool to calculate joint angles, and then the joint angles are filtered in preparation for Ground Contact Model Personalization. The second task of the module guides users through using the Ground Contact Model Personalization tool to create a personalized foot-ground contact model that closely reproduces experimental ground reaction forces and moments. Users explore two different friction models for the foot-ground contact model: Coulomb friction and viscous friction. The deliverables for this module include an analysis of the experimental data tracking errors for both ground contact models, and a brief explanation of which friction model worked best for this subject.

The third module teaches users how to perform the Tracking Optimization process with joint torque controls. This process involves creating dynamically consistent joint torque-driven walking predictions using the NMSM Pipeline’s Tracking Optimization tool and OpenSim’s Inverse Dynamics tool. The first task of the module guides users through using the Inverse Dynamics tool to calculate joint loads with their previously calculated and filtered Inverse Kinematics joint angles and experimental ground reaction data. The second task of the module guides users through predicting a dynamically consistent torque-driven walking motion using the Tracking Optimization tool. The deliverables for this module include an analysis of the tracking quality for all experimental quantities included in the Tracking Optimization run, and a brief explanation of which quantities have the worst error and how potentially to improve that error.

The fourth module teaches users how to design treatments for walking impairments using the Design Optimization tool with joint torque controls. This process involves using the NMSM Pipeline’s Verification Optimization and Design Optimization tools to design either a new MTG walking motion or an HTO surgical intervention that achieves a target value for the subject’s peak KAM in both knees. The first task of the module guides users through using the Verification Optimization tool to verify that the joint torque controls found by Tracking Optimization are able to produce the same motion, joint loads, and ground reactions without tracking these quantities explicitly. The second task of the module guides users through using the Design Optimization tool to design a treatment that achieves a target value for the subject’s peak KAM.

This module gives users freedom to design either an HTO surgery or an MTG motion, or both, with the goal of reducing the subject’s peak KAM to the target value of 2.5% BWxHT for both knees. The design element for the HTO Design Optimization is a fixed offset for the knee adduction angle in each knee of the model. No extra cost or constraint terms are added for this problem. The design element for the MTG Design Optimization is a knee adduction moment minimization cost term for both legs that penalizes moment values greater than a specified value (termed “max allowable error” for all NMSM Pipeline cost terms). For this problem, cost tracking and constraint deviation terms are added for foot marker position, foot orientation, and torso orientation to better replicate the results of (21). These cost and constraint terms are added in Verification Optimization without any design elements to verify that they are consistent with the overall problem formulation. For both Design Optimizations, the value of the design element is iterated over three Design Optimization runs to find the value for each knee that achieves the target peak KAM of 2.5% BWxHT. The design variables for both optimizations are the joint positions, velocities, and accelerations for unlocked coordinates in the OpenSim model along with all lower body joint torque controls, while the cost function for both optimizations also minimizes changes in lower body joint torque controls and upper body joint angles to predict a dynamically consistent walking motion. The deliverables for this module include an analysis of the peak KAM produced by the designed treatment and a brief discussion on the limitations of the predictive simulation to achieve the desired clinical goal.

#### b. Tutorial 2 Layout

The second tutorial guides users through using the NMSM Pipeline computational treatment design process for a situation where modeling of muscles and neural control is clearly needed. The tutorial uses the NMSM Pipeline’s Muscle-tendon Model Personalization, Neural Control Model Personalization, and muscle synergy-driven Treatment Optimization tools and OpenSim’s Muscle Analysis tool to design a synergy-based FES prescription for a single subject with stroke. The tutorial also consists of four modules, with two Model Personalization modules and two Treatment Optimization modules. The tools used in this tutorial generally take longer to run compared to the first tutorial, and so for some modules, users work through smaller runs on the tools and are given pre-generated results for bigger problems to facilitate data analysis. The inputs and outputs of all modules are presented in Figure 2. The tutorial document can be found in Additional File 2.pdf (Supplementary Material B - Tutorial 2).

**Figure 2:**
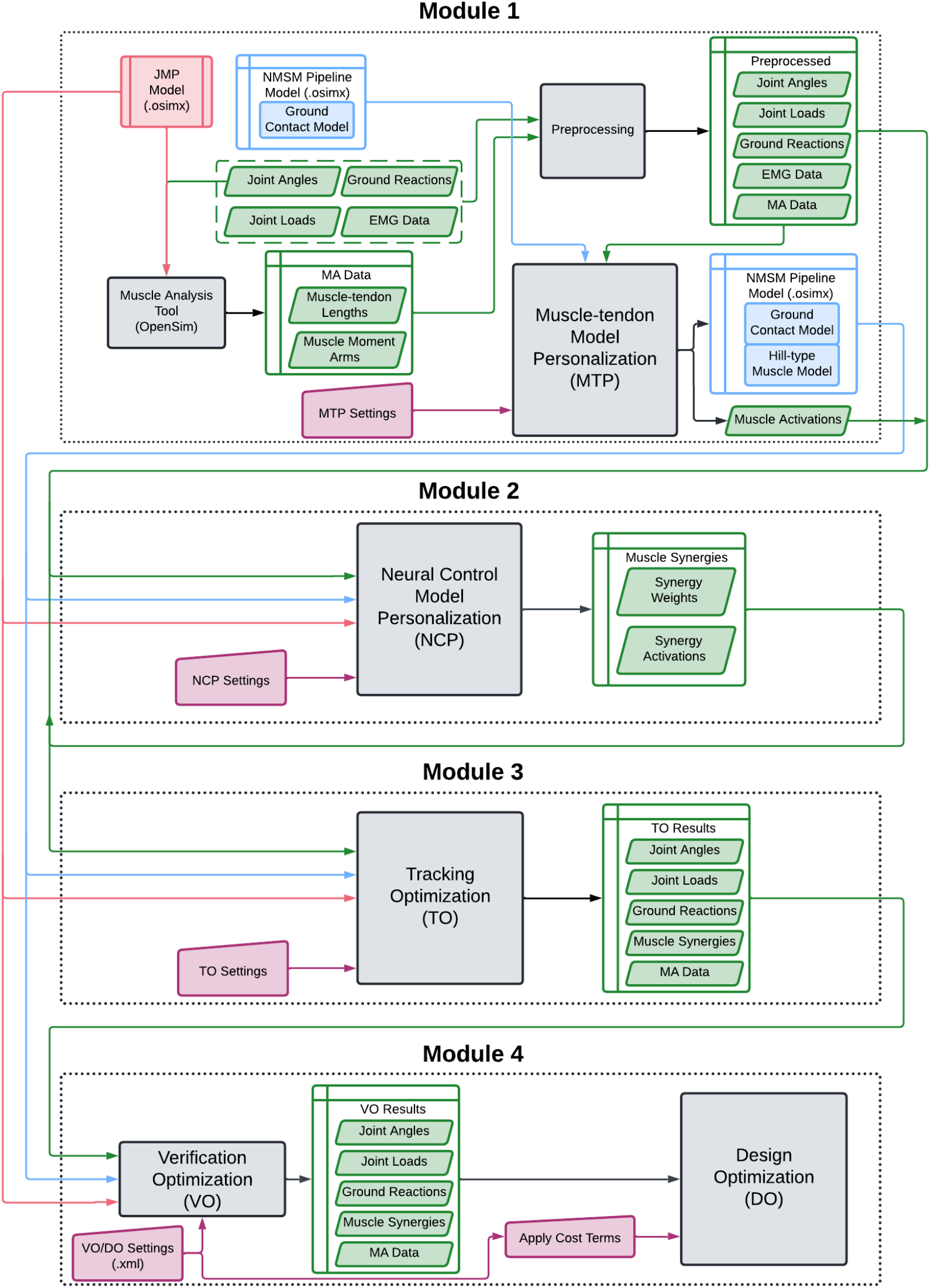
Flowchart showing the inputs and outputs of each module in Tutorial 2. The legend can be found in Figure 1. Green elements indicate data files, red elements indicate OpenSim models, blue elements indicate NMSM Pipeline models, and purple elements indicate NMSM Pipeline settings files. Joint angles, joint loads, ground reactions, EMG data, and a post-JMP model are provided at the beginning of module 1.

The first module teaches users how to perform the Muscle-tendon Model Personalization process. This process involves predicting muscle forces for the subject using the NMSM Pipeline’s Muscle-tendon Model Personalization tool and Data Preprocessing utility and OpenSim’s Muscle Analysis tool. The first task of this module focuses on using OpenSim’s Muscle Analysis tool to calculate muscle-tendon lengths and moment arms that will be used by Muscle-tendon Model Personalization. The second task of this module has users employ the NMSM Pipeline’s Data Preprocessing utility to crop, filter, and resample data in preparation for Muscle-tendon Model Personalization. The third task of this module has users use the NMSM Pipeline’s Muscle-tendon Model Personalization tool to personalize muscle-tendon parameters to best predict muscle forces, guiding them through an iterative process to obtain the best Muscle-tendon Model Personalization result. The deliverables for this module include a brief explanation of how the Muscle-tendon Model Personalization tool works, an analysis of the four iterations students perform, and a brief discussion of why Muscle-tendon Model Personalization is important for muscle-driven predictive simulations.

The second module teaches users how to perform the Neural Control Model Personalization process. This process involves creating and analyzing a muscle synergy neural control model for the subject using the NMSM Pipeline’s Neural Control Model Personalization tool. The first task of this module briefly shows users how to format their Neural Control Model Personalization input data directory after running Muscle-tendon Model Personalization. The second task of this module guides users through choosing the appropriate number of muscle synergies to use in their Neural Control Model Personalization run by analyzing the variance accounted for (VAF) of the EMG data as the number of synergies increases. The third task for this module guides users through creating and running an initial Neural Control Model Personalization problem. After users run their Neural Control Model Personalization problem, they are given premade results for data analysis with five synergies and bilaterally symmetric synergy vectors between legs. The deliverables for this module are extensive and guide users through the full data analysis and diagnosis of the subject’s neural impairment. Users are instructed to analyze the shape of the synergy activations when bilaterally symmetric synergy vectors are specified, identify which synergies are healthy, impaired, or compensatory, and identify the function of each of the synergies. After completing the deliverables, users should have a strong grasp on how to use Neural Control Model Personalization to gain insight into a subject’s neural control impairment.

The third module teaches users how to perform the Tracking Optimization process with muscle-synergy controls. This process involves creating a dynamically consistent muscle synergy-driven walking motion using the NMSM Pipeline’s Tracking Optimization tool and Surrogate Muscle Model utility. The first task of this module guides students through creating a surrogate muscle model using the NMSM Pipeline and OpenSim’s Muscle Analysis tool. The second task of this module guides students through creating and running a synergy-driven Tracking Optimization settings file. Synergy-driven Tracking Optimizations are often difficult to get to converge to a good solution, and so extra attention is given toward analyzing the convergence of the optimization and how to troubleshoot potential issues. Because synergy-driven Tracking Optimization runs are computationally expensive, users were instructed to run only one Tracking Optimization problem, and premade results were used for further data analysis. The deliverables for this module guide users through analysis of their Tracking Optimization results and the premade Tracking Optimization results with a mix of quantitative analysis of experimental tracking results and qualitative analysis of the gait motion produced by Tracking Optimization.

The fourth module teaches users how to design treatments for walking impairments using the Design Optimization tool with muscle-synergy controls. This process involves using the NMSM Pipeline’s Verification Optimization and Design Optimization tools to design a synergy-based FES prescription to improve walking symmetry for a person with stroke. The first task of this module guides users through using the Verification Optimization tool to verify that the synergy controls produced by Tracking Optimization are able to produce the same motion, joint loads, muscle activations, and ground reactions without tracking these quantities explicitly. The second task of this module prompts users to plan their FES prescription. The goal for this task is to choose a set of scale factors to apply to the synergy activations such that the amplitude of each synergy activation is equal between legs. The third task of this module guides users through implementing their FES prescription in Design Optimization to equalize the propulsive and braking impulses between the subject’s feet. The target propulsive and braking impulses are calculated by averaging the experimental propulsive and braking impulses, and cost terms are used to achieve this target value. Additional cost terms are added that track a scaled version of the synergy activations such that their shapes stay nearly the same but their amplitudes can increase or decrease. It is assumed that users are stimulating only the impaired synergies and that the subject will be capable of self-correcting any compensatory synergies in response to the stimulation. Next, users analyze their Design Optimization results, explain their rationale for the scale factors they chose, and present the potential limitations of implementing their FES prescription to a clinical setting.

## Results

Simulation results for both tutorials are presented in this section. For simulations that tracked experimental quantities, root mean squared error (RMSE) values are reported. Iteration counts and computation times for each module are reported for runs performed using a Windows 11 PC workstation possessing an AMD Ryzen 9 7950x 16-core processor with 32 GB of DDR5 RAM.

### a. Tutorial 1 Results

The Joint Model Personalization module yielded substantially lower marker errors for both the sequential and simultaneous runs. Prior to Joint Model Personalization, Inverse Kinematics marker tracking errors during gait were an average of 0.92 cm with a maximum error of 2.1 cm. The sequential Joint Model Personalization run reduced the average marker tracking error for gait to 0.65 cm and the maximum error to 1.6 cm. The simultaneous Joint Model Personalization run reduced the average marker tracking error for gait to 0.57 cm and the maximum error to 1.6 cm (Figure 3). The maximum marker tracking errors for both the simultaneous and sequential runs were larger than the errors presented in previous studies using the same personalization method, but the average marker tracking errors were of similar magnitude to those studies (60,79). For an analysis of marker tracking errors produced by a scaled generic OpenSim model, refer to Additional File 3.pdf (Supplementary Material C - Scaled Generic Model Comparison). The simultaneous Joint Model Personalization run produced the lowest marker tracking errors and so was used for the remainder of the tutorial. This Joint Model Personalization run terminated after 6 iterations and 0.37 hours of wall clock time.

**Figure 3:**
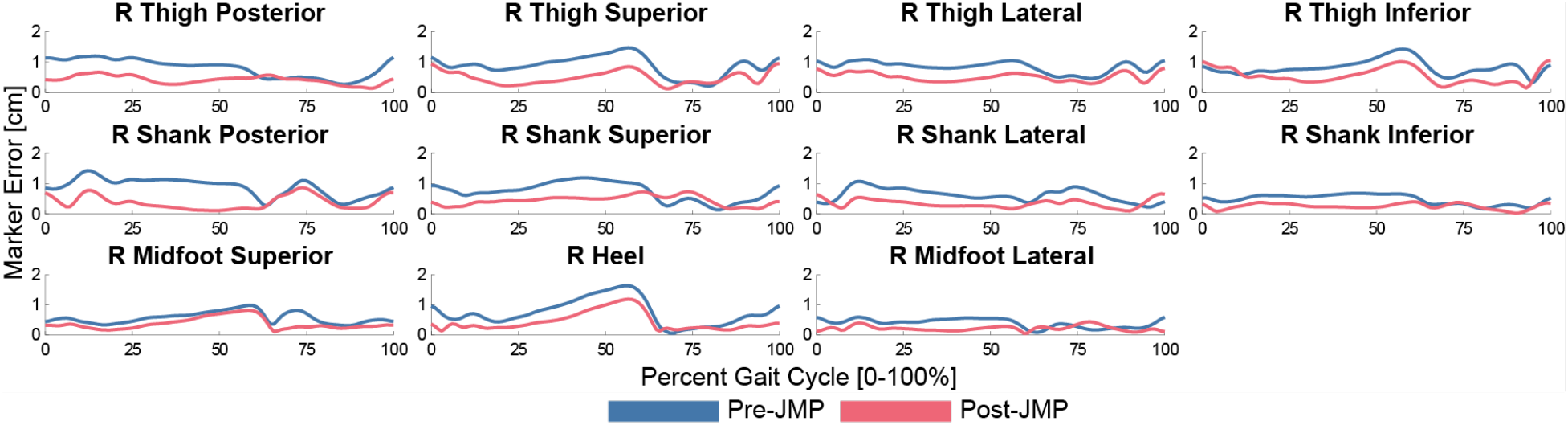
Right leg inverse kinematics marker tracking errors before Joint Model Personalization (blue) and after Joint Model Personalization (red). The x-axis represents percent gait cycle from right heel-strike to right heel-strike [0-100%], and the y-axis represents marker distance errors [cm]. Joint Model Personalization greatly reduced average and maximum marker tracking errors for all lower limb markers included in the optimization.

The Ground Contact Model Personalization module calibrated a foot-ground contact model for each foot that could accurately reproduce experimental ground reaction forces and moments with minimal changes in foot kinematics. The Ground Contact Model Personalization model with viscous friction achieved an average ground reaction force tracking RMSE of 5.9 N and an average ground reaction moment tracking RMSE of 0.81 Nm with average RMSEs for calcaneus orientation and translation tracking and toes angle tracking of 1.3 degrees, 0.38 cm, and 1.1 degrees, respectively. The Ground Contact Model Personalization model with Coulomb friction achieved an average ground reaction force tracking RMSE of 4.9 N and an average ground reaction moment tracking RMSE of 0.77 Nm with average RMSEs for calcaneus orientation and translation tracking and toes angle tracking of 1.2 degrees, 0.37 cm, and 1.1 degrees, respectively (Figure 4). All ground reaction and foot kinematic RMSE values were comparable to results published in a previous study that used a similar ground contact model (60) and better than results published in studies that used a different ground contact model (3). The Ground Contact Model Personalization model with Coulomb friction produced the lowest tracking errors and so was used for the subsequent Treatment Optimization. This Ground Contact Model Personalization run is terminated after 87 iterations and 0.19 hours of wall clock time.

**Figure 4:**
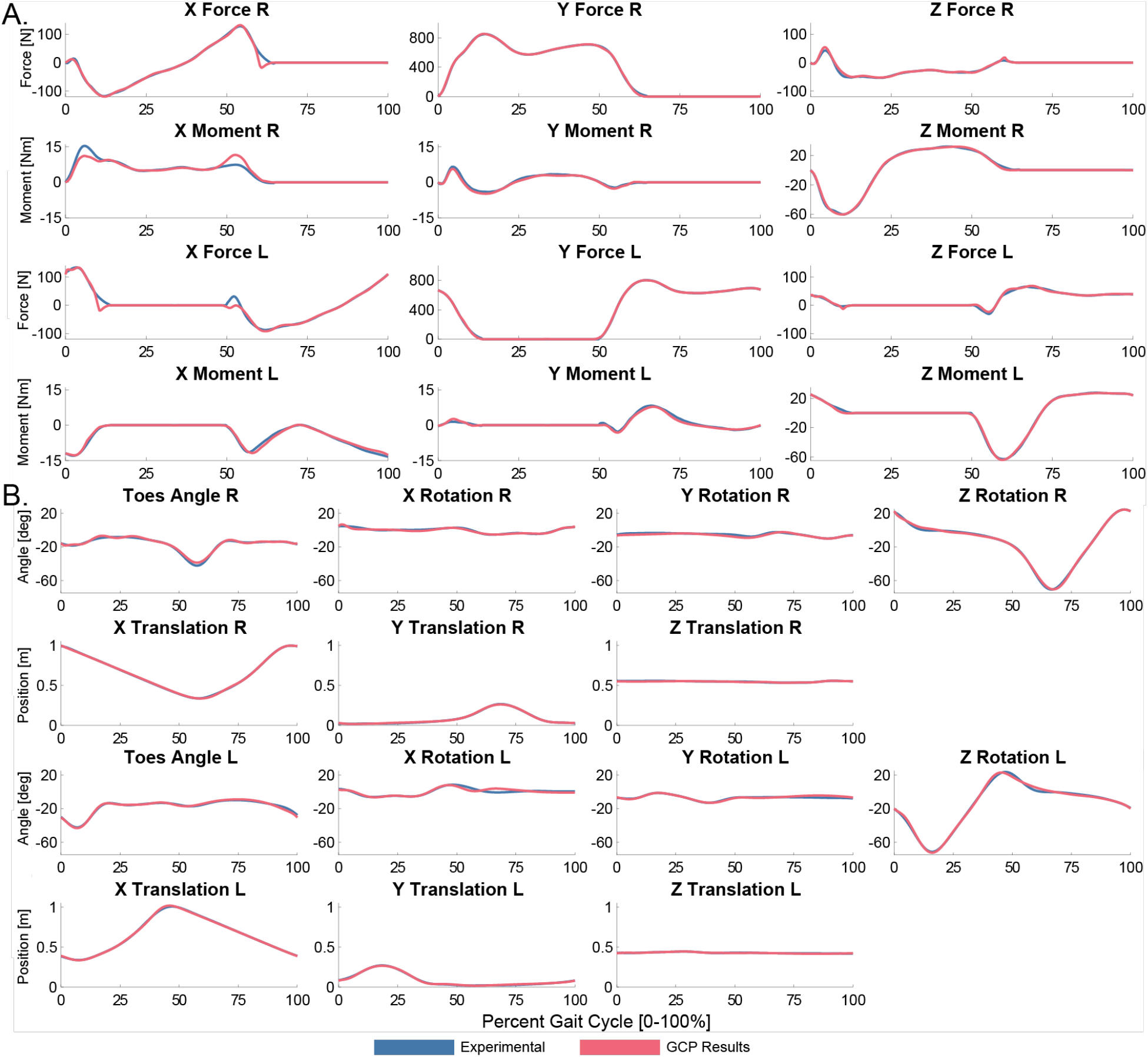
(A) Ground reaction forces and moments and (B) kinematic tracking quality from Ground Contact Model Personalization using Coulomb friction. The x-axis represents percent gait cycle from right heel-strike to right heel-strike [0-100%]. For subfigure A, the y-axis represents forces [N] or moments [Nm]. For subfigure B, the y-axis represents coordinate angle [deg] or foot translation [m]. Ground reactions produced by Ground Contact Model Personalization (red) closely tracked experimental ground reactions (blue) with minimal changes in experimental foot kinematics, indicating excellent Ground Contact Model Personalization results.

The Tracking Optimization module produced a dynamically consistent walking motion that closely tracked all joint angles, joint loads, and ground reactions. All joint rotations were tracked to within 1.8 degrees RMSE, and all joint translations were tracked to within 0.29 cm RMSE. All lower limb joint moments were tracked to within 2.0 Nm. (Figure 5) The peak KAM for the right and left legs was -44.3 Nm and -43.15 Nm, respectively (Figure 6). Ground reaction forces were all tracked to within 9 N RMSE and ground reaction moments to within 2.1 Nm RMSE. The torque-driven Tracking Optimization tracking results were of comparable quality to those produced in a similar three-dimensional predictive simulation study that tracked actual subject experimental data (33). This Tracking Optimization run converges in 95 iterations and 0.19 hours of wall clock time.

**Figure 5:**
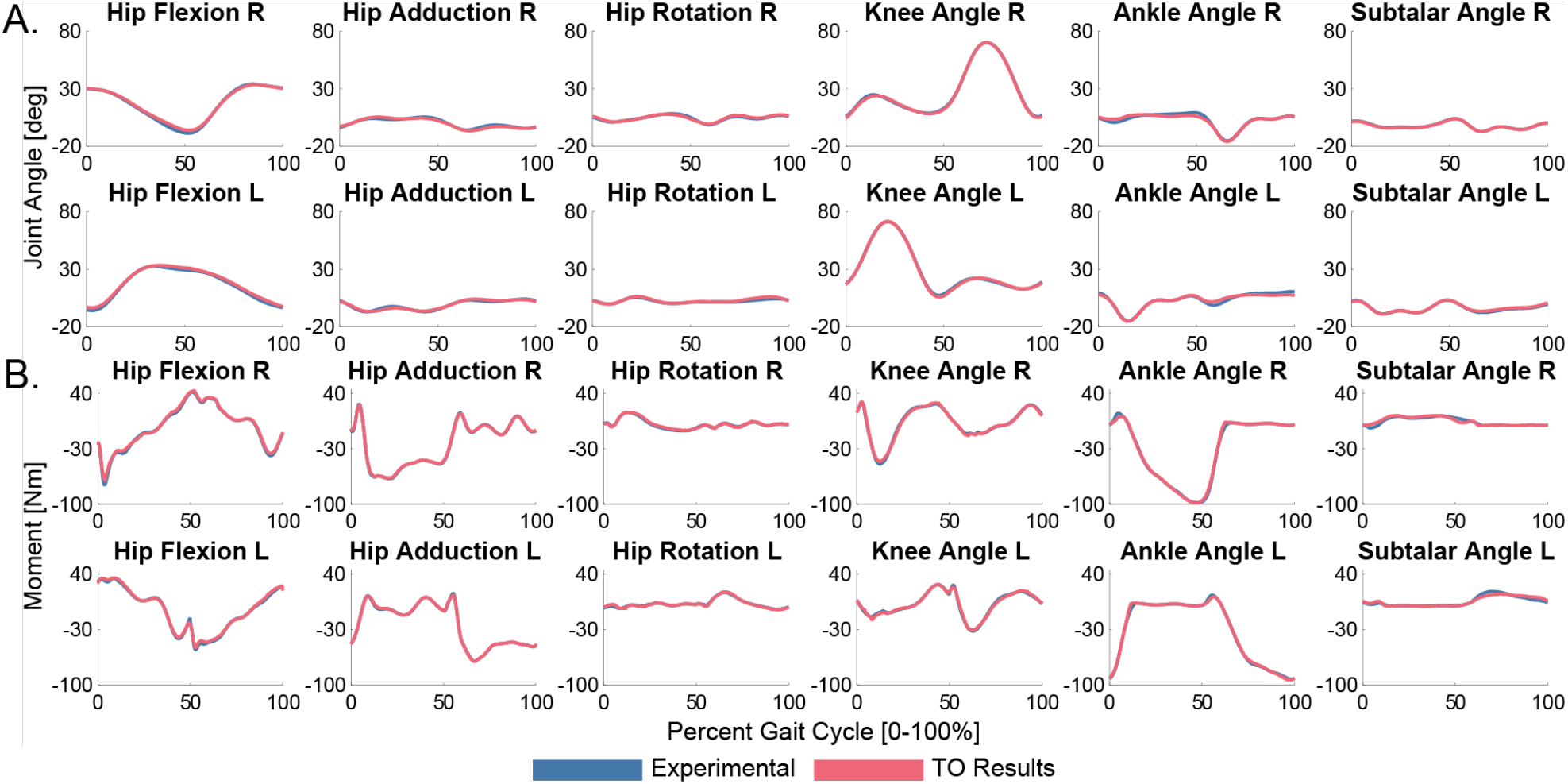
Torque-driven Tracking Optimization (TO) results for (A) joint angles and (B) joint moments. The x-axis represents percent gait cycle from right heel-strike to right heel-strike [0-100%]. For subfigure A, the y-axis represents joint angles [deg]. For subfigure B, the y-axis represents joint moments [Nm]. The Tracking Optimization results (red) closely reproduced all lower limb experimental joint angles and moments (blue) while achieving dynamic consistency.

**Figure 6:**
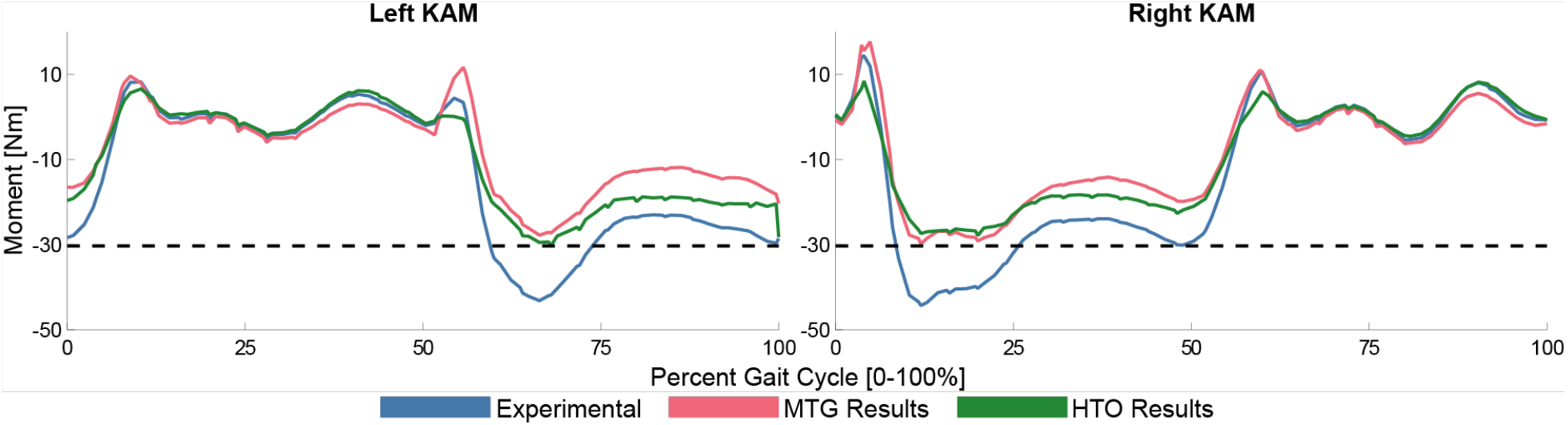
Left and right knee adduction moment (KAM) plots for the experimental (blue), high tibial osteotomy (HTO) (red), and medial thrust gait (MTG) (green) motions. The dashed black line represents the target value for the magnitude of the peak knee adduction moment, which was 30.3 Nm. The x-axis represents percent gait cycle from right heel-strike to right heel-strike [0-100%]. The y-axis represents the knee adduction moment [Nm]. Both the HTO and MTG predictive simulations produced a peak KAM at or slightly below the target value.

The Verification Optimization results closely tracked the Tracking Optimization solution with negligible RMSE across all quantities, thus verifying that the Tracking Optimization problem was well-formulated and ready for Design Optimization. The HTO Verification Optimization run converges in 3 iterations and 0.02 hours of wall clock time and the MTG Verification Optimization run converges in 4 iterations and 0.02 hours of wall clock time.

The HTO and MTG Design Optimization runs were both able to reduce the peak KAM to below the desired value of 30.3 Nm (Figure 6). For the HTO problem, corrections of 3 degrees, 6 degrees, and 9 degrees were tested, and ∼6 degrees was found to be the optimal correction for this subject. For the MTG problem, maximum allowable errors of 10 Nm, 15 Nm, and 20 Nm for the KAM were tested, and ∼15 Nm was found to be the optimal value for this subject. The final HTO Design Optimization run converged in 60 iterations and 0.12 hours of wall clock time, while the final MTG Design Optimization run converged in 181 iterations and 0.38 hours of wall clock time.

### b. Tutorial 2 Results

The Muscle-tendon Model Personalization and Neural Control Model Personalization modules yielded accurate joint moment matching results for the final runs that were used in Tracking Optimization. The final Muscle-tendon Model Personalization problem formulation produced personalized muscle-tendon models that matched experimental joint moments to within 5.1 Nm RMSE with maximum absolute errors within 4.5 Nm (Figure 7). The Muscle-tendon Model Personalization moment tracking quality was at least as good as previous studies that used an EMG-driven modeling procedure to calibrate muscles spanning the hip, knee, and ankle simultaneously (22,60,63). The final Muscle-tendon Model Personalization run converged in 508 iterations and 0.08 hours of wall clock time.

**Figure 7:**
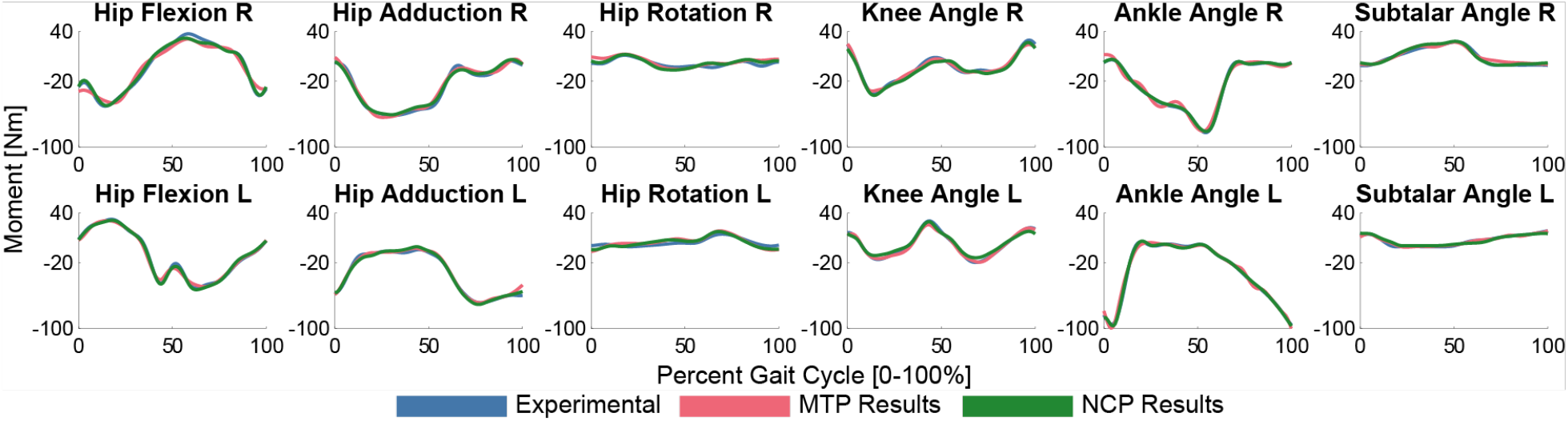
Muscle-tendon Model Personalization (MTP) (red) and Neural Control Model Personalization (NCP) (green) tracking quality for experimental joint moments (blue). The x-axis represents percent gait cycle from right heel-strike to right heel-strike [0-100%]. The y-axis represents joint moments [Nm]. Both Muscle-tendon Model Personalization and Neural Control Model Personalization closely reproduced the experimental joint moments, while Neural Control Model Personalization created a smoothing effect on the joint moments that produced better experimental moment tracking.

The Neural Control Model Personalization tool produced a muscle synergy model with identical synergy vectors for both legs and that matched joint moments to within 3.2 Nm RMSE and muscle activations from Muscle-tendon Model Personalization with a total percent VAF of 91% and a worst individual muscle RMSE of 0.092. The Neural Control Model Personalization moment tracking results were very similar to those reported in a similar study using this computational method (60). This Neural Control Model Personalization run converged in 641 iterations and 0.55 hours of wall clock time.

The Tracking Optimization module produced a dynamically consistent synergy-driven walking motion that closely tracked all joint angles, joint loads, ground reactions, and muscle activations. All lower limb joint rotations were tracked to within 3.2 degrees RMSE and all joint translations were tracked to within 0.57 cm RMSE. All lower limb joint loads were tracked to within 6.0 Nm (Figure 8). Horizontal ground reaction forces were all tracked to within 8.2 N RMSE, with vertical ground reaction forces being tracked to within 27 N RMSE, while all ground reaction moments were tracked to within 2.6 Nm RMSE. All muscle activations were tracked to within 0.075 RMSE. All of these quantities were similar to values reported in other published synergy-driven predictive simulation studies that used actual subject walking data (23,33,60). The synergy activations produced by Tracking Optimization and their corresponding synergy vectors are provided in Figures 9 and 10, respectively. The cosine similarities between the non-paretic and paretic leg synergy vectors were 0.99, 0.99, 0.97, 0.99, and 0.98 for synergies 1 through 5 respectively (Figure 10). This Tracking Optimization run converges in 270 iterations and 6.25 hours of wall clock time.

**Figure 8:**
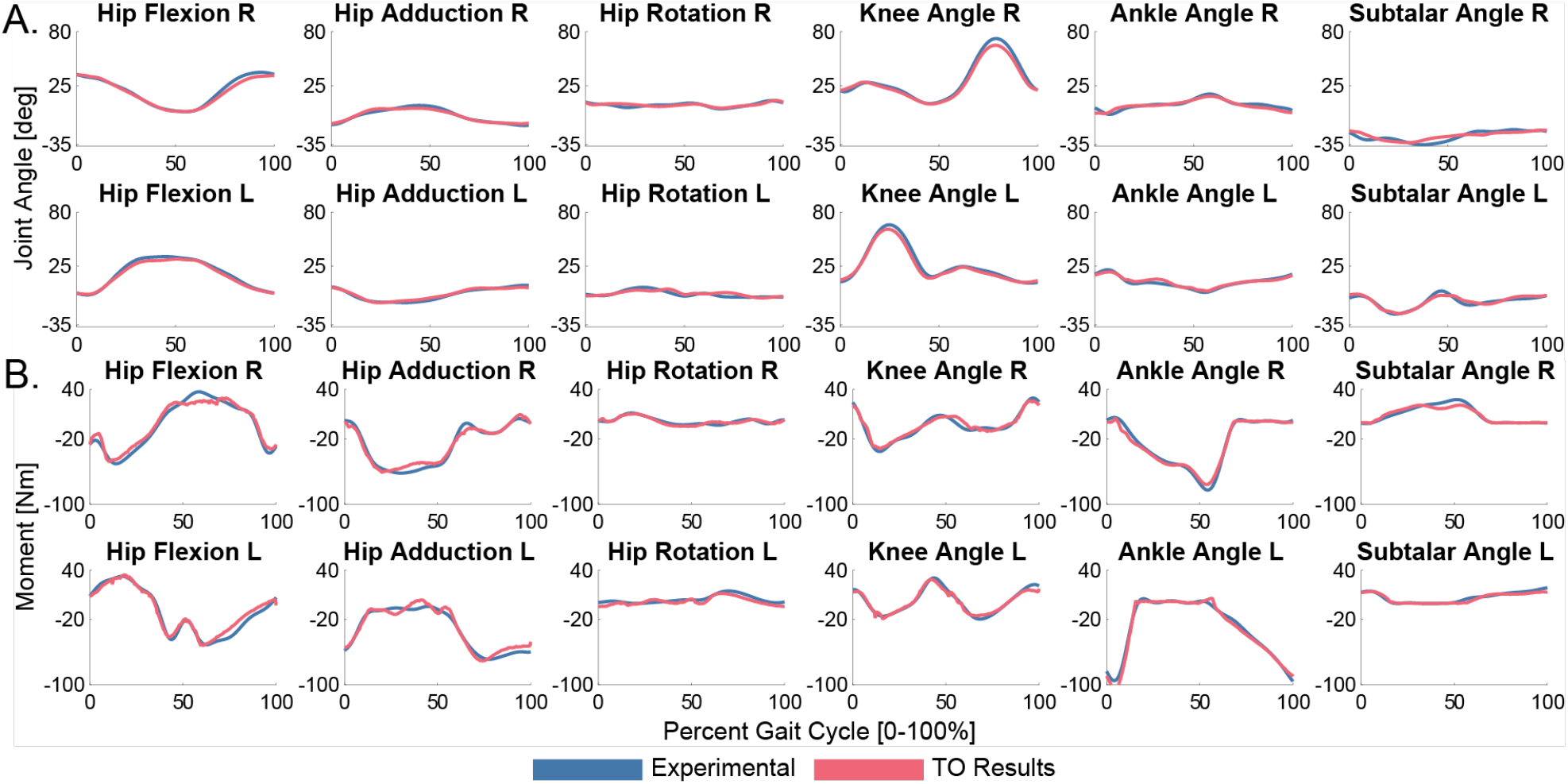
Synergy-driven Tracking Optimization results for (A) joint angles and (B) joint moments. The x-axis represents percent gait cycle from right heel-strike to right heel-strike [0-100%]. For subfigure A, the y-axis represents the joint angles [deg]. For subfigure B, the y-axis represents the joint moments [Nm]. The Tracking Optimization results (red) closely reproduced all lower limb experimental joint angles and moments (blue).

**Figure 9:**
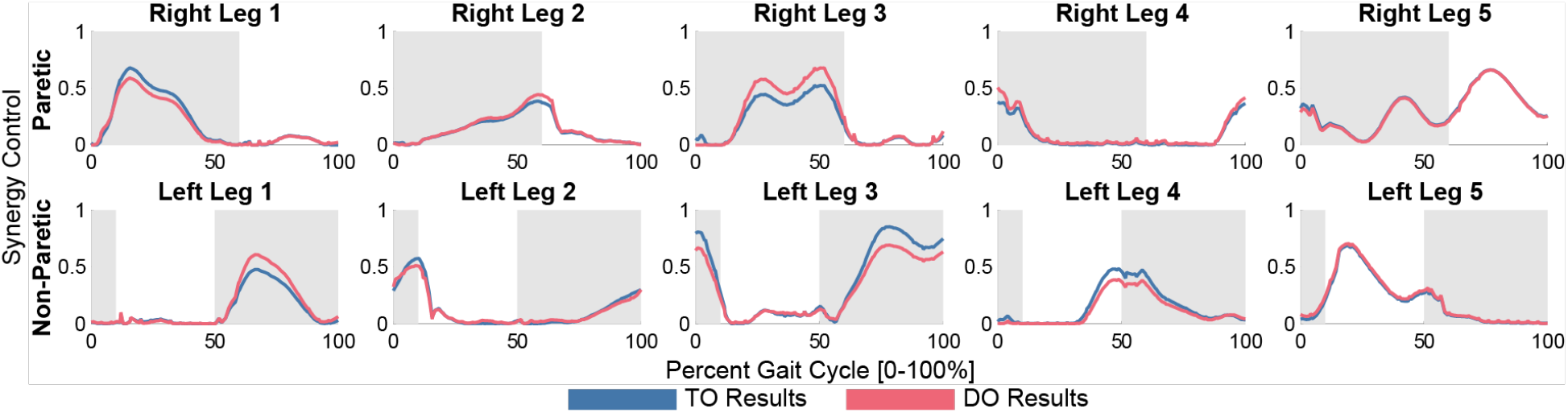
Tracking Optimization (TO) (blue) and Design Optimization (DO) (red) synergy activations throughout the gait cycle. The x-axis represents percent gait cycle from right heel-strike to right heel-strike [0-100%]. The y-axis represents synergy activation amplitude [unitless, 0-1]. The stance phases for each leg are shaded. The associated synergy vectors produced by Tracking Optimization are shown in Figure 10.

**Figure 10:**
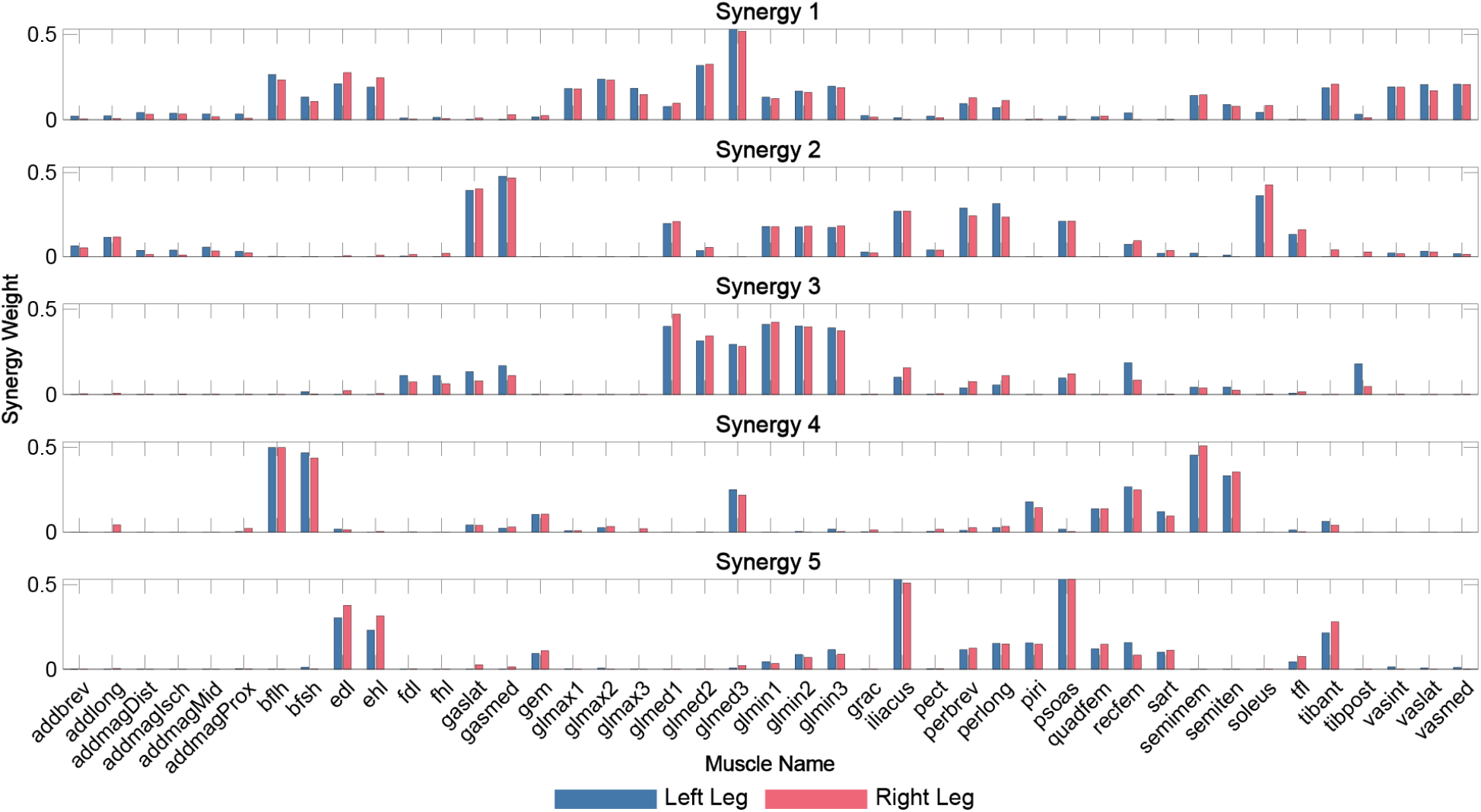
Left (non-paretic, blue) and right (paretic, red) synergy vectors produced by Tracking Optimization. The x-axis contains one muscle per pair of bars, and the y-axis represents the value of the synergy vectors (unitless). By design, the synergy vectors were nearly symmetric. Neural Control Model Personalization constrained the synergy vectors to be the same between the two legs, and Tracking Optimization was allowed to make small changes to them to facilitate achieving dynamic consistency. Synergy vectors were not changed in Design Optimization.

The Verification Optimization results closely tracked the Tracking Optimization solution with negligible RMSE across all quantities, thus verifying that the Tracking Optimization solution was valid and ready for Design Optimization. This Verification Optimization run converges in 55 iterations and 0.28 hours of wall clock time.

The synergy-driven Design Optimization run with cost terms for synergy activation scaling and propulsive and braking impulse targets successfully equalized the propulsive and braking impulses to within 5% error of each other (Table 2). The presented results used a synergy activation scaling strategy where the right and left leg synergy activations were scaled up or down to their average amplitude found by Tracking Optimization (Figure 9). Corresponding synergy vectors are shown in Figure 10 and were unchanged from Tracking Optimization. Symmetry indices for predicted propulsive and braking impulses were also calculated using the equation

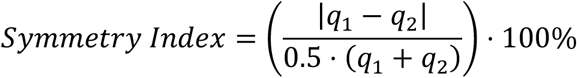

where *q*_1_ and *q*_2_ are discrete positive values used for quantifying symmetry. Symmetry indices are reported for propulsive and braking impulses between each leg to quantify the compensation required by the non-paretic leg, and for propulsive and braking impulses within each leg to quantify the relative strength of each leg. High asymmetry between legs but low asymmetry within legs would indicate a strength discrepancy between legs. Design Optimization improved all symmetry indices within each leg and between the two legs (Table 3) from an average of 100.3% (highly asymmetric) to 0.9% (almost perfectly symmetric). The Design Optimization run converged in 229 iterations and 1.10 hours of wall clock time.

**Table 2:** Propulsive and braking impulses after Tracking Optimization (TO) and after Design Optimization (DO). The Design Optimization problem could equalize the propulsive and braking impulses to within 0.08 Ns of the target value of 10.77 Ns.

|  |  | Propulsive Impulse [Ns] | Braking Impulse [Ns] |
| --- | --- | --- | --- |
| TO Results | Right Leg | 5.92 | 15.64 |
|  | Left Leg | 18.42 | 5.35 |
| DO Results | Right Leg | 10.77 | 10.80 |
|  | Left Leg | 10.84 | 10.69 |

**Table 3:** Symmetry indices for propulsive and braking impulses after Tracking Optimization (TO) and after Design Optimization (DO). Symmetry indices are reported for 1) propulsive impulses between the right and left leg, 2) braking impulses between the right and left leg, 3) propulsive and braking impulses on the right leg, and 4) propulsive and braking impulses on the left leg.

| Symmetry Pair |  | Symmetry Index (%) |
| --- | --- | --- |
| TO Results | Right-Left Propulsive | 102.7 |
|  | Right-Left Braking | 98.1 |
|  | Right Leg Propulsive-Braking | 90.2 |
|  | Left Leg Propulsive-Braking | 110.0 |
| DO Results | Right-Left Propulsive | 0.7 |
|  | Right-Left Braking | 1.0 |
|  | Right Leg Propulsive-Braking | 0.3 |
|  | Left Leg Propulsive-Braking | 1.4 |

## Discussion

This paper presented a pair of neuromusculoskeletal modeling tutorials using the NMSM Pipeline and OpenSim that provide extensive details on how to approach the computational treatment design process. Extra attention is given to “small” details that significantly influence optimization convergence and results quality, yet these details are rarely presented in journal publications. Currently, most researchers can learn how to use tools such as OpenSim through a set of tutorials that explore the main tool functionality but lack explanation of the finer details that are critical for research tutorials. While extensive details are useful, including them makes tutorial development much harder and time consuming. Development of the in-depth tutorials presented here took three individuals a year and a half, which is not realistic for every software tutorial. These tutorials are based on real research first, and then the instructions were written after results were obtained. This process required working through the simulation pipeline numerous times to verify that the instructions work as intended and to test different scenarios that could impact optimization convergence. The result is a pair of comprehensive tutorial instructions for how to approach computational treatment design and how to use the NMSM Pipeline for this purpose.

The knee OA tutorial walks users through successfully reproducing the validated results in (21) or predicting the functional outcome of an HTO surgery. Reproducing the results of (21) required a specific problem formulation in which foot position and orientation were constrained. The original paper did not use a ground contact model but rather prescribed the motion of the feet with respect to ground and applied slightly variable ground reactions to them. The HTO solution is not validated, but the optimal correction of 6 degrees falls within the typical range of correction for HTO surgery (90).

The two knee OA treatment predictions have limitations that may affect their prediction accuracy. First, the HTO surgery was simulated by simply changing the knee adduction angle. However, the actual surgery adjusts the angle of the tibia below the knee joint. This limitation could be addressed by adding a specific HTO joint to the tibia that anatomically represents the surgery. Next, the surgery simulation did not use muscles, which could result in the predictive simulation producing a motion that the subject’s neural control cannot achieve. While the predicted medial thrust gait motion has been experimentally validated for a single subject (21), there is no guarantee that other subjects will be able to achieve it. For both treatment problems, it is possible that including personalized muscle-tendon models and a personalized neural control model could yield more accurate predictions. Nevertheless, both the HTO and MTG simulations as performed in the tutorials produced physiologically realistic results.

The stroke tutorial successfully walks users through designing a synergy-based FES intervention to correct gait propulsive force asymmetry. The treatment design in the tutorial relies on two key assumptions that are not common in the literature. First, we assumed that synergy vectors are nearly identical between the two legs. This assumption relies on the hypothesis that synergy activations are sourced from the brain while synergy vectors reside in the spinal cord (91,92). Previous studies have reported that when muscle synergies are calculated from EMG data for walking, synergy vectors are “remarkably similar” between paretic legs of stroke survivors, non-paretic legs of stroke survivors, and legs of healthy subjects when analyzed using the same number of synergies (93,94). As stated in Clark *et al.* (2010), “Indeed, when the data for all post-stroke participants were analyzed using four modules (even when our VAF criteria could be met with fewer modules), we found that the module muscle weightings were remarkably similar between the healthy and post-stroke groups.” Thus, the primary differences between these three groups may occur in the corresponding time-varying synergy activations. From this perspective, while “merged” synergies may be a more compact representation of some post-stroke walking EMG patterns (95), “merging” may actually be “mathematically masking” remaining potential within the post-stroke central nervous system. These observations suggest that it is at least reasonable to explore computational treatment design to improve walking post-stroke using similar synergy vectors between the two legs.

Second, we assumed that synergy activation pairs (i.e., corresponding synergy activations in the two legs) can be classified as either impaired-compensatory, compensatory, or healthy based only on their relative amplitudes between the two legs (67,68). An impaired-compensatory synergy pair possesses a lower amplitude synergy activation on the paretic side and a higher amplitude synergy activation on the non-paretic side. The lower amplitude synergy activation on the paretic side is assumed to be impaired by the stroke and the higher amplitude synergy activation on the non-paretic side is assumed to be compensating for the corresponding impaired synergy activation. In contrast, a compensatory synergy activation pair possesses a higher amplitude synergy activation on the paretic side and a lower amplitude synergy activation on the non-paretic side. The higher amplitude synergy activation on the paretic side is assumed to be compensating for impaired synergy activations on the paretic side, while the lower amplitude synergy activation on the non-paretic side is assumed to be compensating as well. A healthy synergy activation pair possesses comparable amplitude synergy activations on the paretic and non-paretic side. These definitions emerge from the decision to use nearly identical synergy vectors for the two legs. Based on these definitions, when the subject’s impaired synergies (Right Leg 2, Right Leg 3, Right Leg 4) were “stimulated” in the predictive simulation, we assumed that the subject a) would be able to self-correct his compensatory synergy activations (Right Leg 1, Left Leg 1, Left Leg 2, Left Leg 3, Left Leg 4) back to healthy bilateral symmetry in response to the stimulation, which might not be possible depending on the severity of the subject’s neural control impairment, and b) would not alter his healthy synergy activations (Right Leg 5, Left Leg 5) substantially.

An important question is whether these three categories (impaired-compensatory, compensatory, or healthy) for non-paretic and paretic leg synergy activation pairs are supported by experimental evidence from post-stroke FES studies. Skvortsov *et al.* (2025) (96) reported that FES applied to paretic leg muscles for a single session produced a significant increase in knee joint range of motion and a significant decrease in tibialis anterior activation in the non-paretic leg. Similarly, Palmer *et al.* (2017) (97) showed that after one session of FES, the ankle moment showed increased symmetry between the two legs, primarily due to an increase in the paretic ankle moment but also due to a decrease in the non-paretic ankle moment. Taken together, these two studies indicate that paretic leg FES can produce changes in non-paretic leg muscle activations, joint kinematics, and joint moments, consistent with the way we allowed non-paretic leg synergy activations to change in our predictive simulations. Interestingly, when we performed an additional Design Optimization run that did not allow non-paretic leg synergy activations to change in response to paretic leg FES, the optimization would not converge. Equalizing propulsive and braking impulses for the paretic but not the non-paretic leg resulted in too much propulsive impulse to maintain the same walking speed. In addition, Lim *et al.* (2021) (98) reported that paretic leg FES can decrease the amplitude of at least one paretic leg synergy activation, which is counterintuitive given that FES can only increase muscle activations. This finding is consistent with the way we allowed paretic leg synergy activation 1, which we categorized as compensatory, to decrease in our predictive simulations. While the way we simulated non-paretic and paretic leg synergy activation changes is consistent with the limited information available in the literature, future studies should explore in more detail how synergy activations change in both legs in response to paretic leg FES.

Another limitation with our method for simulating FES is that most research laboratories can stimulate only up to eight muscles per leg (67,68,99), which makes stimulating an entire synergy as done in this tutorial unrealistic. In a clinical setting, the primary muscles in the impaired synergies would need to be identified and stimulated. These limitations are significant and highlight that the synergy-based FES treatments designed with this computational method need to be evaluated experimentally.

The stroke tutorial works through the model personalization and treatment optimization processes using the same dataset as in the NMSM Pipeline article but with substantial differences (60). Firstly, the OpenSim model used in the present tutorial was updated from the original article, featuring an updated subtalar joint, knee joint kinematics, knee muscle geometry paths, initial model muscle-tendon model parameter values (100), and muscle moment arms for hip adductor muscles (101). The Joint Model Personalization tool for this article personalized the knee, ankle, and subtalar joints simultaneously using functional joint trials and a gait trial, as opposed to (60) which personalized only the knee and ankle joints using a single gait trial. The Ground Contact Model Personalization tool was re-run using a Coulomb friction model and corrected the electrical center of the force plates relative to the laboratory coordinate system, neither of which were done in the original article. The Muscle-tendon Model Personalization tool was re-run with the new Joint Model Personalization results and new OpenSim model. The iterative process to achieve good Muscle-tendon Model Personalization results was documented as well, providing more insight than did the NMSM Pipeline article into how to achieve the best Muscle-tendon Model Personalization results. The Neural Control Model Personalization tool was re-run with 5 muscle synergies per leg instead of 6 as in the NMSM Pipeline article. Finally, the Design Optimization process was formulated significantly differently. The NMSM Pipeline article used the Design Optimization tool to achieve a target value of metabolic cost and equalized muscle synergies using a complex user defined cost term. The present tutorial aimed to equalize the propulsive and braking impulses between the subject’s feet to achieve improved propulsion by simply scaling synergy activation amplitudes to the average value between legs. When users work through the tutorial, they are also given freedom for how to scale the synergies so that they can compare the differences between equalization methods.

The methods in these tutorials are formulated to work for these specific clinical problems with these specific subjects. They are not guaranteed to work for every clinical problem, or even for every subject having the clinical problems presented in this article. Rather, the goal of the tutorials is to teach users how to utilize the full breadth of NMSM Pipeline functionality for advanced problems encountered in actual research. Attention is given to **how** settings files are created, with details explaining each selected setting, and **why** settings file options are used so that users can extrapolate the reasoning to their problems. However, how each NMSM Pipeline tool run is formulated remains subjective and will be informed by the unique characteristics of the subject and clinical condition being modeled. These tutorials give examples of using personalized neuromusculoskeletal models to diagnose impairments and design treatments, but as with most clinical problems, subjective decision making is still an important part of the treatment design process. Responsibility falls on individual researchers to determine if the NMSM Pipeline can be used to address their research or clinical problems.

An important question for this study is the extent to which the presented research is truly reproducible. If a user starts with the provided generic OpenSim model and the provided experimental data, reads the tutorial instructions, uses the provided NMSM Pipeline and OpenSim XML settings files created by the authors, and runs each NMSM Pipeline and OpenSim tool in the correct order, the presented research will be 100% reproducible. To our knowledge, no previously published neuromusculoskeletal modeling study has demonstrated 100% reproducibility, providing readers with everything needed to go from initial model and data to final model and results. In contrast, if a user starts with the same model and data, reads the tutorial instructions, creates their own XML settings files based on the tutorial instructions, and runs each NMSM Pipeline and OpenSim tool in the correct order, the presented research will be less than 100% reproducible due to flexibility or ambiguity in some tutorial instructions or misunderstanding of some tutorial instructions. To assess how much less, we evaluated the tutorial assignments submitted by students in the combined undergraduate/graduate mechanical engineering course (see Additional File 4.pdf, Supplementary Material D: Reproducibility Evaluation). We found that for each module in both tutorials, between 64% and 95% of students were able to completely or closely reproduce the results generated by the authors, thereby demonstrating strong reproducibility. For further discussion on the reproducibility of our research results, please refer to the reproducibility information provided in Additional File 4.pdf.

## Conclusions

The detailed tutorials presented in this study serve as an introduction to developing and using personalized predictive simulations to design subject-specific treatments for walking impairments using the NMSM Pipeline. The NMSM Pipeline has made predictive simulations and computational treatment design using personalized neuromusculoskeletal models more accessible than was previously possible, enabling these tutorials to be developed. The key learning outcome from these tutorials is a working knowledge of personalized neuromusculoskeletal modeling techniques within the NMSM Pipeline and OpenSim so that researchers can apply this knowledge to their own problems. These tutorials are accessible to beginning users of the NMSM Pipeline and do not require any coding knowledge, making them valuable for both clinicians and newcomers to the biomechanics field. We hope that by working through these tutorials, users will develop an enhanced understanding of the NMSM Pipeline and will be able to apply the NMSM Pipeline’s tools to their own research and clinical problems. The tutorial documents can be found in the supplementary materials of this publication (Additional File 1.pdf and Additional File 2.pdf) and on the NMSM Pipeline website https://nmsm.rice.edu. Tutorial materials on the website will be updated periodically to keep up with new functionality as new features for the NMSM Pipeline are released.

## Supplementary Information

The most recent version of all tutorials described in this article can be found at https://nmsm.rice.edu. All supplementary materials are described below:

File name: Additional File 1.pdf

File format: .pdf

Title: Supplementary Material A - Tutorial 1

Description: The document containing all steps for the first tutorial.

File name: Additional File 2.pdf

File format: .pdf

Title: Supplementary Material B - Tutorial 2

Description: The document containing all steps for the second tutorial.

File name: Additional file 3.pdf

File format: .pdf

Title: Supplementary Material C - Scaled Generic Model Comparison

Description: A document comparing how well scaled generic OpenSim models and NMSM Pipeline personalized models can reproduce experimental walking data.

File name: Additional File 4.pdf

File format: .pdf

Title: Supplementary Material D - Reproducibility Evaluation

Description: A document containing an analysis of how well tutorial results could be reproduced by undergraduate and graduate students at Rice University.

## Declarations

### Ethical approval and consent to participate

The de-identified experimental walking data used in this study were collected and disseminated as part of previously published studies (23,62). Student feedback on the tutorials was collected via a survey that was exempted by the Rice University institutional review board.

### Consent for publication

Not applicable

### Availability of data and materials

The NMSM Pipeline software, along with the tutorial documents, data, models, and settings files described in this article, are available from the “Neuromusculoskeletal Modeling (NMSM) Pipeline” project on SimTK.org (https://simtk.org/projects/nmsm). Users are welcome to ask questions, share feedback, and request new functionality through this project’s forum. Documentation including implementation details, examples, and tutorials is available at https://nmsm.rice.edu.

### Authors contributions

R.S. and B.F. wrote the manuscript text; G.L. and S.W. provided extensive feedback on the manuscript text; B.F. wrote the knee OA tutorial and designed the simulations used in the tutorial; R.S. wrote the stroke tutorial; R.S. and G.L. prepared the simulations used in the stroke tutorial; R.S. prepared manuscript figures; S.W. wrote much of the code used for the tutorials; R.S., G.L, S.W., and B.F. provided input on the tutorial design; B.F. oversaw the research projects on which the tutorials were based.

## Supporting information

All Supplementary Materials

## Acknowledgements

The authors would like to thank the students of MECH497/597 at Rice University during fall semester 2025 for their valuable input on refining the tutorials presented in this article.

## List of abbreviations

NMSM: Neuromusculoskeletal modeling
NMS: Neuromusculoskeletal
OA: Osteoarthritis
HTO: High Tibial Osteotomy
MTG: Medial Thrust Gait
KAM: Knee Adduction Moment
EMG: Electromyography
FES: Functional Electrical Stimulation
API: Application Programming Interface
VAF: Variance Accounted For
RMSE: Root Mean Squared Error

