## Supplementary material for "Reproducible Research: Computational Design of Personalized Clinical Treatments for Walking Impairments Caused by Knee Osteoarthritis and Stroke": All Supplementary Materials

### Supplementary Material A: Tutorial 1

#### TORQUE-DRIVEN SKELETAL MODEL TREATMENT OPTIMIZATION NMSM Pipeline Advanced Tutorial 1

**Tutorial Developer:** B.J. Fregly, Rice Computational Neuromechanics Lab, Rice University

##### Simulation Project Materials

The materials for this tutorial can be downloaded from SimTK at https://simtk.org/projects/nmsm in the “NMSM Advanced Tutorials” section.

##### Simulation Project Overview

The goal of this simulation project is to teach you the Neuromusculoskeletal Modeling (NMSM) Pipeline’s computational treatment design process for clinical applications (Fregly, 2021; Hammond *et al.*, 2025) where use of a personalized torque-driven skeletal model would be sufficient and modeling of muscles or neural control would not be needed. Specifically for this project, you will a) develop a personalized torque-driven three-dimensional skeletal model and then b) use the personalized model to design a modified walking motion as well as a high tibial osteotomy surgical plan to reduce both peak adduction moment peaks for a subject with bilateral medial compartment knee osteoarthritis. This project will involve similar tasks to those performed by Prof. Fregly in his 2007 journal article “Design of Patient-specific Gait Modifications for Knee Osteoarthritis Rehabilitation” (Fregly *et al.*, 2007), where the peak knee adduction moment was used as a surrogate measure for medial knee contact force. The specific treatment design goals for the modified walking motion and high tibial osteotomy surgery are provided in the project details below.

The five tools that you will use within the NMSM Pipeline are indicated in the table below, along with the abbreviations used to reference each tool and required supporting OpenSim tools:

| NMSM Pipeline Toolset | NMSM Pipeline Tool | Used |
| --- | --- | --- |
| Model Personalization | Joint Model Personalization (JMP)<br>(with OpenSim Scale Model tool) | ✓ |
|  | Muscle-tendon Model Personalization (MTP) |  |
|  | Neural Control Model Personalization (NCP) |  |
|  | Ground Contact Model Personalization (GCP)<br>(with OpenSim Inverse Kinematics tool) | ✓ |
| Treatment Optimization | Tracking Optimization (TO)<br>(with OpenSim Inverse Dynamics tool) | ✓ |
|  | Verification Optimization (VO) | ✓ |
|  | Design Optimization (DO) | ✓ |

For the **Model Personalization** toolset, since this simulation lab will require only a personalized skeletal model without including personalized models of muscle-tendon actuators or neural control, you will need only the **Joint Model Personalization** tool and the **Ground Contact Model Personalization** tool. In contrast, for the **Treatment Optimization** toolset, you will still need all three available tools, where your personalized model will be controlled by torque actuators due to the omission of muscles.

This simulation project is broken down into three modules, one for each **Model Personalization** tool and a third for all three **Treatment Optimization** tools. For each module, detailed instructions are provided below to walk you through all the necessary steps. To ensure that poor results for one module do not affect your ability to complete subsequent modules, final results for each module are provided to use as the starting point for the next module if necessary.

To run each required OpenSim or NMSM Pipeline tool, you will generate an initial **xml** settings file using the appropriate tool selection within the OpenSim GUI Tools menu. Once you have generated an initial tool settings file in the OpenSim GUI, you can edit the settings file for subsequent tool runs either within the OpenSim GUI or using a text editor. Runs for OpenSim tools will be performed through the OpenSim GUI, while runs for NMSM Pipeline tools will be performed in Matlab.

The full-body OpenSim model **Full\_Body\_Walking\_Model.osim** that you will use for this project contains slightly modified knee joint models. Instead of having the knee joint connect the femur body to the tibia body, the knee joint in the slightly modified model connects the femur body to a non-standard proximal tibia body. The proximal tibia body is then connected to the tibia body via a custom joint that allows only X axis rotation, which is locked. OpenSim can calculate inverse dynamics loads only about joint axes present in the model. Since the knee adduction moment should be calculated about the X axis of the tibia body, adding this locked custom joint provides the correct joint axis direction for calculating the adduction moment for each knee.

All experimental marker motion and ground reaction data needed to complete this simulation project have been pre-processed for you and are ready to use without further modification. The experimental data that you will need for this project come from the following trials summarized below:

• Trial04\_Static – standing static trial
• Trial06\_AnkleR – isolated right ankle motion trial
• Trial07\_KneeR – isolated right knee motion trial
• Trial08\_HipR – isolated right hip motion trial
• Trial09\_AnkleL – isolated left ankle motion trial
• Trial10\_KneeL – isolated left knee motion trial
• Trial11\_HipL – isolated left hip motion trial
• Trial12\_Gait – walking trial collected at self-selected speed of 1.2 m/s

Marker motion data were collected using a Vicon video-based motion capture system (Vicon Corporation, Oxford, United Kingdom), while ground reaction data were collected using a Bertec split-belt instrumented treadmill (Bertec Corporation, Columbus, OH, United States) with belts tied to the same speed. The experimental data from each trial above have already been converted so that all length data are in units of meters, all moment data are in units of Newton-meters, and all data are reported using coordinate axes consistent with OpenSim model conventions (i.e., +X is directed anteriorly, +Y is directed superiorly, and +Z is directed out to the right).

The specific datafiles that you need to complete the project are organized for you in the **Data** folder. For each OpenSim or NMSM Pipeline tool run that you need to perform, a sub-directory within the **Data** folder is provided containing all of the pre-processed data that you will need. For example, for the OpenSim **Scale Model** tool run described below, all necessary data are provided in the **Scale Model** folder.

#### MODULE 1: JOINT MODEL PERSONALIZATION

In this module, you will use the OpenSim Scale Model tool and the NMSM Pipeline **Joint Model Personalization** (JMP) tool to personalize lower body joint functional axis positions and orientations in a scaled generic full-body OpenSim model. The personalization process will be performed using marker motion data obtained from the isolated joint motion trials as well as the gait trial.

##### Module Task 1: Model Scaling

The starting point for **Joint Model Personalization** is always a scaled generic OpenSim model (or, when available, an OpenSim model possessing subject-specific bone models obtained from the subject's imaging data). For the present project, you will scale a generic full-body OpenSim model (Ragagopal *et al.*, 2016) modified to improve kinematic modeling of the ankles and knees (van den Bogert *et al.*, 1994; Hammond *et al.*, 2025) and musculoskeletal geometry modeling of the knees and hips (Lai *et al.*, 2017; Ulrich *et al.*, 2022). However, to improve the subsequent **Ground Contact Model Personalization** process, you will follow a non-standard model scaling process as described below. The interaction between tool settings, data, and models required to perform this module task is shown in the figure below:

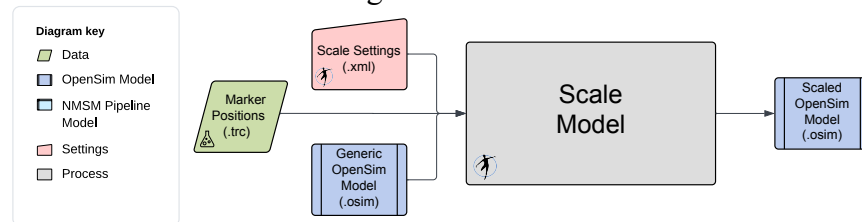

To perform the non-standard model scaling process, you will work with two generic OpenSim models:

1) **Full\_Body\_Walking\_Model.osim** – This model is the generic full-body OpenSim model described above.

To facilitate the model scaling process, some joints are already locked in the model that are typically unlocked and unlocked other joints in the model that are typically locked, as summarized in the tables below.

Notable locked joints in the model include the following:

| Joint | To Be Unlocked Later | To Remain Locked |
| --- | --- | --- |
| pelvis_list | ✓* |  |
| knee_adduction_r |  | ✓ |
| Knee_adduction_l |  | ✓ |
| mtp_angle_r | ✓ |  |
| mtp_angle_l | ✓ |  |
| lumbar_bending | ✓* |  |
| lumbar_rotation | ✓* |  |
| pro_sup_r |  | ✓ |
| pro_sup_l |  | ✓ |

Whether or not the starred (\*) joints should be locked depends on the subject being modeled. Locking these joints during the model scaling process helps ensure that the final model pose in the standing static trial is physically realistic for this particular subject (e.g., the pelvis is level when viewed from the front, the torso is not tilted to the side or twisted about a vertical axis).

Notable unlocked joints in the model include the following:

| Joint | To Be Locked Later | To Remain Unlocked |
| --- | --- | --- |
| knee_frontal_r | ✓ |  |
| knee_frontal_l | ✓ |  |

The knee frontal angles and the knee adduction angles are similar to each other but serve two different purposes. The knee frontal angles are used in the definition of the knee joints. By unlocking these angles during the model scaling process, we ensure that the static frontal plane alignment of both knees in the model is realistic for the subject (i.e., both knees should end up with a slight varus, or bow-legged, alignment) when the model is in the static pose. Having the correct frontal plane knee alignment in the static pose is important since this pose is used to “glue” the experimental markers onto the model’s body segments. If the frontal plane knee alignment is incorrect in the static pose, then the tibial markers will not be placed on the model correctly at the end of the model scaling process. In contrast, the knee adduction angles are used for calculating the knee adduction moments via inverse dynamics. By keeping these angles locked, we create a joint axis that is directed along the X axis of each tibia and that OpenSim can use for calculating a knee adduction moment.

2) **Feet\_Only\_Walking\_Model.osim** – This model contains only the feet from the full-body OpenSim model. This model is needed so that the feet in the model can be properly aligned with the experimental foot marker locations in the static standing trial. If a foot is misaligned with respect to its experimental foot marker locations, then the personalized ground contact model to be developed later for that foot will not work properly.

To construct this model, the following entities are removed from the full-body model:

• All bodies except the right and left calcaneus and toes bodies.
• All joints except the right and left mtp joints.
• All constraints and forces.
• All markers except for markers attached to the calcaneus and toes bodies of each foot.

In addition, following entities are added to the feet-only model:

• Two 6 DOF **Custom Joints** to connect the calcaneus body of each foot to the ground. The rotation sequence for the right calcaneus with respect to ground is defined as (-X, -Y, Z), while the rotation sequence for the left calcaneus with respect to ground is defined as (X, Y, Z). In this way, the same rotations applied to both feet produce anatomically consistent poses for the two feet (e.g., both feet toed out by the same amount). Both X rotations and both Z rotations are locked to zero to prevent each foot from tilting to one side or the other and frontward or backward.
• A **Coordinate Coupler Constraint** to make the Y translation of the left calcaneus the same as that of the right calcaneus. This constraints ensures that both feet are the same

height above the ground in the standing static pose.

The only experimental data needed for the model scaling process is marker data contained in the static trial datafile `Trial04_Static_markers_reordered.trc`. In addition, you will need to know the mass of the subject, which was 72.8 kg based on force plate data collected during the static standing trial.

When iterating the model scaling steps outlined below, you should always select the **Preview static pose (no marker movement)** checkbox at the bottom of the **Scale Tool** menu. This selection will cause the model scaling process to be performed but without replacing markers on the scaled model with the experimental marker locations. You should then select **File** ⇒ **Preview Experimental Data** in the OpenSim GUI to visualize the experimental marker data from the static trial. You can then compare how far the markers on the scaled model are away from the corresponding experimental marker locations.

##### Step 1: Perform model scaling for the full-body model

• Load the full-body model `Full_Body_Walking_Model.osim` into the OpenSim GUI.
• Select the **Scale Model** tool to scale the full-body model using the static trial datafile `Trial04_Static_markers_reordered.trc`.
• On the main **Scale Tool** menu, make the output model name **Full\_Body\_Walking\_Model-Scaled**, select **Scale Model** and **Adjust Model Markers**, and pick a small time window of roughly 0.1 sec for averaging the static trial marker data. Set the scaled model mass to 72.8 kg, and check **Preserve mass distribution during scale**.
• On the **Scale Factors** tab, create measurements called **PelvisWidth** (for scaling the pelvis body) using the R\_ASIS and L\_ASIS markers, **FemurLengthR** (for scaling the right femur and patella bodies) using the R\_ASIS and R\_Knee\_Lateral markers, **TibiaLengthR** (for scaling the right tibia body) using the R\_Knee\_Lateral and R\_Ankle\_Lateral markers, **FemurLengthL** (for scaling the left femur and patella bodies) using the L\_ASIS and L\_Knee\_Lateral markers, **TibiaLengthL** (for scaling the left tibia body) using the L\_Knee\_Lateral and L\_Ankle\_Lateral markers, **FootLength** (for scaling the talus, calcaneus, and toes bodies on both legs) using heel and toe markers on both feet, and **ForearmLength** (for scaling the radius, ulna, and hands on both arms) using the elbow and wrist markers on both arms. You will use different femur and tibia lengths for the two legs since the subject had a slight leg length discrepancy. Do not scale the pretalus body in either leg, since it is a massless intermediate reference frame that only adds a rotational offset as needed to implement the van den Bogert *et al.* (1994) ankle model. Since the shoulder markers were not placed on the shoulder but rather on the straps of a safety harness behind the shoulders, do not use the shoulder markers to create a measurement for estimating torso height or upper arm length. Instead, input manual scale factors for torso body and the humerus bodies, and try different values until your scaled model matches the elbow and wrist marker positions well. You will probably need to make the scale factor for the torso body less than 1 and the scale factor for the two humerus bodies greater than 1.
• On the **Static Pose Weights** tab under **Marker Name**, select markers and associated weights as follows:
  • The heel and toe markers on both feet using a weight of 10 to cause experimental foot marker positions to be matched closely. Do not pick any other foot markers. Note that the

feet will not invert or evert during the scaling process since the subtalar joint axis on both feet has been temporarily locked to a value of 0.

• All medial and lateral ankle and knee markers using a weight of 1.
• The two ASIS markers and the central sacral marker using a weight of 10 to cause experimental pelvis marker positions to be matched closely.
• The chest marker and the two elbow and wrist markers using a weight of 1.

By matching the foot and pelvis markers closely, we will ensure that the lower body joint angles represent the subject's experimental leg position well. Since the shoulder marker positions are not closely related to the anatomical shoulder locations, do not select the shoulder markers on this tab.

• Run the **Scale Model** tool to create a scaled full-body model. Note that this scaled model will have adjusted mass properties.
• Click on the **Navigator** tab and save the scaled model by right-clicking on it and selecting **Save As**. Use `Full_Body_Walking_Model-Scaled.osim` as the saved model name.

##### Step 2: Perform model scaling for the feet-only model

• Load the feet-only model `Feet_Only_Walking_Model.osim` into the OpenSim GUI.
• Select the **Scale Model** tool to scale both feet using the static trial datafile `Trial04_Static_markers_reordered.trc`.
• On the main **Scale Tool** menu, make the output model name **Feet\_Only\_Walking\_Model-Scaled**, select **Scale Model** and **Adjust Model Markers**, and pick the same small time window of 0.1 sec for averaging the static trial marker data. Don't worry about the scaled model mass at this point.
• On the **Scale Factors** tab, create a **FootLength** scale factor using the heel and toe markers for both feet, just like for the full-body model.
• On the **Static Pose Weights** tab, select only the heel and toe markers on each foot, use a weight of 1 for the heel markers, and use a weight of 10 for the toe markers. Give the toe markers more weight since we trust their height above the floor more than we trust the height of the heel markers above the floor.
• Run the **Scale Model** tool to create the scaled model with the experimental markers properly attached to each foot.
• Save the scaled model by right-clicking on it and selecting **Save As**. Use `Feet_Only_Walking_Model-Scaled.osim` as the saved model name.

##### Step 3: Adjust the scaled full-body model for subsequent tasks

Several remaining adjustments must be made to the scaled full-body OpenSim model `Full_Body_Walking_Model-Scaled.osim` before Model Personalization and Treatment Optimization tasks can be performed:

• Copy the scaled OpenSim model `Full_Body_Walking_Model-Scaled.osim` to a new file called `Full_Body_Walking_Model-Scaled_Adjusted.osim`.
• Open model `Full_Body_Walking_Model-Scaled_Adjusted.osim` in a text editor and make the following three changes:
  • Change the model name at the top of the file to `Full_Body_Walking_Model-Scaled_Adjusted`.

• Correct an OpenSim bug where the `<scale>` field in some `<TransformAxis>` blocks contains a negative number. Search for `<scale>-` in your model and replace with `<scale>` (i.e., minus sign deleted) to eliminate non-physical negative scale factors. This OpenSim bug should be fixed in version 4.6 whenever it is released.
• Copy the foot marker locations from OpenSim model `Feet_Only_Walking_Model-Scaled.osim` and replace the corresponding foot marker locations in OpenSim model `Full_Body_Walking_Model-Scaled_Adjusted.osim`, which ensures that the foot marker locations in the full-body model are as consistent as possible with foot marker locations when both feet were flat on the ground.
• Load model `Full_Body_Walking_Model-Scaled_Adjusted.osim` into the OpenSim GUI.
• On the **Coordinates** tab, unlock the following locked coordinates in the model:
  • `pelvis_list`
  • `mtp_angle_r`
  • `mtp_angle_l`
  • `lumbar_bending`
  • `lumbar_rotation`
• On the **Coordinates** tab, set the value of the following unlocked coordinates to 0 and then lock them:
  • `knee_frontal_r`
  • `knee_frontal_l`
• Re-save the modified model as `Full_Body_Walking_Model-Scaled_Adjusted.osim`.

#### Module Task 2: Joint Model Personalization

Now that you have an appropriately scaled full-body walking model with correctly placed markers on the body segments (especially the feet), you are ready to personalize the lower body kinematic structure of your model using the NMSM Pipeline's **Joint Model Personalization** tool. The interaction between tool settings, data, and models required to perform this module task is shown in the figure below:

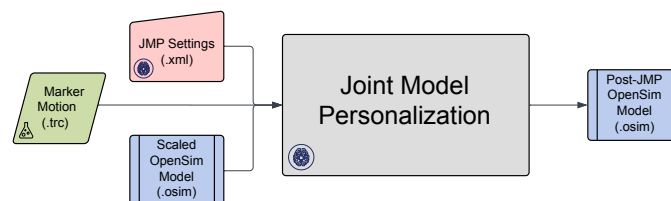

The experimental data needed for the **Joint Model Personalization** process is marker data contained in the isolated joint motion, gait, and combined datafiles located in the **JMP** directory and listed below:

• `Trial06_AnkleR_markers_cropped.trc` – 101 time points of marker data for one cycle of isolated right ankle joint motion
• `Trial07_KneeR_markers_cropped.trc` – 101 time points of marker data for one cycle of isolated right knee motion
• `Trial08_HipR_markers_cropped.trc` – 101 time points of marker data for one cycle of isolated right hip motion
• `Trial09_AnkleL_markers_cropped.trc` – 101 time points of marker data for one cycle of isolated left ankle motion

• `Trial10_KneeL_markers_cropped.trc` – 101 time points of marker data for one cycle of isolated left knee motion
• `Trial11_HipL_markers_cropped.trc` – 101 time points of marker data for one cycle of isolated left hip motion
• `Trial12_Gait_LowerBody_markers_cropped.trc` – 101 time points of marker data for one cycle of reference walking motion
• `Trial04_Static_markers_cropped.trc` – 5 time frames of marker data from the static standing trial
• `Trials06_07_08_LegR_markers_cropped.trc` – 303 time points of concatenated marker data from all isolated joint motion trials for the right leg only
• `Trials09_10_11_LegL_markers_cropped.trc` – 303 time points of concatenated marker data from all isolated joint motion trials for the left leg only
• `Trials06_07_08_09_10_11_BothLegs_markers_cropped.trc` – 606 time points of concatenated marker data from all isolated joint motion trials for both legs
• `Trials08_11_BothHips_markers_cropped.trc` – 202 time points of concatenated marker data from the isolated hip motion trials for both legs
• `Trials_Joints_Gait_markers_cropped.trc` – 707 time points of concatenated marker data from all isolated joint motion trials plus the gait trial

Each datafile contains only lower body markers from the pelvis down, as these markers are the only ones needed to personalize lower body joint functional axes and joint centers. The marker datafile for each type of motion has been splined and cropped to 101 time points representing one complete motion cycle. In this way, each type of motion contributes the same number of marker distance errors to the JMP optimization process when datafiles from multiple types of motions are concatenated into a single combined datafile.

For this module task, you will perform the **Joint Model Personalization** process three times for all lower body joints in the scaled adjusted model by following a three-step process:

**Step 1:** You will personalize the lower body joints *individually*. In this step, you will perform the personalization process for one (ankle and knee) or two (hips) joints at a time using marker data from the isolated joint motion trials, always running a new JMP task manually starting from the original scaled adjusted OpenSim model. This module step is where you will explore how to formulate an appropriate JMP problem for each type of lower body joint. You will use the knowledge gained in this step to formulate the JMP problems that you will solve in the subsequent two steps.

**Step 2:** You will personalize the lower body joints *sequentially*. In this step, you will perform the personalization process one joint at a time using marker data from the isolated joint motion trials, automatically running a sequence of JMP tasks that always starts from the updated OpenSim model produced by the previous JMP task. The last JMP task in the sequence will utilize the gait trial marker data to personalize all joints together.

**Step 3:** You will personalize the lower body joints *simultaneously*. In this step, you will perform the personalization process for all joints together using a concatenated datafile containing marker data from all isolated joint motion trials plus the gait trial. Only a single JMP task will be used for this step.

With the sequential approach, improving marker tracking accuracy for one joint (e.g., the knee) could potentially worsen marker tracking accuracy for another joint (e.g., the ankle), but the benefit is reduced computation time compared to the simultaneous approach. Thus, for this module task, you will be investigating the accuracy-speed tradeoff between the sequential and the simultaneous approaches.

For all three steps, you will create a JMP settings file using the **Joint Model Personalization** tool in the OpenSim GUI. For each JMP settings file, you should use the following general guidelines when creating a JMP task:

• Select the marker motion datafile that corresponds to the single joint (ankle or knee), two joint (hips), or multiple joint (gait, or all joints + gait) optimization problem you are trying to solve.
• Accept the default time range so that you will also get one complete motion cycle of marker data possessing 101 time points for each type of motion included in the datafile.
• Select markers from the following list of “dynamic” markers so that markers are present on the parent and child bodies of all joints included in a JMP task (e.g., when personalizing the right knee functional axis, select all markers on the right femur and right tibia):
  • R\_ASIS, L\_ASIS, and Sacral
  • R\_Thigh\_Superior, R\_Thigh\_Inferior, R\_Thigh\_Lateral, and R\_Thigh\_Posterior
  • R\_Shank\_Superior, R\_Shank\_Inferior, R\_Shank\_Lateral, and R\_Shank\_Posterior
  • R\_Heel, R\_Midfoot\_Superior, and R\_Midfoot\_Lateral
  • L\_Thigh\_Superior, L\_Thigh\_Inferior, L\_Thigh\_Lateral, and L\_Thigh\_Posterior
  • L\_Shank\_Superior, L\_Shank\_Inferior, L\_Shank\_Lateral, and L\_Shank\_Posterior
  • L\_Heel, L\_Midfoot\_Superior, and L\_Midfoot\_Lateral
• Select joints for which parent and/or child frame rotations are to be adjusted, and select bodies for which uniform scaling is to be applied and/or markers are to be moved along selected body axes.
• For each selected joint, do not select any parent or child frame translations. This decision will prevent joints from disarticulating (e.g., the head of the femur will not pull out of the socket in the pelvis).
• For each selected joint, do not select any parent or child frame rotation about the primary axis of rotation for the joint (e.g., the Z axis for the knee). This decision will prevent the optimization from trying to change a joint orientation that is redundant with motion about the primary joint functional axis.
• For each selected joint, keep the default rotation bounds at 0.5, noting that this value is actually in radians and not degrees.
• To personalize an ankle joint, select the ankle joint and corresponding subtalar joint on the desired side, and do not select any bodies. To personalize a knee joint, select the knee joint on the desired side, and do not select any bodies. To personalize both hip joints together, select the pelvis body and allow it to be scaled, and do not select any joints. When personalizing all joints together, consider selecting the tibia and/or femur bodies and allowing both bodies to be scaled and/or markers to be moved along the body segment X and Y directions.
• Never allow markers to be moved on the pelvis, calcaneus, or toes bodies. We trust the marker locations on these bodies (especially on the feet), so allowing them to move is likely to result in an anatomically unrealistic solution.
• To decide which parent and child frame rotations should be changed for which joints, use the

information provided in the table below:

| Joint | Functional Axis | Parent Frame Rotation |  |  | Child Frame Rotation |  |  |
| --- | --- | --- | --- | --- | --- | --- | --- |
|  |  | X | Y | Z | X | Y | Z |
| Subtalar | X |  | ✓ |  |  | ✓ | ✓ |
| Ankle | Z | ✓ | ✓ |  |  |  |  |
| Knee | Z | ✓ | ✓ |  | ✓ | ✓ |  |
| Hip | N/A |  |  |  |  |  |  |

These parent and child frame rotation changes ensure that the final joint structure remains consistent with the joint's physical anatomy. Feel free to modify these choices if you think that some alternate settings might work better!

• Personalize the orientation of a joint functional axis only if at least ~30 deg of rotation occurs about that axis in the marker data being used for the JMP run (e.g., in the gait trial marker data, the subtalar joint experiences a rotation of < 30 deg, so it's functional axis should not be personalized using gait trial data alone) (Chèze *et al.*, 1998).
• For each JMP settings file that you create through the OpenSim GUI, open the settings file in a text editor, increase the value of `<function_tolerance>` to 1.0e-04, and increase the value of `<max_function_evaluations>` to 1000. These changes will make your JMP runs converge faster while also allowing enough iterations for tasks that require a large number of function evaluations per iteration.

**Minor Bug Note:** If you want to read a previously-created JMP settings file back into the OpenSim GUI to modify it, you will need to re-save your settings file with a new name. If you re-save with the same file name, your changes will not be saved in our original settings file.

Once you have created and saved a JMP settings file, you will run the file and generate results by following the instructions below:

• Open Matlab, load the NMSM Pipeline project if necessary, and change directories to where your JMP settings file is located.
• Run the JMP tool in Matlab using the settings file you just saved by inputting the following commands into Matlab:

```
>> parpool
>> tic
>> JointModelPersonalizationTool('JMP_Settings.xml')
>> toc
```

In this example, the name of the JMP settings file is `JMP_Settings.xml`. The `parpool` command will start a parallel pool of workers (which can take a minute or so) if your computer has multiple cores. The `tic` and `toc` commands will tell you how much wall clock time elapsed between when you started your JMP run and when it finished.

• After completing a JMP run, ALWAYS VISUALIZE YOUR POST-JMP MODEL IN THE OPENSIM GUI!!! When visualizing a post-JMP model, you should not only look at how the model looks in the default pose but also run an OpenSim **Inverse Kinematics** analysis on the post-JMP model using the marker motion `.trc` file used for your JMP optimization. If your settings file allows unrealistic changes in joint positions or orientations, body scaling, or marker locations on the body segments, you may get significantly reduced marker distance errors but an anatomically unrealistic model.

• Plot your pre- and post-JMP marker distance errors using the Matlab function `plotJmpResultsFromSettingsFile.m` as shown below:
  `>> plotJmpResultsFromSettingsFile('JMPSettings.xml')`
  This function will also output the average and maximum marker distance errors for your pre- and post-JMP OpenSim models.

Before performing the three **Joint Model Personalization** steps below, you should review the section describing the **Joint Model Personalization** process in our recently published journal article describing the design and functionality of the NMSM Pipeline (Hammond *et al.*, 2025). That section provides helpful information on how to formulate **Joint Model Personalization** problems that reduce marker tracking errors while respecting the anatomic structure of the joints (e.g., ensuring that the “ball” at the head of the femur does not dislocate from the “cup” in the pelvis).

##### Step 1: Perform joint model personalization for lower body joints individually

• Load the scaled adjusted model `Full_Body_Walking_Model-Scaled_Adjusted.osim` into the OpenSim GUI. NMSM Pipeline tools will not be accessible in the OpenSim GUI **Tools** menu unless a model to personalize is loaded first.
• Select **Tools** ⇒ **User Plugins** ⇒ `rcnlPlugin.dll` to load the NMSM Pipeline tools into the OpenSim GUI **Tools** menu.
• Select the **Joint Model Personalization** tool to set up one JMP task to be saved as one JMP settings file to personalize one joint (or two joints for the hips) at a time. Repeat this process for the remaining joints, always using the same scaled adjusted OpenSim model as the initial model, and always using isolated joint motion data as the input marker motion data. First, personalize the right ankle by creating a JMP settings file just for the right ankle using the isolated joint motion datafile for the right ankle. Next, personalize the right knee by creating a JMP settings file just for the right knee using the isolated joint motion datafile for the right knee. Finally, personalize both hips together by creating a JMP settings file for both hips using the isolated joint motion datafile for both hips. You do not need to personalize the ankle and knee joints on the left side, since performing the process for just the right side will provide the knowledge you need to perform the next two steps.
• Once a JMP settings file is completed, save it to your hard disk using the **Save** command at the bottom of the tool menu. Name these three JMP settings files `JMP_Settings_RAnkle.xml`, `JMP_SettingsRKnee.xml`, and `JMP_Settings_BothHips.xml`.

##### Step 2: Perform joint model personalization for lower body joints sequentially

• Load the scaled adjusted model `Full_Body_Walking_Model-Scaled_Adjusted.osim` into the OpenSim GUI. NMSM Pipeline tools will not be accessible in the OpenSim GUI **Tools** menu unless a model to personalize is loaded first.
• Select **Tools** ⇒ **User Plugins** ⇒ `rcnlPlugin.dll` to load the NMSM Pipeline tools into the OpenSim GUI **Tools** menu (if you have not done so already).
• Select the **Joint Model Personalization** tool to set up a sequence of 6 JMP tasks to be saved as one JMP settings file that personalizes lower body joints one at a time. Each JMP task in the sequence should use the appropriate marker motion data for that task (e.g., the JMP task that personalizes the right ankle should use marker motion data from the isolated right ankle motion trial). As each task in the sequence is completed, the next task will use as its starting

point an updated OpenSim model produced by the previous task. It is up to you to define the JMP task sequence. The only limitation is that the gait trial must be used alone as the final task in the sequence. Possible task sequences from which to choose are listed below:

• AnkleR ⇒ AnkleL ⇒ KneeR ⇒ KneeL ⇒ BothHips ⇒ Gait
• AnkleR ⇒ AnkleL ⇒ BothHips ⇒ KneeR ⇒ KneeL ⇒ Gait
• KneeR ⇒ KneeL ⇒ BothHips ⇒ AnkleR ⇒ AnkleL ⇒ Gait
• KneeR ⇒ KneeL ⇒ AnkleR ⇒ AnkleL ⇒ BothHips ⇒ Gait
• BothHips ⇒ AnkleR ⇒ AnkleL ⇒ KneeR ⇒ KneeL ⇒ Gait
• BothHips ⇒ KneeR ⇒ KneeL ⇒ AnkleR ⇒ AnkleL ⇒ Gait

Select the sequence that you believe has the best chance of producing the lowest marker distance errors. You need to perform the sequential approach for only one sequence!

• Name your output model file **Full\_Body\_Walking\_Model-JMP\_Sequential.osim**
• Once your JMP settings file is completed, save it to your hard disk using the **Save** command at the bottom of the tool menu and name it **JMP\_Settings\_Sequential.xml**.
• Run **Joint Model Personalization** using the instructions provided above.

##### Step 3: Perform joint model personalization for lower body joints simultaneously

• Load the scaled adjusted model **Full\_Body\_Walking\_Model-Scaled\_Adjusted.osim** into the OpenSim GUI.
• Select **Tools** ⇒ **User Plugins** ⇒ **rcnlPlugin.dll** to load the NMSM Pipeline tools into the OpenSim GUI **Tools** menu (if you have not done so already).
• Select the **Joint Model Personalization** tool to set up a single JMP task to be saved as one JMP settings file that personalizes all lower body joints together. The one JMP task should use the marker motion data from each isolated joint motion trial plus the gait trial. In this way, all functional axes of the lower body joints will be exercised within a single datafile. The fact that the motions are discontinuous between one motion and the next is irrelevant, since internally JMP performs repeated OpenSim **Inverse Kinematics** analyses, and these analyses do not require continuity between time frames of marker data.
• Name your output model file **Full\_Body\_Walking\_Model-JMP\_Simultaneous.osim**
• Once your JMP settings file is completed, save it to your hard disk using the **Save** command at the bottom of the tool menu and name it **JMP\_Settings\_Simultaneous.xml**.
• Run **Joint Model Personalization** using the instructions provided above.

##### Step 4: Select the model with the best personalized joints

• Compare the average and maximum marker distance errors for the gait trial produced by the sequential and simultaneous approaches.
• Identify the approach that produced the lowest errors for the gait trial marker data and that is also the most realistic physically (as determined by visual inspection in the OpenSim GUI – e.g., look for approximate bilateral symmetry in joint coordinate system orientations in the body segments, approximate bilateral symmetry in joint angles in the static pose, reasonable body scale factors, reasonable marker location changes).
• Copy the OpenSim model produced by the “best” approach and give it the new name **Full\_Body\_Walking\_Model-Post\_JMP.osim**.
• In a text editor, open the model and change the model name at the top of the file to **Full\_Body\_Walking\_Model-Post\_JMP**.

##### Step 5: Redefine the default pose in preparation for Ground Contact Model Personalization

• Load model `Full_Body_Walking_Model-Post_JMP.osim` into the OpenSim GUI.
• Select the **Inverse Kinematics** tool and run an inverse kinematics analysis using the static trial marker data with the following marker/coordinate selections and weights:
  - Deselect all medial and lateral ankle and knee markers
  - Deselect the non-central sacral markers.
  - Deselect both shoulder markers and the back marker.
  - Select all foot and remaining pelvis markers using a weight of 10.
  - Select all tibia, arm, and chest markers using a weight of 1.
  - Select all thigh markers using a weight of 0.1.
  - Select the `lumbar_rotation` coordinate, leave the **Default Value** as zero, and set the weight to 0.1.

Note that these weights represent good choices when performing an inverse kinematics analysis for a motion trial. Lightly tracking a `lumbar_rotation` coordinate value of 0 will prevent the torso from twisting unrealistically since a single marker near the midline of the torso is not sufficient to make the `lumbar_rotation` unique.

• Make the static pose the new default pose by selecting the **Coordinates** tab, then **Poses>**, and finally **Set Default**. Making this final static pose the new default pose will place the feet in the correct pose and at the correct height above the floor for contact element placement on the bottom of each foot during the Ground Contact Model Personalization process in the next module.
• Save the model in this final pose by right-clicking on it and selecting **Save**.
• Take a screen shot of the model in the final static pose with the experimental markers shown as well.

##### Deliverables

1. OpenSim **Scale Model** tool settings file for scaling `Feet_Only_Walking_Model.osim` along with scaled model file `Feet_Only_Walking_Model-Scaled.osim`.
2. OpenSim **Scale Model** tool settings file for scaling `Full_Body_Walking_Model.osim` along with scaled model file `Full_Body_Walking_Model-Scaled.osim`.
3. Scaled adjusted model file `Full_Body_Walking_Model-Scaled_Adjusted.osim`.
4. Screen shot of model `Full_Body_Walking_Model-Scaled_Adjusted.osim` in the static pose found by inverse kinematics with the experimental markers shown as well (screenshot shown below):

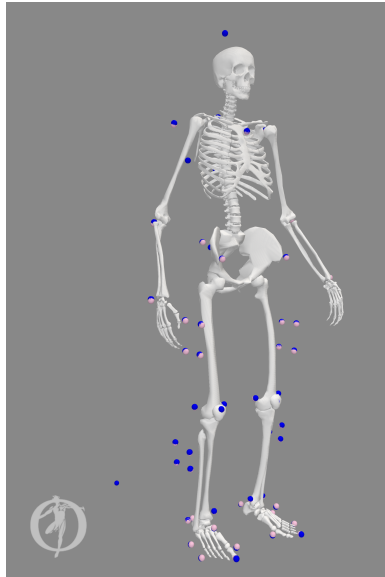

5. Wall clock time along with pre- and post-JMP marker distance errors for each JMP run performed in the three steps above (complete the table below):

| JMP Run | Wall Clock | Pre-JMP Model |  | Post-JMP Model |  |
| --- | --- | --- | --- | --- | --- |
|  | Time (hrs) | Avg Error (m) | Max Error (m) | Avg Error (m) | Max Error (m) |
| 1. Right Ankle |  |  |  |  |  |
| 1. Right Knee |  |  |  |  |  |
| 1. Both Hips |  |  |  |  |  |
| 2. Sequential |  |  |  |  |  |
| 3. Simultaneous |  |  |  |  |  |

6. A plot comparing pre- and post-JMP marker distance errors across all time frames for each of the 5 JMP runs shown in the table above.
7. A JMP settings file for each of the 5 JMP runs shown in the table above.
8. Post-JMP OpenSim model `Full_Body_Walking_Model-JMP_Sequential.osim` produced by the sequential approach and `Full_Body_Walking_Model-JMP_Simultaneous.osim` produced by the simultaneous approach.
9. A description of the task sequence that you chose to use for the sequential approach along with a brief argument for why you believe this sequence might produce lower marker distance errors than would other sequences.
10. A brief paragraph explaining when researchers should use the sequential approach and when they should use the simultaneous approach for **Joint Model Personalization**.

#### MODULE 2: GROUND CONTACT MODEL PERSONALIZATION

In this module, you will use the OpenSim **Inverse Kinematics** tool and the NMSM Pipeline **Ground Contact Model Personalization** (GCP) tool to personalize foot-ground contact model properties in a new NMSM Pipeline model to be associated with your post-JMP OpenSim model. The personalization process will be performed using marker motion and ground reaction data obtained from the gait trial.

##### Module Task 1: Inverse Kinematics for Gait Motion

The starting point for **Ground Contact Model Personalization** is always a post-JMP OpenSim model whose joint positions/orientations in the body segments, body scale factors, and/or marker locations on the body segments have been calibrated using the **Joint Model Personalization** tool. As indicated above, you will use model **Full\_Body\_Walking\_Model-Post\_JMP.osim** as your starting point. You will then perform an OpenSim **Inverse Kinematics** analysis on this model using one cycle of gait trial marker data to generate the input joint motions needed for performing **Ground Contact Model Personalization**. The interaction between tool settings, data, and models required to perform this module task is shown in the figure below:

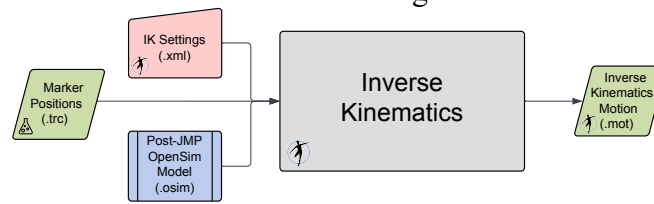

The only experimental data needed for the **Inverse Kinematics** analysis is marker data contained in the gait trial datafile **Trial12\_Gait\_markers.trc**. This file contains data for all experimental markers, including those on the torso and arms.

###### Step 1: Perform inverse kinematics using the gait trial marker data.

• Load model **Full\_Body\_Walking\_Model-Post\_JMP.osim** into the OpenSim GUI and select the **Inverse Kinematics** tool.
• Under **IK Trial**, select the marker data from gait trial **Trial12\_Gait\_markers.trc** and keep the default time range from 0 to 4 seconds.
• On the **Weights** tab, select following marker selections and use the following weights:
  - Deselect all medial and lateral toe, ankle, and knee markers
  - Deselect the non-central sacral markers.
  - Deselect both shoulder markers and the back marker.
  - Select all remaining foot and pelvis markers using a weight of 10.
  - Select all tibia, arm, and chest markers using a weight of 1.
  - Select all thigh markers using a weight of 0.1.
  - Select the **lumbar\_rotation** coordinate, set a **Manual value** of zero, and set the weight to 0.1.
• Name your **Output Motion File** **Trial12\_Gait\_IK\_results.mot**.
• Once your **Inverse Kinematics** settings file is completed, save it to your hard disk using the **Save** command at the bottom of the tool menu and name it **IK\_Settings\_Gait.xml**.

• Run your **Inverse Kinematics** analysis and verify that your results file was written to your hard disk. If not, right click on IK Results in the GUI **Navigator** pane and select **Save As** to save the motion file to your hard disk.
• The inverse kinematics results generated in this step cover more than the gait cycle of interest, since the filtering process to be performed in the next step introduces end effects. These effects distort each filtered curve near the start time and end time, causing significant errors in not only the filtered joint position data but also the joint velocity data to be calculated from it. Taking filtered inverse kinematics data from a time window that is not close to the start time or end time eliminates the presence of end effects caused by the filtering process.

##### Step 2: Filter your inverse kinematics motion file

• Open Matlab and change directories to your **Inverse Kinematics** data folder.
• Run the provided Matlab program **filterIKResults.m** without providing any inputs. The correct default inputs will be used automatically.
• After the program finishes, verify that a new inverse kinematics motion file called **Trial12\_Gait\_IK\_results\_filtered.mot** is now present in your **Inverse Kinematics** data folder, and copy this file to your **GCP** data folder.
• Filtered inverse kinematics results are needed since each GCP run will need to differentiate the IK joint position results to generate joint velocity results. These joint velocity estimates are in turn needed to calculate point velocities relative to ground for the contact elements placed on the bottom of each foot by the **Ground Contact Model Personalization** process. Contact element velocities are inputs to vertical contact force nonlinear damping and horizontal contact force friction calculations.
• After generating your filtered inverse kinematics results file, open two versions of model **Full\_Body\_Walking\_Model-Post\_JMP.osim** in the OpenSim GUI, load the unfiltered motion into the first model, and load the filtered motion into the second model (it can help to make the second a model a different color). Then sync both motions and animate them together to ensure that the motion of the feet produced by the filtered IK results closely follow the motion of the feet produced by the unfiltered IK results.

##### Module Task 2: Ground Contact Model Personalization

Now that you have inverse kinematics data available for your walking trial, you are ready to personalize the ground contact model properties of both feet in your model using the NMSM Pipeline's **Ground Contact Model Personalization** tool. You will also need to use filtered experimental ground reaction data for this task. The interaction between tool settings, data, and models required to perform this module task is shown in the figure below:

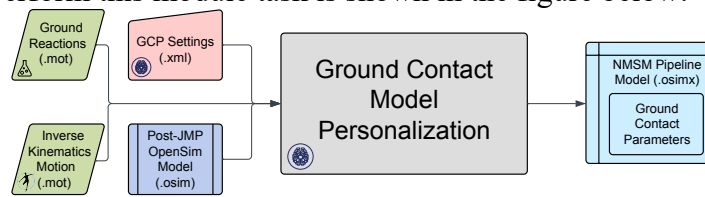

The experimental data needed for the **Ground Contact Model Personalization** process is filtered inverse kinematic joint motion data and filtered ground reaction data contained in the datafiles located in the **GCP** folder and listed below:

• `Trial12_Gait_IK_results_filtered.mot` – filtered joint motion data for multiple walking cycles.
• `Trial12_Gait_forces_filtered.mot` – filtered ground reaction data for multiple walking cycles. Note that the sampling frequency for the ground reaction data has been decreased to match the sampling frequency of the marker and inverse kinematics data, which is a requirement for running the GCP tool.

Each datafile contains data for 4 seconds of walking. You will use only a single cycle of walking data, defined by a specified start time and end time, to personalize foot-ground contact models for both feet. Furthermore, the foot-ground contact model personalization process will be performed such that both feet possess the same contact model stiffness, damping, and friction properties.

Note that experimental marker data are not an input to this tool. Instead, your inverse kinematics results generated with tight tracking of foot markers are used as an input. Consequently, GCP tool runs actually track model (rather than experimental) marker trajectories generated by applying your filtered inverse kinematics motion to your OpenSim model.

For this module task, you will perform the **Ground Contact Model Personalization** process two times for both feet together by following a two-step process:

**Step 1:** You will personalize both feet together assuming Coulomb friction acts on each contact element.

**Step 2:** You will personalize both feet together assuming viscous friction acts on each contact element.

For both steps, you will create a GCP settings file using the **Ground Contact Model Personalization** tool in the OpenSim GUI. For each GCP settings file, you should use the following general guidelines:

• Load the post-JMP model `Full_Body_Walking_Model-Post_JMP.osim` into the OpenSim GUI. NMSM Pipeline tools will not be accessible in the OpenSim GUI **Tools** menu unless a model to personalize is loaded first.
• Select **Tools** ⇒ **User Plugins** ⇒ `rcnlPlugin.dll` to load the NMSM Pipeline tools into the OpenSim GUI **Tools** menu.
• Select the **Ground Contact Model Personalization** tool to set up a sequence of three GCP tasks to be saved as one GCP settings file that personalizes both feet together. Note that you will not create the three GCP tasks in the GUI. A sequence of three default tasks will be created for you automatically when you save your GCP settings file.
• For the **Input Model**, choose your post-JMP model `Full_Body_Walking_Model-Post_JMP.osim`.
• Leave **Osimx File** blank, since you will be creating a new `.osimx` file rather than appending to an existing one.
• For **Input Dir**, pick your `GCP` data folder.
• For **Motion File**, pick `Trial12_Gait_IK_results_filtered.mot`.
• For **Ground Reaction File**, pick `Trial12_Gait_forces_filtered.trc`.
• For **Results Dir**, create a folder called `gcpResults` within your `GCP` data folder and pick that folder.

• Add two **Ground Contact Personalization Surfaces**, one for the right foot and one for the left foot. For both, use a **Time Range** of 2.235 to 3.280 seconds and a **Belt Speed** of 1.2 m/s. For **Force Columns** (**`_vx,y,z`**), **Moment Columns** (**`_mx,y,z`**), and **Electrical Center** (**`_px,y,z`**), use data from force plate 2 for the right foot and from force plate 1 for the left foot. For **Hindfoot Body**, pick the calcaneus **`calcn`** body for the appropriate foot. For various markers, pick the appropriate markers on each foot, noting that **Medial Marker** and **Lateral Marker** represent model markers placed on the toes axis of each foot. For the left foot, make sure to check the **Left Foot** checkbox.
• The **`midfoot_superior`** model marker on each foot is a required input for this settings file since the point used for calculating ground reaction moments is the current location of the midfoot superior marker projected onto the floor. Ground reaction moments can be calculated about any desired point, and this choice of point keeps all three components of ground reaction moment “small” for the GCP calibration process. Choice of a point far from the center of the foot would produce large ground reaction moment components, and errors in those components would then swamp matching ground reaction force components during a GCP run.
• Once you have completed these steps in the OpenSim GUI, **Save** your setting file in your **GCP** folder and name it **`GCP_Settings.xml`**. Note that your saved settings file will already have three pre-defined tasks included in it – a first task that focuses on reproducing just the vertical component of ground reaction force for both feet, a second that focuses on reproducing all three components of ground reaction force for both feet, and a third that focuses on reproducing all three components of ground reaction force and all three components of ground reaction moment for both feet. Each of these tasks contains default parameter values that will work well in most cases, and thus users do not have to create the three individual tasks themselves in the OpenSim GUI.
• To finished configuring settings file **`GCP_Settings.xml`** that you created in the OpenSim GUI, open it in a text editor (ideally Notepad++ on Windows or BBEdit on Mac) and make the following changes to each **`<GCPTask>`**:
  • At the top of the settings file, change the value of **`<kinematics_filter_cutoff>`** to 6.
  • For the first task, change the value of **`<neighborStandardDeviation>`** to 0.3. For **`<RCNLCostTerm>`** items, change **`<max_allowable_error>`** for **`rotation`** to 0.0175, for **`ground_reaction_moment`** to 20, and for **`neighbor_spring_constant`** to 1000. Also add a new cost term called **`kinematic_periodicity`** and set its **`<max_allowable_error>`** to 3, which is a multiple of the ratio between corresponding original initial and final joint positions.
  • For the second task, make the same changes as for the first task plus change **`<max_allowable_error>`** for **`horizontal_grf`** to 5. Keep **`<restingSpringLength>`** set to the default selection of **`false`**.
  • For the third task, make the same changes as for the second task and leave **`<max_allowable_error>`** for **`ground_reaction_moment`** set to 0.5. In addition, set **`<electricalCenterX>`** and **`<electricalCenterZ>`** to **`true`** so that GCP will also calibrate the electrical center location of each force plate in the plane of the force plate.
  • At the bottom of the settings file, where settings are provided that apply to all tasks, change **`<initial_resting_spring_length>`** to 0.01, **`<initial_spring_constant>`** to 6000, **`<initial_damping_factor>`** to 0.5, and **`<initial_dynamic_friction_coefficient>`** to 0.3, **`<diff_min_change>`** to 1e-4, **`<step_tolerance>`** to 1e-5, and **`<max_iterations>`** to 25.

**Minor Bug Note:** If you want to read a previously-created GCP settings file back into the OpenSim GUI to modify it, it will not work. Instead, you will need to modify existing GCP settings files directly using a text editor.

Once you have created and edited a GCP settings file, you will run the file and generate results by following the instructions below:

• Open Matlab, load the NMSM Pipeline project if necessary, and change directories to where your GCP settings file is located.
• Run the GCP tool in Matlab using the settings file you just saved by inputting the following commands into Matlab:

```
>> parpool
>> tic
>> GroundContactPersonalizationTool('GCP_Settings.xml')
>> toc
```

In this example, the name of the GCP settings file is `GCP_Settings.xml`. The `parpool` command will start a parallel pool of workers (which can take a minute or so) if your computer has multiple cores. The `tic` and `toc` commands will tell you how much wall clock time elapsed between when you started your GCP run and when it finished.

• Plot your GCP results using the Matlab function `plotGcpResultsFromSettingsFile.m` as shown below:

```
>> plotGcpResultsFromSettingsFile('GCP_Settings.xml')
```

This function will output root-mean-square errors in matching ground reaction forces, ground reaction moments, hindfoot joint translations, hindfoot joint rotations, and toes joint rotations for your two personalized foot-ground contact models, along with a color plot showing the distribution of spring stiffness values over the bottom surface of the foot.

Before performing the two **Ground Contact Model Personalization** steps below, you should review the section describing the **Ground Contact Model Personalization** process in our recently published journal article describing the design and functionality of the NMSM Pipeline (Hammond *et al.*, 2025). That section provides helpful information on how to formulate **Ground Contact Model Personalization** problems that reduce match ground reaction forces and moments closely without changing the motion of each foot substantially.

##### Step 1: Perform ground contact model personalization using Coulomb friction

• Copy the GCP settings file created above and name it `GCP_Settings_Coulomb.xml`.
• Open this file in a text editor.
• Change the name of the `<results_directory>` to `gcpResultsCoulomb`.
• Run **Ground Contact Model Personalization** with this settings file using the instructions provided above.

##### Step 2: Perform ground contact model personalization using viscous friction

• Copy the original GCP settings file created above and name it `GCP_Settings_Viscous.xml`.
• Open this file in a text editor.
• Change the name of the `<results_directory>` to `gcpResultsViscous`.
• Change `<dynamicFrictionCoefficient>` to `false` everywhere it appears in the file.

• For the second and third tasks, change `<viscousFrictionCoefficient>` to `true`.
• Change the value of `<initial_dynamic_friction_coefficient>` to 0.
• Change the value of `<initial_viscous_friction_coefficient>` to 0.3.
• Run **Ground Contact Model Personalization** with this settings file using the instructions provided above

**Optional Exploration:** For whichever friction model you decide is better, explore changing the following settings to see if they improve the ability of your personalized foot-ground contact models to reproduce the experimental ground reaction force/moment and foot motion data:

• Run GCP without filtering the input inverse kinematics data (i.e., use the original inverse kinematics results file `Trial12_Gait_IK_results.mot`).
• Double the grid density by changing the value of `<grid_width>` to 10 and `<grid_height>` to 22.
• Reduce the value of `<neighborStandardDeviation>` to 0.2.
• Reduce the `<max_allowable_error>` for `neighbor_spring_constant` to 500.
• Reduce the `<latching_velocity>` to 0.05.

##### Deliverables

1. Wall clock time (sec) for your GCP runs with Coulomb friction and viscous friction:
  Coulomb friction wall clock time: \_\_\_\_\_ min
  Viscous friction wall clock time: \_\_\_\_\_ min
2. RMS errors for both feet in matching experimental ground reaction forces (N), ground reaction moments (Nm), toes and hindfoot rotations (deg), and hindfoot translations (m) when using Coulomb friction and viscous friction (complete the table below):

| Quantity | Direction | Coulomb Friction |  | Viscous Friction |  |
| --- | --- | --- | --- | --- | --- |
|  |  | Right Foot | Left Foot | Right Foot | Left Foot |
| RMS Force Error (N) | Anterior |  |  |  |  |
|  | Vertical |  |  |  |  |
|  | Lateral |  |  |  |  |
| RMS Moment Error (Nm) | X |  |  |  |  |
|  | Y |  |  |  |  |
|  | Z |  |  |  |  |
| RMS Rotation Error (deg) | Toes |  |  |  |  |
|  | Y |  |  |  |  |
|  | X |  |  |  |  |
|  | Z |  |  |  |  |
| RMS Translation Error (m) | X |  |  |  |  |
|  | Y |  |  |  |  |
|  | Z |  |  |  |  |

2. For each friction model and each foot, a plot comparing post-GCP ground reaction forces and moments with experimental ground reaction forces and moments.
3. For each friction model and one foot, a color plot showing the distribution of spring stiffness values over the bottom surface of the foot.
4. The GCP settings files for the two GCP runs whose results are shown in the table above.
5. Post-GCP NMSM Pipeline models `Full_Body_Walking_Model-GCP_Coulomb.osimx`

produced by GCP using Coulomb friction and `Full_Body_Walking_Model-GCP_Viscous.osimx` produced by GCP using viscous friction.

6. A brief paragraph explaining which type of friction model – Coulomb or viscous – you think is more physically realistic based on the results in your table above and the various plots produced by the function `plotGcpResultsFromSettingsFile`.

#### MODULE 3: TRACKING OPTIMIZATION

In this module, you will use the NMSM Pipeline **Tracking Optimization** (TO) tool with your personalized skeletal model to create a dynamically consistent walking simulation that reproduces experimental joint motion, joint moment, and ground reaction data as closely as possible for a single gait cycle. **Tracking Optimization** results always provide the starting point for the NMSM Pipeline **Treatment Optimization** process. In the subsequent module, you will perform a **Verification Optimization** (VO) starting from your TO results to confirm that your TO results are reliable, followed by a **Design Optimization** (DO) starting from your VO results to design either a high tibial osteotomy surgery or a modified walking motion that reduces the peak adduction moment in both knees of a subject with bilateral medial knee osteoarthritis.

In preparation for this module, compare the root-mean-square translation, rotation, ground reaction force, and ground reaction moment errors produced using Coulomb and viscous friction in the two GCP runs you performed for the previous module. Select the NMSM Pipeline foot-ground contact model with the most physically realistic results and ideally the lowest errors. For “physically realistic,” look for ground reaction force and moment results that make sense physically (e.g., the ground reaction forces and moments should be zero during swing phase when the foot is off the ground and zero at the transitions into and out of contact). Copy this NMSM Pipeline model and give it the new name `Full_Body_Walking_Model-Post_GCP.osimx`.

The TO run for this module will be performed using the following model and data files:

• OpenSim model file: `Full_Body_Walking_Model-Post_JMP.osim`
• NMSM Pipeline model file: `Full_Body_Walking_Model-Post_GCP.osimx`
• Inverse kinematics data file: `Trial12_Gait_IK_results_filtered.mot` (to be cropped – this data file was an input to, not an output from, **Ground Contact Model Personalization**)
• Ground reaction data file: `updated_Trial12_Gait_forces_filtered.mot` (to be cropped)
• Inverse dynamics data file: `Trial12_Gait_ID_results.sto` (to be created and cropped)

Once all necessary input data files are available, they will be cropped to a single gait cycle between 2.235 to 3.280 seconds, which is the same gait cycle used for the **Ground Contact Model Personalization** process. The cropping process will be performed using Matlab program `cropGaitData.m` provided with the project.

Note that this step uses an updated rather than the original ground reaction data file. This updated file was produced by **Ground Contact Model Personalization** and contains a small shift in the electrical center location of each force plate. The electrical center is the point about which moments are summed for a force plate. The location of this point often contains small errors (especially for split-belt instrumented treadmills) that make the ground reaction data inconsistent with the video motion capture data. **Ground Contact Model Personalization** corrected for this inconsistency during the third GCP task that matched all three components of ground reaction moments as well as all three components of ground reaction forces.

##### Module Task 1: Inverse Dynamics

The interaction between tool settings, data, and models required to perform this module task is shown in the figure below:

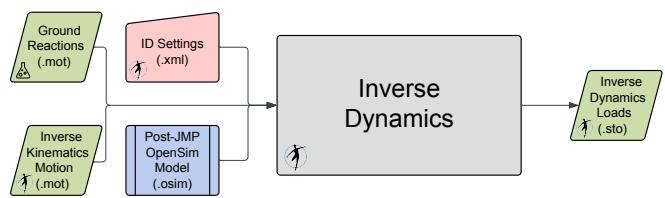

##### Step 1: Perform inverse dynamics using the gait trial experimental data.

• Copy the ground reactions data file **updated\_Trial12\_Gait\_forces\_filtered.mot** from the **GCP\gcpResults\GRFData** folder for your selected friction model to your **Inverse Dynamics** folder and rename it **Trial12\_Gait\_forces\_filtered\_updated.mot**.
• Load model **Full\_Body\_Walking\_Model-Post\_JMP.osim** into the OpenSim GUI and select the **Inverse Dynamics** tool.
• On the **Main Settings** tab, make the following selections:
  - Under **Input**, for the **From ...** option, select your filtered inverse kinematics motion data file **Trial12\_Gait\_IK\_results\_filtered.mot**, check the **Filter coordinates** checkbox, and enter 6 Hz.
  - Under **Time**, keep the default **Time range to process** as 0 to 4 seconds, which was populated when you selected your inverse kinematics motion file.
  - Under **Output**, make the **Directory** your **Inverse Dynamics** data folder.
• On the **External Loads** tab, select the **External Loads** checkbox and click the pencil icon to bring up the **External Forces** set up window. In this window, select the ground reaction data to use as follows:
  - Click the folder icon next to **Force data file** and select your ground reactions data file **Trial12\_Gait\_forces\_filtered\_updated.mot**.
  - Click **Add** to add ground reaction data to the left and right feet. Use force plate 2 data for the right foot and force plate 1 data for the left foot. Make sure the force and torque are applied to the calcaneus body and that the force and point data are expressed in the ground coordinate system.
  - Save your ground reactions **External Forces** settings file (a sub-settings file within inverse dynamics) as **External\_Forces\_Settings\_Gait.xml** using the **Save...** button at the bottom of the **External Forces** window.
• Once your **Inverse Dynamics** settings file is completed, save it to your hard disk using the **Save...** command at the bottom of the tool window and name it **ID\_Settings\_Gait.xml**.
• Run your **Inverse Dynamics** analysis and verify that the default inverse dynamics results file **inverse\_dynamics.sto** was saved to your **Inverse Dynamics** data folder. Rename this file **Trial12\_Gait\_ID\_results.sto**.
• The inverse dynamics results generated in this step cover more than the gait cycle of interest, since the spline fitting process used to calculate joint velocities and accelerations has end effects that distort the values of the derivatives near the start time and end time. Performing inverse dynamics over a larger time window than needed and then cropping the results to the desired time window afterward eliminates these end effects.

##### Step 2: Crop all of your input data files to one gait cycle

• Open Matlab and change directories to your **Inverse Dynamics** data folder.
• Run the provided Matlab program **cropGaitData.m** with no inputs. The program will automatically pick the correct gait trial data and crop it using a start time of 2.235 second and

an end time of 3.280 seconds.

- After the program finishes, verify that new data files with the following names are present in your **Inverse Dynamics** data folder:

- **Trial12\_Gait\_IK\_results\_filtered\_cropped.sto**
- **Trial12\_Gait\_ID\_results\_cropped.sto**
- **Trial12\_Gait\_forces\_filtered\_updated\_cropped.sto**

Note that all three data files now have a **.sto** extension. OpenSim is agnostic between **.mot** and **.sto** file extensions but prefers the **.sto** extension. Consequently, from here forward, **.sto** files will be used throughout the **Treatment Optimization** process.

#### Module Task 2: Tracking Optimization

For this module task, you will use the NMSM Pipeline **Tracking Optimization** tool to produce a one-cycle dynamically consistent walking simulation based on your personalized post-JMP OpenSim model and your associated personalized post-GCP NMSM Pipeline foot-ground contact model. The optimization will seek to spread out errors in matching experimental joint motion from inverse kinematics, ground reaction forces and moments, and joint moments from inverse dynamics while driving the residual forces and torques acting on the pelvis segment to near zero. The personalized OpenSim/NMSM Pipeline skeletal model created by the **Model Personalization** process will provide the starting point for this module. The interaction between tool settings, data, and models required to perform this module task is shown in the figure below:

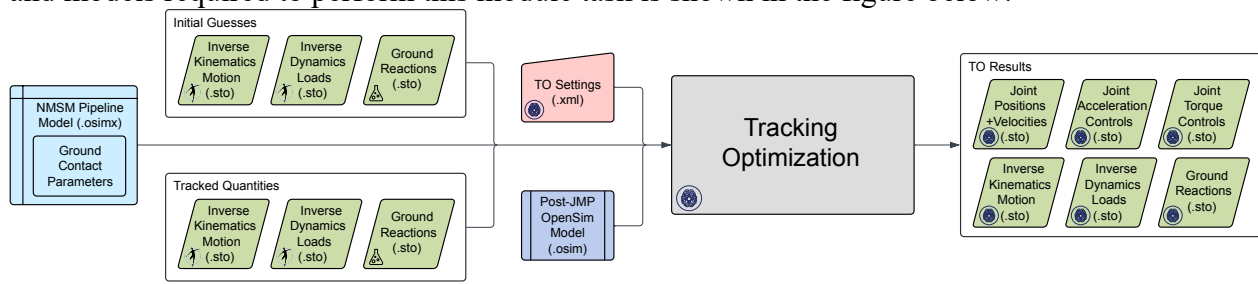

##### Step 1: Organize your input model and data files in preparation for your TO run

- Copy your OpenSim model file **Full\_Body\_Walking\_Model-Post\_JMP.osim** and your NMSM Pipeline model **Full\_Body\_Walking\_Model-Post\_GCP.osimx** file to your **TO** data folder.
- Create a folder called **inputData** within your **TO** data folder.
- Copy the three cropped gait data files created from your **Inverse Dynamics** folder to your **inputData** folder.
- Make the following changes within your **inputData** folder:
  - Move **Trial12\_Gait\_IK\_results\_filtered\_cropped.sto** to the **IKData** folder and rename it **Trial12\_Gait.sto**.
  - Move **Trial12\_Gait\_ID\_results\_filtered\_cropped.sto** to the **IDData** folder and rename it **Trial12\_Gait.sto**.
  - Move **Trial12\_Gait\_force\_filtered\_updated\_cropped.sto** to the **GRFData** folder and rename it **Trial12\_Gait.sto**.

All three of these gait data files have the same file name by design. The folder in which each file is located tells the NMSM Pipeline what each data file is.

#### Step 2: Create your Tracking Optimization settings file

• Load the post-JMP model `Full_Body_Walking_Model-Post_JMP.osim` into the OpenSim GUI. NMSM Pipeline tools will not be accessible in the OpenSim GUI **Tools** menu unless a model to simulate is loaded first.
• Select **Tools** ⇒ **User Plugins** ⇒ `rcnlPlugin.dll` to load the NMSM Pipeline tools into the OpenSim GUI **Tools** menu.
• Select the **Tracking Optimization** tool to set up a TO run that reproduces the inverse kinematics, inverse dynamics, and ground reaction data for the one-cycle gait motion as closely as possible while making the simulated motion dynamically consistent.
• On the **Settings** tab, make the following choices in the **Input** section:
  • Leave the pre-selected model name and path unchanged.
  • For **Osimx File:**, choose `Full_Body_Walking_Model-Post_GCP.osimx`.
  • For **Initial Guess Dir:**, choose your `inputData` directory.
  • For **Tracked Quantities Dir:**, also chose your `inputData` directory since the same data will be used for both an initial guess for the solution and the tracked data for the cost function and constraints.
  • For **Trial Prefix:**, input `Trial12_Gait`.
• Still on the **Settings** tab, make the following choice in the **Output** section:
  - For **Results Dir:**, create a new folder in your `TO` directory called `toResults`.
• Still on the **Settings** tab, make the following choice in the **Optimal Control Solver Settings File** section:
  - Click the folder icon on the **Browse..** line and select the file `gpopsSettings.xml` in your `TO` directory. This file contains default GPOPS-II optimal control solver settings that tend to work well for any **Treatment Optimization** problem.
• Still on the **Settings** tab, for the **States Coordinates List:**, select all unlocked (i.e., changeable) generalized coordinates in your OpenSim model `Full_Body_Walking_Model-Post_JMP.osim`. DO NOT select any locked coordinates for your states.
• On the **RCNL Controllers** tab, scroll down to the **RCNL Torque Controller** section, click on **Edit...**, and select the following coordinates to be controlled by torque controllers: `hip_flexion_r`, `hip_adduction_r`, `hip_rotation_r`, `knee_angle_r`, `ankle_angle_r`, `subtalar_angle_r`, `hip_flexion_l`, `hip_adduction_l`, `hip_rotation_l`, `knee_angle_l`, `ankle_angle_l`, `subtalar_angle_l`.
  Torque controllers are placed on only those joints that would be controlled by muscles if muscles were present in the model.
• On the **Cost/Constraints** tab, in the **Cost Terms** section at the top, click on **Add...** and add the cost terms and associated maximum allowable errors as outlined in the table below:

| <type> | Entities | <max_allowable_error> |
| --- | --- | --- |
| `generalized_coordinate_tracking` | `pelvis_tx pelvis_tz` | 0.02 |
| `generalized_coordinate_tracking` | `pelvis_tilt pelvis_list`<br>`pelvis_rotation`<br>`hip_flexion_r`<br>`hip_adduction_r`<br>`hip_rotation_r`<br>`knee_angle_r`<br>`ankle_angle_r`<br>`hip_flexion_l` | 0.1745 |
|  | hip_adduction_l<br>hip_rotation_l<br>knee_angle_l<br>ankle_angle_l |  |
| generalized_coordinate_tracking | subtalar_angle_r<br>mtp_angle_r<br>subtalar_angle_l<br>mtp_angle_l | 0.0873 |
| generalized_coordinate_tracking | lumbar_extension | 0.6981 |
| generalized_coordinate_tracking | lumbar_bending<br>lumbar_rotation<br>arm_flex_r arm_add_r<br>arm_rot_r elbow_flex_r<br>arm_flex_l arm_add_l<br>arm_rot_l elbow_flex_l | 0.3491 |
| generalized_speed_tracking | pelvis_tx pelvis_tz | 0.2 |
| generalized_speed_tracking | pelvis_tilt pelvis_list<br>pelvis_rotation<br>hip_flexion_r<br>hip_adduction_r<br>hip_rotation_r<br>knee_angle_r<br>ankle_angle_r<br>hip_flexion_l<br>hip_adduction_l<br>hip_rotation_l<br>knee_angle_l<br>ankle_angle_l | 1.7453 |
| generalized_speed_tracking | subtalar_angle_r<br>mtp_angle_r<br>subtalar_angle_l<br>mtp_angle_l | 0.8727 |
| generalized_speed_tracking | lumbar_extension<br>lumbar_bending<br>lumbar_rotation<br>arm_flex_r arm_add_r<br>arm_rot_r elbow_flex_r<br>arm_flex_l arm_add_l<br>arm_rot_l elbow_flex_l | 3.4907 |
| inverse_dynamics_load_tracking | hip_flexion_r_moment<br>hip_adduction_r_moment<br>hip_rotation_r_moment<br>knee_angle_r_moment<br>ankle_angle_r_moment<br>subtalar_angle_r_moment<br>hip_flexion_l_moment | 10 |
|  | hip_adduction_l_moment<br>hip_rotation_l_moment<br>knee_angle_l_moment<br>ankle_angle_l_moment<br>subtalar_angle_l_moment |  |
| external_force_tracking | ground_force_2_vx<br>ground_force_2_vy<br>ground_force_2_vz<br>ground_force_1_vx<br>ground_force_1_vy<br>ground_force_1_vz | 40 |
| external_moment_tracking | ground_moment_2_mx<br>ground_moment_2_my<br>ground_moment_2_mz<br>ground_moment_1_mx<br>ground_moment_1_my<br>ground_moment_1_mz | 10 |

The **pelvis\_tx** and **pelvis\_tz** coordinates should be tracked lightly, since you need only a small amount of error for them to keep the pelvis over a unique location on the treadmill. You should never track the **pelvis\_ty** coordinate since the pelvis height needs to adjust to allow the foot-ground contact models to equilibrate properly. All maximum allowable errors above were scaled up by a factor of two from their original values to speed up convergence. Maximum allowable errors for generalized coordinate tracking were selected so that lower body joint angles are tracked well to facilitate reproducing the correct inverse dynamics joint moments, ankle, subtalar, and toes angles are tracked very well to facilitate reproducing the correct ground reaction forces and moments, and upper body joint angles are tracked less well to balance out the dynamics. Maximum allowable errors for generalized speed tracking were generally chosen to be 10x greater than corresponding errors for generalized coordinate tracking. Inverse dynamic load tracking is applied to all joints for which an **RCNL Torque Controller** is present.

• Still on the **Cost/Constraints** tab, in the **Constraint Terms** section at the bottom, click on **Add...** and add the constraint terms outlined in the table below:

| <type> | Entities | <max_error> |
| --- | --- | --- |
| generalized_coordinate_periodicity | pelvis_tx pelvis_ty<br>pelvis_tz | 0.01 |
| generalized_coordinate_periodicity | pelvis_tilt pelvis_list<br>pelvis_rotation<br>hip_flexion_r<br>hip_adduction_r<br>hip_rotation_r<br>knee_angle_r<br>ankle_angle_r<br>subtalar_angle_r<br>mtp_angle_r<br>hip_flexion_l<br>hip_adduction_l | 0.0175 |
|  | hip_rotation_l<br>knee_angle_l<br>ankle_angle_l<br>subtalar_angle_l<br>mtp_angle_l<br>lumbar_extension<br>lumbar_bending<br>lumbar_rotation<br>arm_flex_r arm_add_r<br>arm_rot_r elbow_flex_r<br>arm_flex_l arm_add_l<br>arm_rot_l elbow_flex_l |  |
| generalized_speed_periodicity | pelvis_tx pelvis_ty<br>pelvis_tz | 0.1 |
| generalized_speed_periodicity | pelvis_tilt pelvis_list<br>pelvis_rotation<br>hip_flexion_r<br>hip_adduction_r<br>hip_rotation_r<br>knee_angle_r<br>ankle_angle_r<br>subtalar_angle_r<br>mtp_angle_r<br>hip_flexion_l<br>hip_adduction_l<br>hip_rotation_l<br>knee_angle_l<br>ankle_angle_l<br>subtalar_angle_l<br>mtp_angle_l<br>lumbar_extension<br>lumbar_bending<br>lumbar_rotation<br>arm_flex_r arm_add_r<br>arm_rot_r elbow_flex_r<br>arm_flex_l arm_add_l<br>arm_rot_l elbow_flex_l | 0.0873 |
| kinetic_consistency | hip_flexion_r_moment<br>hip_adduction_r_moment<br>hip_rotation_r_moment<br>knee_angle_r_moment<br>ankle_angle_r_moment<br>subtalar_angle_r_moment<br>hip_flexion_l_moment<br>hip_adduction_l_moment<br>hip_rotation_l_moment | 0.1 |
|  | knee_angle_1_moment<br>ankle_angle_1_moment<br>subtalar_angle_1_moment |  |
| root_segment_residual_load | pelvis_tx_force<br>pelvis_ty_force<br>pelvis_tz_force | 1 |
| root_segment_residual_load | pelvis_tilt_moment<br>pelvis_list_moment<br>pelvis_rotation_moment | 0.1 |
| external_force_periodicity | ground_force_2_vx<br>ground_force_2_vy<br>ground_force_2_vz<br>ground_force_1_vx<br>ground_force_1_vy<br>ground_force_1_vz | 5 |
| external_moment_periodicity | ground_moment_2_mx<br>ground_moment_2_my<br>ground_moment_2_mz<br>ground_moment_1_mx<br>ground_moment_1_my<br>ground_moment_1_mz | 1 |

For this problem, `<min_error>` should always be chosen to be the negative of `<max_error>`, though it need not be. Adding periodicity to the constraints makes the simulated gait cycle near-periodic, which is a required condition for performing predictive simulations of a single walking cycle in subsequent **Design Optimization** runs. The closer the specified constraints are to perfectly periodic, the longer the TO run will take to converge. A kinetic consistency constraint is required for all joints with an **RCNL Torque Controller**.

• Once you have finished configuring your TO settings file in the OpenSim GUI, **Save** your settings file in your **TO** folder and name it **TO\_Settings.xml**. Remember that if you want to read your TO settings file back into the OpenSim GUI to modify it (as opposed to simply modifying it in a text editor), you will need to use a different file name when you re-save the settings file due to a bug in the GUI implementation.

##### Step 3: Run your Tracking Optimization settings file and plot your results

Once you have created your TO settings file, run the file and generate results by following the instructions below:

• Open Matlab, load the NMSM Pipeline project if necessary, and change directories to where your TO settings file is located.
• Run the TO tool in Matlab using the settings file you just saved by inputting the following commands into Matlab:

```
>> tic
>> TrackingOptimizationTool('TO_Settings.xml')
>> toc
```

Since none of the Treatment Optimization tools are parallelized through Matlab, you do not need to start a Matlab parallel pool to avoid having parallel processing startup impact your total wall clock time.

- Plot your TO results using Matlab function

`plotTreatmentOptimizationResultsFromSettingsFile.m` as shown below:

```
>> plotTreatmentOptimizationResultsFromSettingsFile('TO_Settings.xml')
```

This function will output plots of experimental and simulated generalized coordinates, generalized speeds, inverse dynamics loads, and ground reaction forces and moments, along with plots of simulated torque controls. At the top of each subplot is a root-mean-square error showing the difference between the experimental and simulated quantity.

**Optional Exploration:** The relative weighting of the cost function and constraint terms significantly affects the speed of **Tracking Optimization** convergence. Explore changing your TO settings file in the ways shown below and note how it changes both the solution and the number of iterations required for convergence:

- Multiply the `<max_allowable_error>` by 2 for each of your cost function terms.
- Divide the `<max_allowable_error>` by 2 for each of your cost function terms.
- Multiply the `<max_error>` and `<min_error>` by 2 for each of your constraint terms.
- Divide the `<max_error>` and `<min_error>` by 2 for each of your constraint terms.

##### Deliverables

- Wall clock time (min) and number of iterations for your TO run.

Wall clock time: \_\_\_\_\_ min

Number of iterations: \_\_\_\_\_

- RMS errors for both feet in matching experimental generalized coordinates (m or deg), generalized speeds (m/s or rad/sec), inverse dynamics loads (Nm), ground reaction forces (N), and ground reaction moments (Nm) (complete the table below):

|  | Generalized Coordinate<br>(m or deg) | Generalized Speed<br>(m/s or rad/s) | Inverse Dynamics<br>Moment (Nm) |
| --- | --- | --- | --- |
| `pelvis_tx_force` |  |  |  |
| `pelvis_ty_force` |  |  |  |
| `pelvis_tz_force` |  |  |  |
| `pelvis_tilt` |  |  |  |
| `pelvis_list` |  |  |  |
| `pelvis_rotation` |  |  |  |
| `hip_flexion_r` |  |  |  |
| `hip_adduction_r` |  |  |  |
| `hip_rotation_r` |  |  |  |
| `knee_angle_r` |  |  |  |
| `ankle_angle_r` |  |  |  |
| `subtalar_angle_r` |  |  |  |
| `mtp_angle_r` |  |  |  |
| `hip_flexion_l` |  |  |  |
| `hip_adduction_l` |  |  |  |
| `hip_rotation_l` |  |  |  |
| `knee_angle_l` |  |  |  |
| `ankle_angle_l` |  |  |  |
| subtalar_angle_l |  |  |  |
| mtp_angle_l |  |  |  |
| lumbar_extension |  |  |  |
| lumbar_bending |  |  |  |
| lumbar_rotation |  |  |  |
| arm_flex_r |  |  |  |
| arm_add_r |  |  |  |
| arm_rot_r |  |  |  |
| elbow_flex_r |  |  |  |
| arm_flex_l |  |  |  |
| arm_add_l |  |  |  |
| arm_rot_l |  |  |  |
| elbow_flex_l |  |  |  |

3. RMS errors for both feet in matching experimental ground reaction forces (N) and ground reaction moments (Nm) (complete the table below):

|  | Ground Reaction<br>(N or Nm) |
| --- | --- |
| ground_force_2_vx |  |
| ground_force_2_vy |  |
| ground_force_2_vz |  |
| ground_force_1_vx |  |
| ground_force_1_vy |  |
| ground_force_1_vz |  |
| ground_moment_2_mx |  |
| ground_moment_2_my |  |
| ground_moment_2_mz |  |
| ground_moment_1_mx |  |
| ground_moment_1_my |  |
| ground_moment_1_mz |  |

4. Absolute values of the peak adduction moment for both knees in units of Nm and in units of percent bodyweight x height (subject bodyweight was 714 N and height was 1.70 m).

|  | Units of Nm | Units of %BWxHt |
| --- | --- | --- |
| knee_adduction_r_moment |  |  |
| knee_adduction_l_moment |  |  |

5. Plots of your generalized coordinate, generalized speed, inverse dynamics load, and ground reaction force and moments errors, as generated by Matlab plotting function `plotTreatmentOptimizationResultsFromSettingsFile`.
6. Your TO settings file for the TO run whose results are shown in the tables above.
7. A brief paragraph discussing what you believe to be the “largest” error in your TO results (realizing that different types of errors have different units), why you believe this error is the “largest one,” and one idea that could potentially improve it.

#### MODULE 4: VERIFICATION AND DESIGN OPTIMIZATION

In this module, you will use the NMSM Pipeline **Design Optimization** (DO) tool to design one or two personalized clinical treatments that reduce the peak adduction moment in both knees of a subject with bilateral medial compartment knee osteoarthritis. One treatment optimization problem will involve the design of personalized high tibial osteotomy surgery while the other will involve the design of personalized gait modifications. *Undergraduate students are required to complete only one of the two treatment optimization problems and may choose between the two, while graduate students are required to complete both treatment optimization problems.*

For both treatment optimization problems, you will need to perform a **Verification Optimization** (VO) followed by a **Design Optimization**. Your **Verification Optimization** will start from your **Tracking Optimization** (TO) solution to confirm that the torque controls produced by your TO run generate the correct lower body walking motion and ground reactions without tracking those quantities in the cost function. Similarly, your subsequent **Design Optimization** will start from your **Verification Optimization** solution to predict how the modeled treatment will affect the subject's walking function and knee adduction moment peaks.

The VO and DO runs for this module will be performed using the following model and data files:

• OpenSim model file: **Full\_Body\_Walking\_Model-Post\_JMP.osim**
• NMSM Pipeline model file: **Full\_Body\_Walking\_Model-Post\_GCP.osimx**
• Inverse kinematics, ground reaction, and inverse dynamics data files: found in your TO results folder **toResults**.

The general **Verification Optimization** and **Design Optimization** processes to be performed for both treatment optimization problems are described in the two module tasks below. Following this general description, specific details are provided for how these general processes should be modified to account for the unique aspects of each treatment optimization problem.

##### Module Task 1: Verification Optimization

For this module task, you will use the NMSM Pipeline **Verification Optimization** tool to verify that the lower body joint torque controls found by your **Tracking Optimization** produce the same walking motion and ground reactions as did your **Tracking Optimization** but without tracking those quantities in the optimization cost function. This “sanity check” optimization will seek to minimize changes in your **Tracking Optimization** lower body torque controls and upper body joint motions so that a dynamically consistent near-periodic walking motion is produced. Results generated by your **Tracking Optimization** will provide the starting point for your **Verification Optimization**. The interaction between tool settings, data, and models required to perform this module task is shown in the figure below:

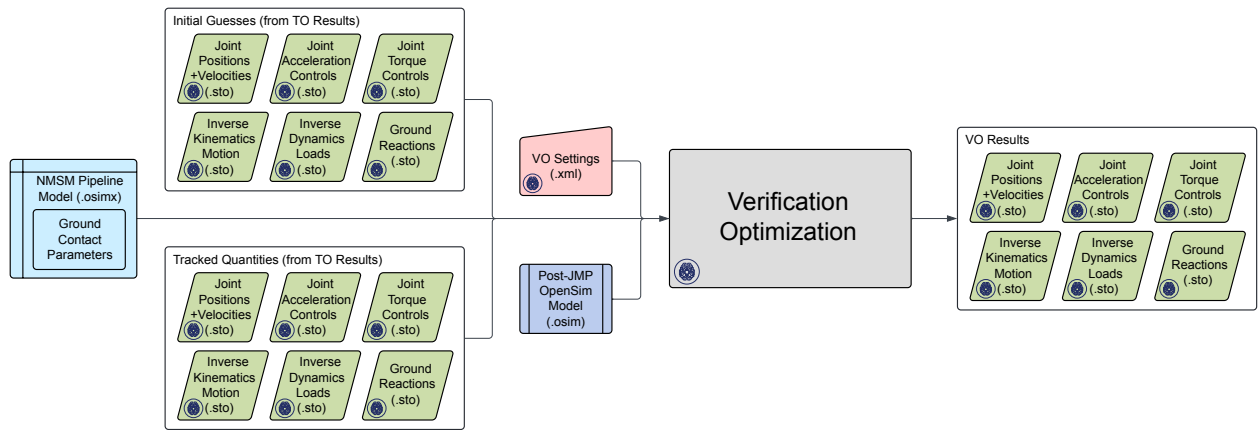

Your VO run will serve as a “dry run” for formulating the optimal control problem that you will solve in a subsequent DO run. You can think of your VO run as a DO run without the addition of the treatment design element. Normally a VO problem will be formulated primarily by removing cost and/or constraint terms from the original TO problem formulation. However, to make a VO run a “dry run” for a subsequent DO run, you will sometimes need to add one or more cost and/or constraint terms that were not present in your original TO problem formulation. When this situation arises, you will need to verify that the added cost and/or constraint terms do not affect the ability of your VO run to converge rapidly (i.e., within a few iterations) to your TO solution. If cost and/or constraint terms added to a VO problem formulation significantly increase the number of iterations required to converge, then the added cost and/or constraint terms are likely inconsistent with your base VO problem formulation containing only cost and constraint terms from your original TO run.

##### Step 1: Create your Verification Optimization settings file

• Copy your OpenSim model file **Full\_Body\_Walking\_Model-Post\_JMP.osim** and your NMSM Pipeline model file **Full\_Body\_Walking\_Model-Post\_GCP.osimx** to your **VO** data folder.
• Create a VO settings file that is a modified version of your previous TO settings file.
• If you do not want to follow the process outlined below to create your VO settings file using the OpenSim GUI, you can instead copy your TO settings file to a new settings file named **VO\_Settings.xml**, open **VO\_Settings.xml** in a text editor, change **TrackingOptimizationTool** to **VerificationOptimizationTool** at the top and bottom of the file, and make any additional changes to the VO settings file as described below.
• If you choose to use the OpenSim GUI to create your VO settings file, start by loading your post-JMP model **Full\_Body\_Walking\_Model-Post\_JMP.osim** into the OpenSim GUI. NMSM Pipeline tools will not be accessible in the OpenSim GUI **Tools** menu unless a model is loaded first.
• Select **Tools** ⇒ **User Plugins** ⇒ **rcnlPlugin.dll** to load the NMSM Pipeline tools into the OpenSim GUI **Tools** menu.
• Select the **Verification Optimization** tool to set up a VO run that tracks the lower body torque controls and upper body joint motions from your TO run while making the simulated motion dynamically consistent.
• On the **Settings** tab, make the following choices in the **Input** section:
  • Leave the pre-selected model name and path unchanged.

• For **Osimx File:**, choose `Full_Body_Walking_Model-Post_GCP.osimx`.
• For **Initial Guess Dir:**, choose your `toResults` directory.
• For **Tracked Quantities Dir:**, also chose your `toResults` directory since the same data will be used for both an initial guess for the solution and the tracked data for the cost function and constraints.
• For **Trial Prefix:**, input `Trial12_Gait`.
• Still on the **Settings** tab, make the following choice in the **Output** section:
  - For **Results Dir:**, create a new folder in your `VO` directory called `voResults`.
• Still on the **Settings** tab, make the following choice in the **Optimal Control Solver Settings File** section:
  - Click the folder icon on the **Browse..** line and select the file `gpopsSettings.xml` in your `VO` directory. This file contains default GPOPS-II optimal control solver settings that tend to work well for any **Treatment Optimization** problem.
• Still on the **Settings** tab, for the **States Coordinates List:**, select all moveable generalized coordinates in your OpenSim model `Full_Body_Walking_Model-Post_JMP.osim`.
• On the **RCNL Controllers** tab, scroll down to the **RCNL Torque Controller** section, click on **Edit...**, and select the following coordinates to be controlled by torque controllers:
  `hip_flexion_r, hip_adduction_r, hip_rotation_r, knee_angle_r, ankle_angle_r, subtalar_angle_r, hip_flexion_l, hip_adduction_l, hip_rotation_l, knee_angle_l, ankle_angle_l, subtalar_angle_l`.
  Torque controllers are placed on only those joints that would be controlled by muscles if muscles were present in the model.
• On the **Cost/Constraints** tab, in the **Cost Terms** section at the top, click on **Add...** and add the cost terms and associated maximum allowable errors as outlined in the table below:

| <type> | Entities | <max_allowable_error> |
| --- | --- | --- |
| `controller_tracking` (was `inverse_dynamics_load_tracking` in Tracking Optimization) | `hip_flexion_r_moment`<br>`hip_adduction_r_moment`<br>`hip_rotation_r_moment`<br>`knee_angle_r_moment`<br>`ankle_angle_r_moment`<br>`subtalar_angle_r_moment`<br>`hip_flexion_l_moment`<br>`hip_adduction_l_moment`<br>`hip_rotation_l_moment`<br>`knee_angle_l_moment`<br>`ankle_angle_l_moment`<br>`subtalar_angle_l_moment` | 10 |
| `generalized_coordinate_tracking` | `mtp_angle_r`<br>`mtp_angle_l` | 0.0873 |
| `generalized_coordinate_tracking` | `lumbar_extension` | 0.6981 |
| `generalized_coordinate_tracking` | `lumbar_bending`<br>`lumbar_rotation`<br>`arm_flex_r arm_add_r`<br>`arm_rot_r elbow_flex_r`<br>`arm_flex_l arm_add_l`<br>`arm_rot_l elbow_flex_l` | 0.3491 |

These cost function terms come directly from your TO settings file except that every joint in the model is now controlled by either 1) controller tracking (lower body joints except for mtp joints), 2) generalized coordinate tracking (upper body joints plus mtp joints), or 3) no tracking (pelvis coordinates). In general, unlike for TO problem formulations, VO problem formulations generally have no joints controlled by both a cost function term and a constraint term (though this general rule can sometimes be violated to improve convergence for a subsequent DO run).

• Still on the **Cost/Constraints** tab, in the **Constraint Terms** section at the bottom, click on **Add...** and add the constraint terms outlined in the table below:

| <type> | Entities | <max_error> |
| --- | --- | --- |
| generalized_coordinate_periodicity | pelvis_tx pelvis_ty<br>pelvis_tz | 0.01 |
| generalized_coordinate_periodicity | pelvis_tilt pelvis_list<br>pelvis_rotation<br>hip_flexion_r<br>hip_adduction_r<br>hip_rotation_r<br>knee_angle_r<br>ankle_angle_r<br>subtalar_angle_r<br>mtp_angle_r<br>hip_flexion_l<br>hip_adduction_l<br>hip_rotation_l<br>knee_angle_l<br>ankle_angle_l<br>subtalar_angle_l<br>mtp_angle_l<br>lumbar_extension<br>lumbar_bending<br>lumbar_rotation<br>arm_flex_r arm_add_r<br>arm_rot_r elbow_flex_r<br>arm_flex_l arm_add_l<br>arm_rot_l elbow_flex_l | 0.0175 |
| generalized_speed_periodicity | pelvis_tx pelvis_ty<br>pelvis_tz | 0.1 |
| generalized_speed_periodicity | pelvis_tilt pelvis_list<br>pelvis_rotation<br>hip_flexion_r<br>hip_adduction_r<br>hip_rotation_r<br>knee_angle_r<br>ankle_angle_r<br>subtalar_angle_r<br>mtp_angle_r | 0.0873 |
|  | hip_flexion_l<br>hip_adduction_l<br>hip_rotation_l<br>knee_angle_l<br>ankle_angle_l<br>subtalar_angle_l<br>mtp_angle_l<br>lumbar_extension<br>lumbar_bending<br>lumbar_rotation<br>arm_flex_r arm_add_r<br>arm_rot_r elbow_flex_r<br>arm_flex_l arm_add_l<br>arm_rot_l elbow_flex_l |  |
| kinetic_consistency | hip_flexion_r_moment<br>hip_adduction_r_moment<br>hip_rotation_r_moment<br>knee_angle_r_moment<br>ankle_angle_r_moment<br>subtalar_angle_r_moment<br>hip_flexion_l_moment<br>hip_adduction_l_moment<br>hip_rotation_l_moment<br>knee_angle_l_moment<br>ankle_angle_l_moment<br>subtalar_angle_l_moment | 0.1 |
| root_segment_residual_load | pelvis_tx_force<br>pelvis_ty_force<br>pelvis_tz_force | 1 |
| root_segment_residual_load | pelvis_tilt_moment<br>pelvis_list_moment<br>pelvis_rotation_moment | 0.1 |
| external_force_periodicity | ground_force_2_vx<br>ground_force_2_vy<br>ground_force_2_vz<br>ground_force_1_vx<br>ground_force_1_vy<br>ground_force_1_vz | 5 |
| external_moment_periodicity | ground_moment_2_mx<br>ground_moment_2_my<br>ground_moment_2_mz<br>ground_moment_1_mx<br>ground_moment_1_my<br>ground_moment_1_mz | 1 |

These constraint terms come directly from your TO settings file with no changes whatsoever.

• Once you have finished configuring your VO settings file in the OpenSim GUI, **Save** your

settings file in your **VO** folder and name it **VO\_Settings.xml**. Remember that if you want to read your VO settings file back into the OpenSim GUI to modify it (as opposed to simply modifying it in a text editor), you will need to use a different file name when you re-save the settings file due to a bug in the GUI implementation.

#### Step 2: Run your Verification Optimization settings file and plot your results

Once you have created your VO settings file, run the file and generate results by following the instructions below:

• Open Matlab, load the NMSM Pipeline project if necessary, and change directories to where your VO settings file is located.
• Run the VO tool in Matlab using the settings file you just saved by inputting the following commands into Matlab:

```
>> tic
>> VerificationOptimizationTool('VO_Settings.xml')
>> toc
```

Since none of the Treatment Optimization tools are parallelized through Matlab, you do not need to start a Matlab parallel pool to avoid having parallel processing startup impact your total wall clock time.

• If your VO problem is formulated well, your VO run should converge in just a few iterations. If your VO run requires more than 10 iterations to converge, then something is wrong with your VO problem formulation, and you should review your VO settings file to try to diagnose the source of the problem. A common problem is to have a constraint term that is redundant with or conflicts with a cost function term. For example, you could have a periodicity constraint on a quantity that you are tracking in the cost function, but the tracked quantity is less periodic than required by the periodicity constraint.
• Plot your VO results using Matlab function `plotTreatmentOptimizationResultsFromSettingsFile.m` as shown below:
  ```
  >> plotTreatmentOptimizationResultsFromSettingsFile('VO_Settings.xml')
  ```
  This function will output plots of your TO and VO generalized coordinates, generalized speeds, inverse dynamics loads, ground reaction forces and moments, and simulated torque controls. At the top of each subplot is a root-mean-square (RMS) error showing the difference between TO and VO quantities.
• For all plotted quantities, verify that your VO results are visually identical to your TO results with extremely low RMS errors.

#### Module Task 2: Design Optimization

Following generation of a successful **Verification Optimization**, you will use the NMSM Pipeline **Design Optimization** tool to design a personalized treatment that reduces the subject's adduction moment peaks in both knees to a desired target value associated with an excellent long-term clinical outcome (Bryan *et al.*, 1997). For either treatment optimization problem, results generated by your **Verification Optimization** will provide the starting point for your subsequent **Design Optimization**. The interaction between tool settings, data, and models required to perform this module task is shown in the figure below:

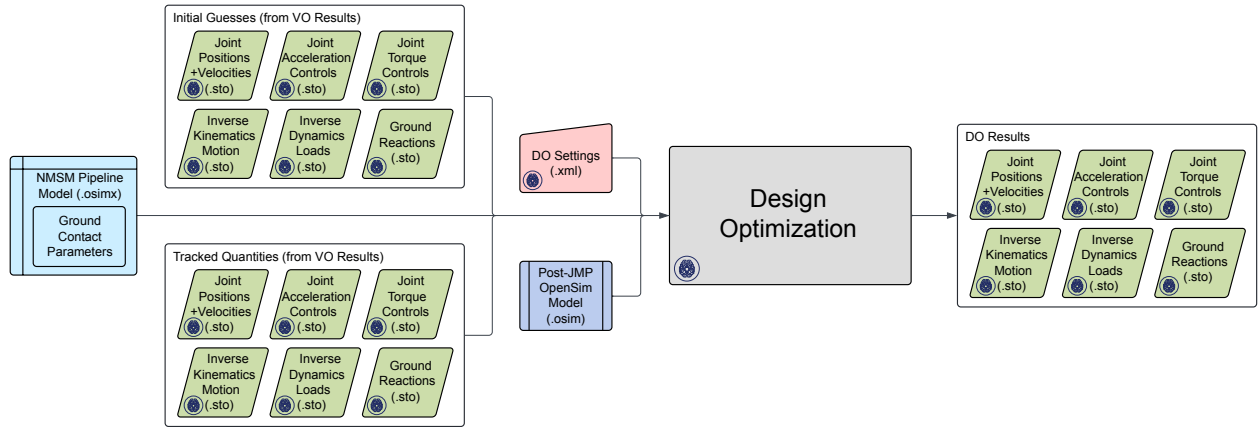

##### Step 1: Create your Design Optimization settings file

• Copy your OpenSim model file `Full_Body_Walking_Model-Post_JMP.osim` and your NMSM Pipeline model `Full_Body_Walking_Model-Post_GCP.osimx` file to your DO data folder.
• Create a DO settings file by modifying your VO settings file in a text editor (Note: You could also create this settings file using the OpenSim GUI, but creating it in a text editor is the fastest and easiest approach). Copy your VO settings file to a new settings file named `DO_Settings.xml`, open `DO_Settings.xml` in a text editor, and make the following changes:
  - Change `VerificationOptimizationTool` to `DesignOptimizationTool` at the top and bottom of the file.
  - Change the results directory to `doResults` in your DO folder.
  - Change the initial guess and tracked quantities directories to `voResults` within your VO folder.

##### Step 2: Run your Design Optimization settings file and plot your results

Once you have created your DO settings file, run the file and generate results by following the instructions below:

• Open Matlab, load the NMSM Pipeline project if necessary, and change directories to where your DO settings file is located.
• Run the DO tool in Matlab using the settings file you just saved by inputting the following commands into Matlab:

```
>> tic
>> DesignOptimizationTool('DO_Settings.xml')
>> toc
```

Since none of the Treatment Optimization tools are parallelized through Matlab, you do not need to start a Matlab parallel pool to avoid having parallel processing startup impact your total wall clock time.

• If your DO problem is formulated well, your DO run should converge in less than 250 iterations.
• Plot your DO results using Matlab function

`plotTreatmentOptimizationResultsFromSettingsFile.m` as shown below:

```
>> plotTreatmentOptimizationResultsFromSettingsFile('DO_Settings.xml')
```

This function will output plots of your VO and DO generalized coordinates, generalized speeds, inverse dynamics loads, ground reaction forces and moments, and simulated torque controls. At the top of each subplot is an RMS error showing the difference between VO and DO quantities.

• Record the predicted absolute value of the peak knee adduction moment for both knees. Recall that your treatment design goal is to achieve a peak knee adduction of 2.5 %BW\*Ht in both knees. For the subject being modeled (BW = 714 N, Ht = 1.70 m), this target value is equivalent to 30.3 Nm.

##### Step 3: Animate your predicted motion in OpenSim

• Load your OpenSim model `Full_Body_Walking_Model-Post_JMP.osim` into the OpenSim GUI twice.
• For the first model, select **File** ⇒ **Load Motion...** and select the IK results file located in your `voResults\IKData` folder.
• For the second model, make the bones a different color, select **File** ⇒ **Load Motion...**, and select the IK results file located in your `doResults\IKData` folder.
• Sync the two motions and then animate them together at a slow **Speed** (e.g., 0.25). Note how much each knee in the predicted DO motion is medialized relative to the corresponding VO motion.

Given this general process for performing **Verification Optimization** and **Design Optimization**, problem-specific modifications required to solve each treatment optimization problem are described below:

#### Treatment Optimization Problem 1: Design of Personalized High Tibial Osteotomy Surgery

##### Step 1: Run a Verification Optimization

• Use the VO settings file described above with no modifications.
• Confirm that convergence to your TO solution occurs within just a few iterations.

##### Step 2: Add an HTO surgical correction to both tibias of your OpenSim model

• Copy your OpenSim model `Full_Body_Walking_Model-Post_JMP.osim` to a new model called `Full_Body_Walking_Model-Post_HTO.osim`.
• Open `Full_Body_Walking_Model-Post_HTO.osim` in a text editor and make the following changes:
  - At the top of the file, change the model name to `Full_Body_Walking_Model-Post_HTO`.
  - Search for coordinate name `knee_adduction_r` and change the default value to -0.0524 (3 deg), keeping the coordinate locked. This change sets the fixed knee adduction angle for the right tibia to 3 deg, emulating a 3 deg surgical correction in frontal plane leg alignment.
  - Repeat for coordinate name `knee_adduction_l`.
  - Save the model with these changes.

##### Step 3: Run a Design Optimization using your post-HTO OpenSim model

• Use the DO settings file described above except with the input model file changed to your post-HTO OpenSim model.
• Confirm that your DO run converges within roughly 250 or fewer iterations.

**Step 4: Repeat using surgical corrections of 6 deg and 9 deg for both tibias**

• Run a **Design Optimization** using surgical corrections of 3 deg for both tibias.
• Repeat using surgical corrections of 6 deg for both tibias.
• Repeat using surgical corrections of 9 deg for both tibias.
• Record the predicted peak adduction moment for each knee in the deliverables table below.

**Step 5: Perform a quadratic fit to estimate the optimal surgical correction for each tibia**

**Step 6: Perform a final Design Optimization using the optimal surgical corrections**

**Deliverables**

1. Wall clock time (min) and number of iterations for your final DO run.
  Wall clock time: \_\_\_\_\_ min
  Number of iterations: \_\_\_\_\_
2. Peak adduction moment for both knees using surgical corrections of 3, 6, and 9 deg for both tibias (complete the table below):

| Surgical Correction<br>(deg) | Peak Adduction Moment (Nm) |  |
| --- | --- | --- |
|  | Right Knee | Left Knee |
| 3 |  |  |
| 6 |  |  |
| 9 |  |  |

3. Optimal surgical correction for each tibia based on a quadratic fit to the data in the table above:
  Optimal surgical correction for right tibia = \_\_\_\_\_ deg
  Optimal surgical correction for left tibia = \_\_\_\_\_ deg
4. Peak adduction moment for both knees predicted by your final DO run that used the post-HTO OpenSim model with optimal surgical correction for each tibia:
  Predicted peak adduction moment for right knee = \_\_\_\_\_ Nm
  Predicted peak adduction moment for left knee = \_\_\_\_\_ Nm
5. Plots of your inverse dynamics loads from your final DO run as generated by Matlab plotting function `plotTreatmentOptimizationResultsFromSettingsFile`.
6. The DO settings file for your final DO run that used the post-HTO OpenSim model with optimal surgical correction for each tibia.
7. A brief paragraph discussing how well the quadratic fit worked for estimating the optimal amount of surgical correction needed for each tibia to achieve post-HTO surgery knee adduction moment peaks of 30.3 Nm for each knee.

**Treatment Optimization Problem 2: Design of Personalized Gait Modifications**

**Step 1: Run a modified Verification Optimization**

• To match the optimization problem formulation used in (Fregly *et al.*, 2007), make the following additions to the VO settings file described above:
  - Add cost terms and associated maximum allowable errors as outlined in the table below:

| <type> | Entities | <max_allowable_error> |
| --- | --- | --- |
| marker_position_tracking<br>(x and z axes only) | R_Heel<br>R_Toe | 0.0025 |
|  | L_Heel<br>L_Toe |  |
| body_orientation_tracking<br>(x, y, and z axes using a yxz sequence) | calcn_r<br>calcn_l<br>torso | 0.0175 |

These cost function terms make both feet follow the experimentally measured motion with respect to the lab coordinate system and make the torso follow the experimentally measured orientation with respect to the lab coordinate system. We do not track the foot marker y positions since we need to give the ground contact model freedom to equilibrate the model dynamics in the vertical direction.

- Add constraint terms and associated maximum/minimum errors as outlined in the table below:

| <type> | Entities | <max_error> |
| --- | --- | --- |
| marker_position_deviation<br>(x, y, and z axes) | R_Heel<br>R_Toe<br>L_Heel<br>L_Toe | 0.0025 |
| body_orientation_deviation<br>(x, y, and z axes using a yxz sequence) | calcn_r<br>calcn_l<br>torso | 0.0175 |

These new constraint terms force the optimizer to find a solution where foot motion and torso orientation remain close to their experimental trajectories. Using a similar cost term and constraint term for marker positions and body orientations seems redundant but helps the optimizer converge more rapidly to a solution that matches the experimental foot motion and torso orientation closely.

• Confirm that convergence to your TO solution occurs within just a few iterations, despite the addition of these new cost function and constraint terms.

##### Step 2: Run a Design Optimization using your post-JMP OpenSim model

• Use the DO settings file described above (i.e., taken directly from your VO settings file) except with the addition of the following cost function term:

| <type> | Entities | <max_allowable_error> |
| --- | --- | --- |
| inverse_dynamics_load_minimization | knee_adduction_r_moment<br>knee_adduction_l_moment | 10 |

• Confirm that your DO run converges within roughly 250 or fewer iterations.

##### Step 3: Repeat using max allowable errors of 15 and 20 Nm for both knee adduction moments

• Run a **Design Optimization** using maximum allowable errors of 10 Nm for both knee adduction moments.
• Repeat using maximum allowable errors of 15 Nm for both knee adduction moments.
• Repeat using maximum allowable errors of 20 Nm for both knee adduction moments.
• Record the predicted peak adduction moment for each knee in the deliverables table below.

##### Step 4: Perform a quadratic fit to estimate the optimal max allowable error for each knee

##### Step 5: Perform a final Design Optimization using the optimal max allowable error values

**Deliverables**

1. Wall clock time (min) and number of iterations for your final DO run.
  Wall clock time: \_\_\_\_\_ min
  Number of iterations: \_\_\_\_\_
2. Peak adduction moment for both knees using max allowable errors of 10, 15, and 20 Nm for both knee adduction moments (complete the table below):

| Max Allowable Error<br>(Nm) | Peak Adduction Moment (Nm) |  |
| --- | --- | --- |
|  | Right Knee | Left Knee |
| 10 |  |  |
| 15 |  |  |
| 20 |  |  |

3. Optimal max allowable error for each knee adduction moment based on a quadratic fit to the data in the table above:
  Optimal max allowable error for right knee adduction moment = \_\_\_\_\_ Nm
  Optimal max allowable error for left knee adduction moment = \_\_\_\_\_ Nm
4. Peak adduction moment for both knees predicted by your final DO run that used the optimal max allowable error for each knee adduction moment:
  Predicted peak adduction moment for right knee = \_\_\_\_\_ Nm
  Predicted peak adduction moment for left knee = \_\_\_\_\_ Nm
5. Plots of your inverse dynamics loads from your final DO run as generated by Matlab plotting function `plotTreatmentOptimizationResultsFromSettingsFile`.
6. The DO settings file for your final DO run that used the optimal max allowable error for each knee adduction moment.
7. A brief paragraph discussing how well the quadratic fit worked for estimating the optimal max allowable error for each knee adduction moment to achieve post-gait modification knee adduction moment peaks of 30.3 Nm for each knee.

### Supplementary Material B: Tutorial 2

#### SYNERGY-DRIVEN NEUROMUSCULOSKELETAL MODEL TREATMENT OPTIMIZATION

##### NMSM Pipeline Advanced Tutorial 2

**Tutorial Developers:** Robert Salati, Geng Li, and B.J. Fregly, Rice Computational Neuromechanics Lab, Rice University

###### Simulation Project Materials

The materials for this tutorial can be downloaded from SimTK at https://simtk.org/projects/nmsm in the “NMSM Advanced Tutorials” section.

###### Simulation Project Overview

The goal of this simulation project is to teach you the Neuromusculoskeletal Modeling (NMSM) Pipeline’s computational treatment design process for clinical applications (Fregly 2021; Hammond et al. 2025) where muscles and neural control models are needed. This project will develop a treatment for a high-functioning stroke survivor using computationally design functional electrical stimulation (FES) patterns. Stroke survivors often have damaged neural pathways which impacts their ability to walk. As such, for this project, we will calibrate muscle-tendon parameters for the subject to accurately model their forces and then personalize a neural control model for the subject. Information from this neural control model will then be used to inform the treatment design process to design an optimal FES pattern (Cheung et al. 2019; Levine et al. 2025). This project will seek to design an FES pattern that compensates for a weakened synergy in the subject’s affected limb and consequently improving their propulsive and braking ground reaction impulses to be more symmetric between limbs.

The five tools that you will use within the NMSM Pipeline are indicated in the table below, along with the abbreviations used to reference each tool and required supporting OpenSim tools:

| NMSM Pipeline Toolset | NMSM Pipeline Tool | Used |
| --- | --- | --- |
| Model Personalization | Joint Model Personalization (JMP) | Provided |
|  | Muscle-tendon Model Personalization (MTP)<br>(with OpenSim Muscle Analysis tool) | ✓ |
|  | Neural Control Model Personalization (NCP) | ✓ |
|  | Ground Contact Model Personalization (GCP) | Provided |
| Treatment Optimization | Tracking Optimization (TO) | ✓ |
|  | Verification Optimization (VO) | ✓ |
|  | Design Optimization (DO) | ✓ |

For the **Model Personalization** toolset, you will only be using **Muscle-tendon Personalization (MTP)** and **Neural Control Personalization (NCP)**. While a synergy-driven treatment optimization does require **Joint Model Personalization (JMP)** and **Ground Contact Personalization (GCP)**, these steps have already been done for you. All 3 Treatment Optimization tools will be used in this project, but the model will now be controlled by synergy-driven muscle-tendon actuators.

This simulation project is broken down into four modules – Muscle-tendon Personalization, Neural Control Personalization, Tracking Optimization, and Verification/Design Optimization. For each module, instructions are provided to walk you through all the necessary steps. To ensure that poor results for one module do not affect your ability to complete subsequent modules, final results will be provided for each module to use as a starting point for subsequent modules. At the end of each module, you will be asked questions aimed at guiding you through interpreting your results.

To run each required OpenSim or NMSM Pipeline tool, you will generate an initial xml settings file using the appropriate tool selection within the OpenSim GUI Tools menu. Once you have generated an initial tool settings file in the OpenSim GUI, you can edit the settings file for subsequent tool runs either within the OpenSim GUI or using a text editor. Runs for OpenSim tools will be performed through the OpenSim GUI, while runs for NMSM Pipeline tools will be performed in Matlab.

This project will use treadmill gait data from a single high-functioning hemiparetic stroke survivor. For this individual, their right side is the paretic/affected side, and their left side is the less-affected side. All experimental joint angle, joint load, ground reaction, and electromyography (EMG) data have been provided to you inside the **InputData** folder but will need to be processed before being used for this project. The initial data provided to you is summarized below:

• **Trial10\_IKResults.mot** – Joint angles from inverse kinematics (IK)
• **Trial10\_forces\_ec\_reordered\_filtered.mot** – Ground reaction forces (GRF) and moments with a shifted electrical center from GCP
• **Trial10\_IDResults.sto** – Joint loads from inverse dynamics (ID)
• **Trial10\_emg\_processed.sto** – Processed EMG data

IK joint angles were solved using video-based marker motion capture (Vicon Corporation, Oxford, United Kingdom) with a post-JMP full-body OpenSim model. Six degree-of-freedom (DOF) ground reaction data were collected from a Bertec split-belt instrumented treadmill (Bertec Corporation, Columbus, OH, United States) with belts tied to the same speed. The experimental data from each trial above have already been converted so that all length data are in units of meters, all moment data are in units of Newton-meters, and all data are reported using coordinate axes consistent with OpenSim model conventions (i.e., +X is directed anteriorly, +Y is directed superiorly, and +Z is directed out to the right).

A scaled, post-JMP OpenSim model file is provided to you as well. The file name is **UF\_Subject\_4\_Scaled\_JMP.osim**.

#### MODULE 1: MUSCLE-TENDON PERSONALIZATION

In this module, you will do data processing and use the NMSM Pipeline Muscle-tendon Personalization (MTP) tool. MTP is classified as “EMG-driven” modeling. In other words, EMG data are an input to the model, rather than an unknown quantity. This is different from models based around Static Optimization, where muscle activity is not known. MTP’s primary inputs are EMG data, kinematic data, and joint moment data. MTP will use EMG data in combination with muscle kinematics to generate individual muscle forces that apply moments about the model joints. These muscle joint moments need to match the experimental joint moment data. The main optimization in MTP calibrates Hill-type muscle model parameters so that the moments produced by muscles closely match experimental joint moments.

The Hill-type muscle model used in this project calculates muscle force according to the equation:

$$F = F_o^M \cdot \left[ a(e(t - d)) \cdot f_l(\tilde{l}^M(t)) \cdot f_v(\tilde{v}^M(t)) + f_p(\tilde{l}^M(t)) \right] \cos(\alpha)$$

Where $F_o^M$ is the maximum isometric force of the muscle, $a$ is the muscle’s activation which is a function of processed EMG data $e$, $t$ is time, $d$ is an electromechanical time delay, $\tilde{l}^M$ and $\tilde{v}^M$ are the normalized muscle fiber length and velocity respectively, and $\alpha$ is the muscle-tendon pennation angle. Tendons are assumed to be rigid, so $\tilde{l}^M$ and $\tilde{v}^M$ are calculated using the following equations:

$$\tilde{l}^M = \frac{l^{MT} - l_s^T}{l_o^M \cos(\alpha)}$$

$$\tilde{v}^M = \frac{v^{MT}}{10 \cdot l_o^M}$$

where $l^{MT}$ is the muscle-tendon length, $l_s^T$ is the tendon slack length, and $l_o^M$ is the optimal muscle fiber length.

Of these variables, MTP calibrates $l_o^M$, $l_s^T$, and $d$. Additionally, MTP calibrates an activation time constant $\tau_{act}$ and activation non-linearity factor $c_3$. Finally, MTP calibrates muscle excitation scale factors to change the maximum amplitude of each EMG signal. For more details on these parameters, refer to (Meyer et al. 2017) and (Zajac 1989).

The MTP design variables are:

• **Max isometric force** – The maximum force a muscle can produce under isometric conditions.
• **Optimal muscle fiber length** – The fiber length at which a muscle can produce the maximum force. At the optimal fiber length, the normalized fiber length is 1.
• **Tendon slack length** – The “resting spring length” of the tendon attaching to the muscle. We are using rigid tendons, so the tendon slack length is equal to the length of the tendon.
• **Electromechanical delay** – A time delay between the measurement of the muscle excitation using an EMG sensor and when the excitation truly occurred.
• **Activation time constant** – A time constant governing the dynamics between a muscle excitation and a muscle activation.

• **Activation non-linearity constant** – A constant governing how non-linear the relationship between muscle excitation and muscle activation is.

The forces produced by a Hill-type muscle model have two components – active force and passive force. Active force is the force produced by contracting a muscle, and is a function of muscle activation $a$, normalized muscle fiber length $\tilde{l}^M(t)$, and normalized muscle fiber velocity $\tilde{v}^M(t)$. Passive force is the force produced by a muscle’s intrinsic structure when the muscle is stretched too much. Passive force is only a function of normalized muscle fiber length $\tilde{l}^M(t)$, and is only present when a muscle’s normalized fiber length exceeds 1. More intuitively, if you are doing stretches before going on a run, the tension you feel in the muscles is due to passive force – you are physically stretching the muscles, and the muscle structure resists that stretching.

MTP has 2 additional sub-tools that work along with the main body of the optimization. The first subtool is Muscle-tendon Length Initialization (MTLI). MTLI’s purpose is to calibrate muscle-tendon length parameters and max isometric force values to make a better initial guess for MTP, and it always runs before MTP. To isolate muscle-tendon length parameters in the Hill-type force equation, MTLI only uses passive muscle forces. While passive force generated by individual muscles is difficult to measure, we can measure joint moments caused by these passive forces. This was done in a study that used a dynamometer to measure joint moments while moving joints through their range of motion with no muscle activation (Silder et al. 2007). MTLI calibrates muscle-tendon length parameters such that these passive joint moments are closely matched.

The second sub-tool is synergy extrapolation (SynX), which runs in parallel to MTP (Ao et al. 2022). SynX uses muscle synergies from the measured muscle excitations to estimate the excitations for muscles that weren’t measured. This is important because even if a muscle isn’t measured, we need to know how much force the muscle is output. If we chose to only include muscles that were measured, but could only measure 8 muscles per leg, the force from each of those muscles will be vastly overestimated. With SynX, we can simultaneously estimate the excitations for missing muscles and calibrate muscle parameters. The minimum number of EMG signals needed to get a good calibration with SynX is 8 EMG signals per leg (Ao and Fregly 2024)

The personalization will be performed using IK joint angles, ID joint loads, and processed EMG data. The interaction between tool settings, data, and models required to perform this module is shown in the figure below:

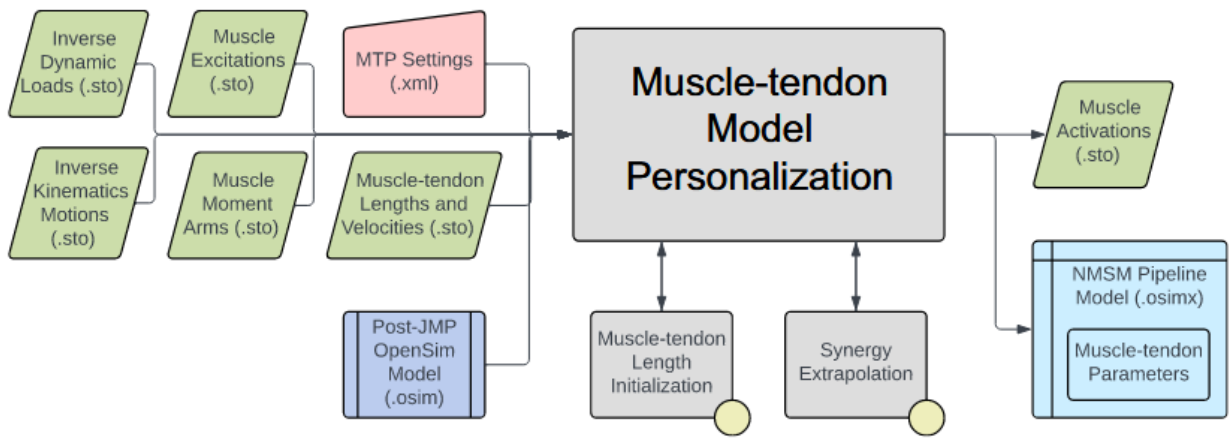

Data for IK joint motions, ID joint loads, and processed muscle excitations have already been provided. A post-JMP OpenSim model has also been provided. Muscle moment arms, and muscle-tendon lengths and velocities will be calculated in the first task for this module. In the next task, data will be processed to be in the correct format to use the MTP tool and the remainder of the NMSM Pipeline. Finally, you will conclude this module by running MTP.

##### Module Task 1: Muscle Analysis

A key step towards using MTP is to calculate muscular kinematic quantities such as moment arms, and muscle-tendon lengths. These quantities can be calculated using OpenSim's Muscle Analysis (MA) tool. The MA tool is a subtool of OpenSim's Analysis tool. A short guide to the Analysis tool can be found here.

The MA tool uses joint motions to calculate muscle-tendon lengths and muscle-moment arms about every joint using geometry calculations. Every modeled muscle has two attachment points on separate bodies such that they cross one or more joints and can apply moments about those joints. Anatomical muscles also have complex interactions with the bodies they're attached to. The OpenSim model used in this project represents these interactions using wrapping surfaces that define how muscles wrap around bones. These wrapping surfaces can be visualized in the OpenSim GUI by expanding the *bodies* tab under the model and then expanding individual bodies. The MA tool provides a quick and easy way to perform geometry calculations with these wrapping surfaces.

###### Step 1: Muscle analysis for gait

In this step, you will conduct MA on gait motion to calculate moment arms and muscle-tendon lengths throughout the gait cycle. These quantities will then be used in the main body of MTP to calculate muscle forces and corresponding joint moments. The input data files needed for this task step are in the **inputData** directory.

1. Load the model **UF\_Subject\_4\_Scaled\_JMP.osim** into the OpenSim GUI
2. Open the *Analyze tool*
3. Load the motion file **Trial10\_IKResults.mot**

4. Change the prefix to **Trial10**
  a. This changes the prefix added at the beginning of every file created by MA. This is important for file organization
5. Change the output directory to **MuscleAnalysis\MADData**
6. Under the *Analyses tab*, add a MuscleAnalysis task.
7. Edit the MuscleAnalysis task and ensure that the tool will compute moment arms for all muscles and all lower body coordinates.
8. Run the MA tool.
9. Verify that the tool creates files inside **MuscleAnalysis\MADData** that all have the prefix “Trial10”.
10. **Note:** There is a bug with MA in which the gastrocnemius muscles have moment arms about the hip. To work around this, copy and paste the MA files inside input data to your new MA folder. These files should replace some of the files you just calculated.

#### Step 2: Muscle analysis for passive moment data

The next MA run will be done on passive joint moment data. MTP uses passive moment data inside Muscle-tendon Length Initialization (MTLI) to generate a good initial guess for the main MTP optimization.

MTLI uses published passive joint moment data. As described earlier, joints are moved through their full range of motion with other joints fixed at constant values. Joint moments are then measured using a dynamometer as the joint moves through its range of motion. These data are described below:

• **Thelen\_AnklePassive\_01** – Ankle passive moments with the fixed at knee at 0°
• **Thelen\_AnklePassive\_02** – Ankle passive moments with the fixed at knee at 15°
• **Thelen\_AnklePassive\_03** – Ankle passive moments with the fixed at knee at 60°
• **Thelen\_AnklePassive\_04** – Ankle passive moments with the fixed at knee at 110°
• **Thelen\_HipPassive\_01** – Hip passive moments with the knee fixed at 15°
• **Thelen\_HipPassive\_02** – Hip passive moments with the knee fixed at 60°
• **Thelen\_HipPassive\_03** – Hip passive moments with the knee fixed at 90°
• **Thelen\_HipPassive\_04** – Hip passive moments with the knee fixed at 110°
• **Thelen\_KneePassive\_01** – Knee passive moments with the ankle fixed at 20°
• **Thelen\_KneePassive\_02** – Knee passive moments with the ankle fixed at -15°
• **Thelen\_KneePassive\_03** – Knee passive moments with the hip fixed at 45°
• **Thelen\_KneePassive\_04** – Knee passive moments with the ankle fixed at 20° and the hip fixed at -15°

Inside the **MuscleAnalysis** folder, there are two folders **thelen\_r** and **thelen\_l** that contain passive joint moment data for then right and left legs respectively. They contain folders **IKData** and **IDData**, where **IDData** contains the passive joint moments, and **IKData** contains the corresponding joint angles. You will run MA for all 12 motion trials in **IKData**, for both legs.

1. Open the Matlab file **passiveTrialMuscleAnalysis.m**
2. Edit the variable **thelenFolder** to say either “thelen\_r” or “thelen\_l”
3. Edit the variable **modelFileName** to be the full path to your **.osim** model file.
4. Run the Matlab script.

5. Repeat for the other leg.

These steps should create a folder called **MAData** inside each of *thelen\_r* or *thelen\_l*, and subfolders for each trial with muscle analysis files inside each folder. If these are not there, verify that all your file paths and names are correct.

##### Module Task 2: Data preprocessing

The next step in this module will be to process your data to get it into the correct format for the remainder of the NMSM Pipeline. This process will contain several parts, all of which are contained in the NMSM Pipeline's preprocessing tool:

1. **Process EMG data:** Starting from the raw EMG file, the script high pass filters, demeans, rectifies, and low pass filters the EMG signals. Next, any remaining negative EMG values are set to zero. EMG signals are then offset so that the minimum value of each signal is 0. Finally, EMG signals are normalized so that the max value of each signal is 1.
2. **Create muscle-tendon velocities:** The script low pass filters the muscle-tendon lengths created by Muscle Analysis, splines the filtered data using GCV splines, and then differentiates the filtered data.
3. **Crop and resample data:** A user input to preprocessing is a set of time pairs for which the data should be cropped to. One file is created for each time pair specified by the variable `trialTimePairs`. The data are then splined and resampled to have 101 time points per trial.
4. **Lowpass filter data:** All data are low pass filtered using the cutoff frequency specified by `inputSettings.cutoffFrequency`.

Data preprocessing is contained entirely in the script `Preprocessing\preprocessing.m`. The EMG data has already been processed, so those corresponding lines of code are commented out.

1. Open `Preprocessing\preprocessing.m`.
2. Edit `lines 12-17` with the paths to your data files.
3. We want to crop the data to one gait cycle (right heelstrike to right heelstrike). You must find the start time and the end time of the right heel strikes. To do this, open the file `Preprocessing\plotInputGroundReactions.m` and find the time of the first and second right heel strike (rounded <u>down</u> to the nearest 0.01s. ie if there is force starting at 0.98 seconds, round down to 0.97s).
4. Edit `startTime` and `endTime` to be the times you found above.
  a. This variable specifies the time ranges we want to crop the data into. This time pair isolates one single gait cycle that we will use for the rest of our modeling.
5. Explore the `preprocessed` directory that was created in your `Preprocessing` folder. This folder contains 5 folders for EMG data, IK data, ID data, GRF data, and MA data.
  a. <u>Keep note of where this folder is. You will be using it for the rest of the project.</u>
  b. <u>Do not change the name of any folders or files inside `preprocessed`. This tool outputs data in a format that the NMSM Pipeline expects from here on out. Changing any of these file or folder names will result in errors.</u>
6. Visualize your preprocessing results with the function `plotPreprocessing.m`
7. Verify that all data look good (ie minimal noise, no sharp discontinuities, other data quality issues)

##### Module Task 3: Muscle-tendon Personalization

Now that you have preprocessed data, you can move onto using the NMSM Pipeline's **Muscle-tendon Personalization (MTP)** tool. This tool will make use of the preprocessed data you just created, so there is no more work to be done for input data.

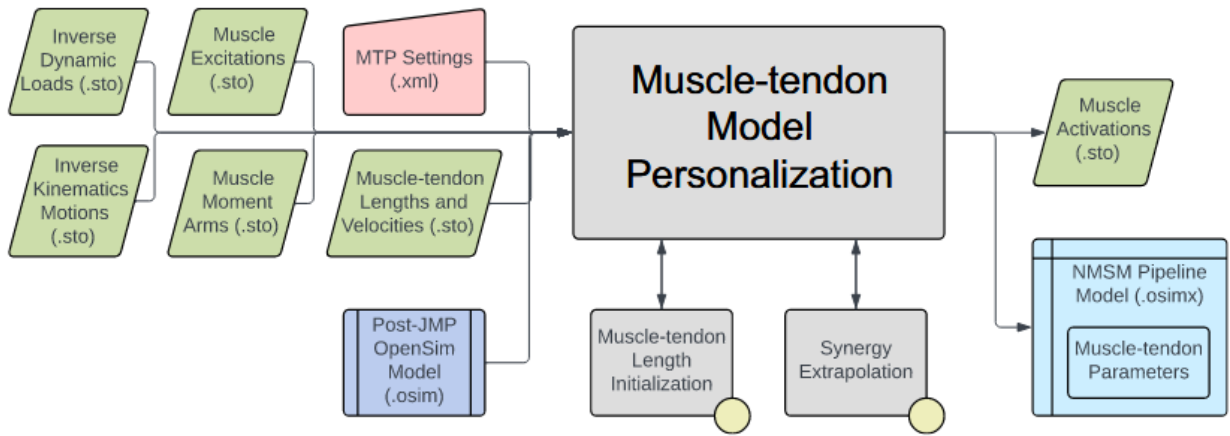

MTP will track this processed experimental data to fit Hill-type muscle model parameters for our model such that the joint moments generated by muscle forces match ID joint moments.

For this subproject, you will iterate through MTP solutions, changing cost term parameters until you arrive to a physiologically realistic solution.

#### Step 1) Explore muscle groups

1. Open the OpenSim model `UF_Subject_4_Scaled_JMP.osim` in the OpenSim GUI.
2. Under the *Forces* tab on the model, explore the muscles available.
3. Take note of the extra groups added.
  a. These are added for organization so that MTP/NCP knows which model muscles to group together in the optimization.
  b. The four important groups are:
    i. **<u>Activation Muscle Groups</u>** – Muscles that we would expect to have similar activation profiles (ie lateral hamstrings; BFSH and BFLH will have similar activations to each other). These groups all have `ActivationGroup` in their name.
    ii. **<u>Normalized Fiber Length Muscle Groups</u>** – Muscles that we would expect to have similar normalized fiber lengths. These groups all have `NormalizedFiberLengthGroup` in their name.
    iii. **<u>Collected EMG Muscle Groups</u>** – Muscle groups that we **do have** experimental EMG data for. Unlike the other groups, these must have the same name as the respective EMG channel name your EMG data file. They cannot be named `[muscle]_collectedEmgMuscleGroup` unless you change the corresponding names in your EMG data file
      `Preprocessing\preprocessed\EMGData\gait_1.sto`
    iv. **<u>Missing EMG Muscle Groups</u>** – Muscle groups that we **do not have** experimental EMG data for. These groups all have `MissingEMGChannelGroup` in their name.

The activation muscle groups and normalized fiber length muscle groups are organized based on how muscles are grouped anatomically. For example, we assume that your vastus muscles (three muscles that are a part of your quadriceps) will all have similar normalized fiber lengths and activation profiles because they serve similar functions. The collected and missing EMG muscle groups vary based on the dataset you are using, and which muscles have EMG data available.

The muscle groups for this project are premade for you but it is important to study them and understand why certain muscles are grouped together.

#### Step 2) Create your base settings files

You will need to create different settings files for the right and left leg. The reason for separating the legs is because SynX does not support multiple synergy sets. For this project, we are using 2 synergy sets – right leg and left leg. Therefore, MTP will need to be split up between the left and right leg. We will start with the right leg.

1. Load your premade post-JMP model `UF_Subject_4_Scaled_JMP.osim` into the OpenSim GUI
2. Open the Muscle-tendon Personalization Tool GUI
3. Set the *Input Osimx File* to be `GroundContactPersonalization\gcpResults\UF_Subject_4_Scaled_JMP_gcp.osimx`
4. Set the *Input Data Directory* to be `Preprocessing\preprocessed`
5. Set the *Output Results Directory* to be `MuscleTendonPersonalization\MTPResultsRightV1`
6. Set your *coordinate list* to be (`hip_flexion_r`, `hip_adduction_r`, `hip_rotation_r`, `knee_angle_r`, `ankle_angle_r`, `subtalar_angle_r`)
7. Set your *Activation Muscle Groups* to be all right leg activation groups
  a. Tip: You can use the *Filter by* box at the top of the selection window to search for only right leg activation groups
8. Set your *Normalized Fiber Length Muscle Groups* to be all right leg fiber length groups
9. Set your *Missing EMG Muscle Groups* to be all right leg missing EMG groups
10. Set your *Collected EMG Muscle Groups* to be all right leg EMG channels (Look at `Preprocessing\preprocessed\EMGData\gait_1.sto`)
11. Enable Muscle Tendon Length Initialization
12. Set the *Passive Data Input Directory* to be `MuscleAnalysis\thelen_r`
13. Enable Muscle Tendon Synergy Extrapolation with **6 synergies**
14. Save this settings file.
15. **Repeat the above steps for the left side of the body, changing all necessary fields.**
  a. You will need to change the Input Osimx File to be the file that is output from the right side MTP run. This file does not exist yet, so just copy and paste the following text into the field: `MTPResultsRightV1\UF_Subject_4_Scaled_JMP_gcp_mtp.osimx`
16. Open `MuscleTendonPersonalization\runMTP.m`, and edit the settings files name and run the script.
  a. It's important that you run the right leg first because we use the output osimx file from the right leg as the input osimx file for the left leg.
  b. <u>MTP does not use any information from the input osimx file in the optimization.</u> The purpose of the input osimx file is so that MTP can concatenate left leg results to the right leg osimx file, keeping everything in the same file.

Now that your first MTP run is complete, the next step is to iterate through settings files to fine tune a solution. To fine tune an MTP solution, there are generally 4 parameters you should change:

1. **Minimum and maximum allowable normalized fiber lengths** – These terms dictate the range that we allow the normalized fiber length to fall in, and how heavily we punish normalized fiber lengths outside that range.

2. **Experimental quantity tracking terms** – These terms affect how closely the optimization tracks joint moments.
  a. **passive\_joint\_moment** in MTLI
    i. How closely passive joint moments should be tracked in MTLI
  b. **measured\_inverse\_dynamics\_joint\_moment** in SynX
    i. How closely <u>all</u> joint moments in the model should track experimental joint moments
    ii. Reduce max allowable error to prioritize SynX more.
  c. **inverse\_dynamics\_joint\_moment** in MTP
    i. How closely joint moments produced by muscles with experimental EMG data should track experimental joint moments.
    ii. Reduce max allowable error to use experimentally measured muscles more.
3. **Max allowable errors and error centers for muscle group-based cost terms** – These terms punish deviations between muscles in a group.
  a. **grouped\_normalized\_muscle\_fiber\_length**
  b. **grouped\_emg\_scale\_factor**
  c. **grouped\_electromechanical\_delay**
4. **Regularization terms** – These terms punish deviations in muscle-tendon parameters away from the error center and ensure the solution is unique.
  a. **optimal\_muscle\_fiber\_length**
  b. **tendon\_slack\_length**
  c. **passive\_muscle\_force**
  d. **activation\_time\_constant**
  e. **activation\_nonlinearity\_constant**
  f. **emg\_scale\_factor**
  g. **muscle\_excitation\_penalty**

Some common problems in an MTP solution and corresponding solutions are:

1. **Poor joint moment tracking quality** (RMSE > 10Nm)
  a. Decrease moment tracking max allowable error
2. **Large muscle activations** (Max activation > 0.7 during gait)
  a. Decrease max allowable error/error center for **emg\_scale\_factor** or **muscle\_excitation\_penalty**.
3. **Large passive muscle forces** (Max passive force > 100)
  a. Decrease max allowable error on **maximum\_normalized\_muscle\_fiber\_length**, or **passive\_muscle\_force**
4. **Large difference between muscle activations and muscle excitations**
  h. Decrease max allowable error on **activation\_nonlinearity\_constant**, or **activation\_time\_constant**
5. **Large difference in activations or normalized fiber lengths between muscles in a group**
  a. Decrease max allowable error on **grouped\_normalized\_muscle\_fiber\_length**, **grouped\_emg\_scale\_factor**, or **grouped\_electromechanical\_delay**
6. **Under/overuse of unmeasured muscles**

a. Change max allowable error on **extrapolated\_muscle\_activation** and **residual\_muscle\_activation**

##### Step 3) Iterate through MTP solutions

###### Iteration 1:

The first problem we will address is the large passive forces, especially in the calf and quad muscles. We will address this problem in a few ways. First, we will punish normalized fiber lengths falling outside of the optimal range of 0.8-1.0. Next, we will choose to optimize absolute length changes for optimal fiber length and tendon slack length (units of meters) instead of scale factors.

1. Create copies of your settings files and rename them.
2. For your new settings files, remember to update your `<results_directory>` fields, and your `<input_osimx_file>` field for the left settings file.
3. Passive forces are caused by normalized fiber lengths exceeding a value of 1, so we can first reduce the max allowable error on the maximum normalized fiber lengths inside MTLI to further punish normalized fiber lengths above 1.
  a. Inside `<MuscleTendonLengthInitialization>`, lower `<max_allowable_error>` of `maximum_normalized_muscle_fiber_length` to 0.01
4. We will also increase the minimum normalized fiber length to avoid the normalized fiber length dropping too low.
  a. Inside `<MuscleTendonLengthInitialization>`:
    i. Increase `<min_normalized_muscle_fiber_length>` to 0.8
    ii. Lower `minimum_normalized_muscle_fiber_length` `<max_allowable_error>` to 0.01
5. To avoid over restricting the optimization, we can reduce the moment tracking value for MTLI as well.
  a. Inside `<MuscleTendonLengthInitialization>`, increase `passive_joint_moment` `<max_allowable_error>` to 15
6. Next, we can change the optimization to optimize length parameters using absolute lengths instead of relative lengths. The previous optimization changed the scale factors of the optimal fiber length and tendon slack length. We will change this to optimize absolute changes in length, instead of scale factors
  a. Inside `<MuscleTendonLengthInitialization>`:
    i. Copy and paste the following:
      `<optimize_absolute_length_changes>true</optimize_absolute_length_changes>`
    ii. Change `optimal_muscle_fiber_length` max allowable error to 0.005 and the error center to 0
    iii. Change `tendon_slack_length` max allowable error to 0.005 and the error center to 0
7. Finally, inside the main optimization, we can reduce the max allowable error on the passive force cost term.
  a. Inside `<MTPTaskList>`, lower `<max_allowable_error>` of `passive_muscle_force` to 10.

After running iteration 1, all passive forces should be below 50N. All normalized fiber lengths should be mostly between the range of 0.6-1.0. Some normalized fiber lengths may be a little bit above 1, but there should be a noticeable improvement over the initial MTP run.

#### **Iteration 2:**

The next observation is that certain muscles' activations are significantly different from their excitations, beyond a simple time shift (psoas & recfem for example). Also, some muscles have very large muscle activations which is not realistic during gait (Left leg psoas and iliacus). We will first address these problems by lowering the max allowable error on the activation non-linearity constant. This will punish high non-linearities that are causing muscle activations to have much higher amplitude than the corresponding muscle excitations. Next, we will tighten max allowable errors on muscle-excitations to punish the large muscle excitations.

1. Create copies of your settings files and rename them
2. For your new settings files, remember to update your `<results_directory>` fields, and your `<input_osimx_file>` field for the left settings file.
3. Change the following cost terms in both settings files:

##### **i. Muscle\_excitation\_penalty**

i. Max allowable error: **0.3**
ii. Error center: **0.3**
iii. This change will punish excitations from getting too high

##### **j. Emg\_scale\_factor**

i. Max allowable error: **0.3**
ii. Error center: **0.3**
iii. This change will also punish excitations from getting too high, but works with EMG scale factors rather than excitation magnitude.

##### **k. Activation\_nonlinearity\_constant**

i. Max allowable error: **0.005**
ii. This change will move muscle activations closer to their excitations to fix muscles like the psoas.

##### Iteration 3:

This last iteration will address issues with SynX activations. Specifically, the right side SynX muscles (sartorius, piriformis, and quadratus femoris) are overactivated on the right leg and underactivated on the left leg. We will change this by tightening the max allowable errors on SynX. This iteration will also use different max allowable errors for the left and right sides (Why might this be?)

1. Create copies of your settings files and rename them
2. For your new settings files, remember to update your `<results_directory>` fields, and your `<input_osimx_file>` field for the left settings file.
3. Right side:
  a. Inside `<MTPSynergyExtrapolation>`:
    i. Change `measured_inverse_dynamics_joint_moment` max allowable error to 2
    ii. Change `extrapolated_muscle_activation` max allowable error to 0.1
  b. Inside `<MTPTask>`:
    i. Change `inverse_dynamics_joint_moment` max allowable error to 3
4. Left side:
  a. Inside `<MTPSynergyExtrapolation>`:
    i. Change `measured_inverse_dynamics_joint_moment` max allowable error to 1
    ii. Change `extrapolated_muscle_activation` max allowable error to 1
  b. Inside `<MTPTask>`:
    i. Change `inverse_dynamics_joint_moment` max allowable error to 3

#### Deliverables:

Inside one PDF file:

1. Submit your final plots of joint moments, normalized fiber lengths, muscle activations, and muscle passive forces for both legs.
2. In 1-2 sentences, explain how the MTP optimization works.
3. In 2-3 sentences each, explain how MTLI and SynX work, and why MTP needs them.
4. Summarize each MTP iteration you did. Information to include is:
  a. What was the goal of the iteration?
  b. What muscles were you aiming to fix?
  c. Did the iteration achieve its goal?
5. Briefly explain why the final settings files were different between the legs (Hint: what problem were we trying to solve in iteration 3? Was it the same problem for both legs?)
6. Compare the muscle activations between the left and right sides. Are there any muscles that have significantly different activations on the left and right sides?
7. Do you think that the final solution you got for MTP is good enough to move forwards? Why or why not?
8. If you were to do one more MTP iteration, what would your goal be? What muscles are you trying to fix, and what cost terms would you change to achieve this goal?
9. When selecting the joint moments to match during MTP, we didn't use the knee adduction moment or the mtp (toes) moment. Why didn't we use these moments?
  a. Toes: (Hint: Do we have joint moments for the toes? Why or why not? If not, how could we solve for toes joint moments?)
  b. Knee adduction: (Hint: What generates the knee adduction moment? Remember that the net moment about a joint is given by:
    $$M^{net} = M^{contact} + M^{muscle} + M^{ligament}$$
    )
10. In one paragraph explain why MTP is important for muscle-driven predictive simulations.

#### MODULE 2: NEURAL CONTROL PERSONALIZATION

In this module, you will use the NMSM Pipeline Neural Control Personalization (NCP) tool. NCP fits a set of muscle synergies to your data that best matches experimental muscle activations and reproduces experimental joint moments. Like MTP, NCP also calculates muscle joint moments and tries to track experimental joint moments. The design variables in NCP however, are the activations themselves. Specifically, NCP calculates a set of synergy activations and synergy vectors that create muscle activations that produce the correct muscle forces. It does this with a cost function that tracks muscle activations from MTP, and experimental joint moments. Because NCP is calculating muscle forces, it is best to use the muscle model calibrated in MTP. The desired output of NCP is a set of muscle synergies that are “functional”, meaning when the muscle synergies are given to the model, they reproduce joint moments and can make the model walk.

It is important that synergies are functional for Treatment Optimization. In the next module, the forces generated by synergy-driven muscles will drive the motion of our gait model. We will then be able to change the muscle synergies to achieve our desired functional outcome for this stroke subject.

This calibration will be performed using ID joint moments, muscle moment arms, muscle-tendon lengths and velocities, and muscle activations (not excitations) generated by MTP. The interaction between tool settings, data, and models required to perform this module is shown in the figure below:

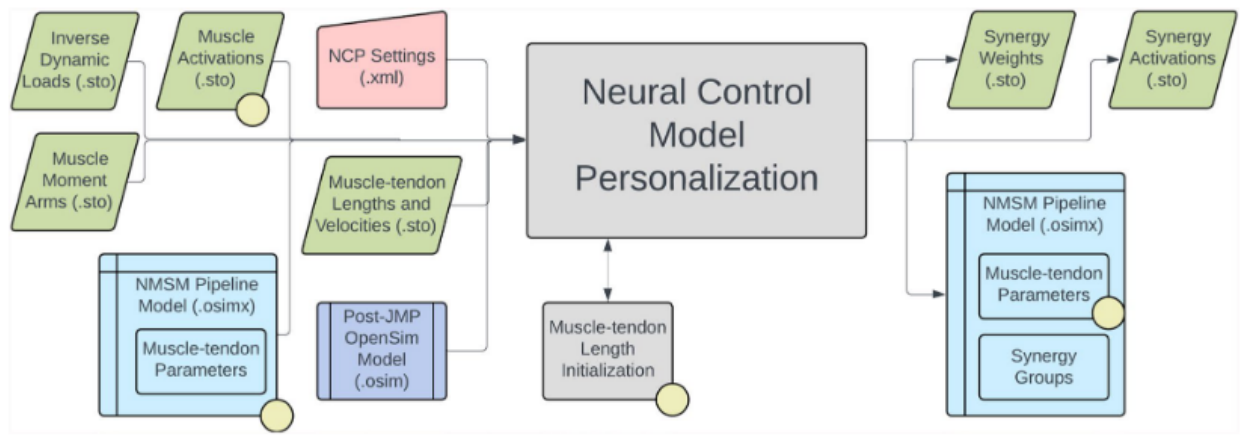

Because we ran MTP first, NCP will use the muscle model and muscle activations calibrated by MTP. The first task of this subproject will be to format your MTP results in the correct form for NCP. The second task will be to estimate how many synergies NCP should use. The third task will be to create an NCP settings file and fit your muscle synergies. The last task is to thoroughly analyze NCP results to diagnose this subject’s impairment.

##### Module Task 1) Format NCP inputs directory

Unlike MTP, NCP does support multiple synergy sets, so we can combine our MTP results into a single NCP run. Therefore, the first task in this subproject is to combine your left and right MTP results.

There is a premade MATLAB script `NeuralControlPersonalization\CombineMTPResults.m` that combines results for you. Open this script and input the paths to your right and left MTP results directories.

The script will concatenate the output joint moments and muscle activations from both runs, and write the concatenated file to a new directory `NeuralControlPersonalization\mtpResultsCombined`. It will also copy the osimx file from your left leg MTP results directory to `mtpResultsCombined`.

##### Module Task 2) Estimate how many synergies to use in NCP.

At their core, muscle synergies are a mathematical decomposition of muscle activations. The more synergies you have, the better that decomposition will represent the original data. There are diminishing returns as more synergies are added, however. For a dataset with 16 EMG signals per leg, 5-6 synergies can typically represent more than 95% of the original data. Adding more synergies beyond that runs the risk of overfitting your experimental data which can make predicting new motions more difficult. To avoid overfitting our data, it is important to use a minimum set of synergies that represent the experimental data well.

The measure we use to quantify how well muscle synergies reproduce the original activations is called *percentage of variance accounted for* (%VAF), represented by the equation

$$\%VAF = \left( 1 - \frac{\sum (x - x')^2}{\sum x} \right) \cdot 100$$

Where $x$ and $x'$ are the experimental EMG data and the reconstructed EMG data, respectively. For neuromusculoskeletal modeling, we aim for our synergies to have 90-95% VAF.

For this task, we will conduct a VAF analysis on our EMG data to estimate how many synergies we should use in our NCP run. Open the script `SynergyVafAnalysis.m` and fill out the required variables. This function will conduct a synergy analysis for $k=4, 5, 6$ synergies, plot the reconstructed muscle activations for visualization, and output %VAF values for each number of synergies.

Choose the minimum number of synergies that produces 95% VAF for the left and right side. You will use this number of synergies for NCP.

##### Module Task 3) Run NCP

Now that you have an initial guess for how many synergies to use, we can create an NCP settings file.

###### Step 1) Initial NCP Settings file

1. Load your post-JMP model **UF\_Subject\_4\_Scaled\_JMP.osim** into the OpenSim GUI
2. Open the Neural Control Personalization Tool GUI
3. Set the *Input Osimx File* to be the Osimx file generated by your final left MTP run.
4. Set the *Input Data Directory* to **Preprocessing\preprocessed**
5. Set the *Output Results Directory* to be **NeuralControlPersonalization\NCPResultsNoBilateral**
6. Set your *coordinate list* to be all lower limb coordinates used in your previous MTP runs (left and right)
7. Set your *Activation Muscle Groups* to be every activation muscle group.
8. Set your *Normalized Fiber Length Groups* to be every normalized fiber length group.
9. Keep MTLI disabled.
10. Set your *MTP Results Directory* to be your combined MTP results directory.
11. Set your *Synergy Set* to be **right\_leg** and **left\_leg** with the number of synergies you decided on. Make sure both legs have the same number of synergies.
  a. Is it a good assumption for a stroke subject that both legs have the same number of synergies?
12. Save this settings file.
13. Open your settings file in a text editor.
14. Open your settings file in a text editor and edit the following cost terms inside of your `<RCNLCostTermSet>`:
  a. **moment\_tracking** – max allowable error = 2
  b. **activation\_tracking** – max allowable error = 0.015
15. Copy and paste the following into your `<RCNLCostTermSet>`:

```
<RCNLCostTerm>
  <type>grouped_activations</type>
  <is_enabled>true</is_enabled>
  <max_allowable_error>0.05</max_allowable_error>
</RCNLCostTerm>
```

16. Open the Matlab script **NeuralControlPersonalization\runNCP**.
17. This first NCP run may take ~30 minutes to run on the Duncan Hall computers.

#### Step 2) Experiment with bilateral symmetry

Your initial NCP run treated each leg as independent from the other. This might not be a realistic assumption, however. During a coordinated motion such as gait, your entire body is working as a system, so it is very likely that the synergies in your left and right leg are not acting independently from one another. This can be changed by including another cost term in NCP that adds coordination between the synergy sets in the body. This is called “bilateral symmetry”.

When decomposing a set of muscle activations into muscle synergies, we are representing muscle activations by a set of time varying commands that control activation timing, and time-invariant weights that control coordination between muscles. There is literature suggesting that the time varying commands are mostly based inside the brain, while the time invariant synergy vector weights are based inside the spinal cord. When using bilateral symmetry, you are imposing that the synergy vectors between each leg must be the same. We are assuming that because the stroke only affects the brain, the synergy vector weights in the spinal cord are unaffected. The synergy activations can still be different from each other. In other words, the timing in activations between the legs can be different, but the way in which muscles are coordinated with each other should stay the same between legs.

1. Create a copy of your NCP settings file
2. Change your *Output Results Directory* to **NCPResultsBilateral5Synergies**.
3. Open your settings file in a text editor
4. Set `<enforce_bilateral_symmetry>` to **true**
5. This settings file will take longer to run than the previous settings file. As such, you are **not** required to run this settings file. Some premade results are inside **NCPResultsBilateral**.
6. Comment out the line to run NCP, plot the premade results, and answer the following questions.

#### Deliverables:

Inside one PDF file, submit all plots outputted by both NCP runs, and answer the following questions:

1. Fill out the following table with your %VAF values from task 2:

| Num Synergies | %VAF Right | %VAF Left |
| --- | --- | --- |
| 4 |  |  |
| 5 |  |  |
| 6 |  |  |

2. In 1-2 sentences, explain how the NCP optimization works.
3. Briefly explain what muscle synergies are, and why we might want to use them in neuromusculoskeletal simulations instead of individual muscle activations.
4. Why do we not want to use more synergies than required to reach 95% VAF?
5. Answer the following questions about bilateral symmetry:
  a. In 1-2 sentences, explain what bilateral symmetry is and why it's important.
  b. What %VAF did you get when running NCP with and without bilateral symmetry? Is this lower or higher than without bilateral symmetry? Does this result make sense?
  c. How did using bilateral symmetry affect your tracking quality of joint moments or muscle activations?
  d. Do you think that using bilateral symmetry is reasonable for a stroke subject?

The next group of questions will focus on analyzing the NCP results. You should use the NCP results with no bilateral symmetry that you created, and the premade results with bilateral symmetry.

First, we will analyze the solution without bilateral symmetry. To identify an impairment between the legs, it is important that we can compare synergies between both legs. This requires matching right leg and left leg synergies to each other. For the solution without bilateral symmetry, we can match synergies by identifying which synergies contain similar muscles on the left and right leg. Mathematically, this can be done by taking dot products between the left and right leg synergy vectors.

Inside **NeuralControlPersonalization**, open **CompareSynergyVectors.m**, fill out your NCP results directory for the run without bilateral symmetry, and click run. This script does two tasks:

1) Normalize synergy vectors – In order to ensure NCP's solution is unique, synergy vectors are normalized such that each vector has a maximum weight of 1. Because we want to take dot products to compare similarity between synergy vectors, we will renormalize all synergy vectors to have a magnitude of 1. In other words, we are turning the synergy vectors into unit vectors.
2) Make a grid of dot products between all combinations of left and right synergy vectors – We dot right\_leg synergy vector 1 with left\_leg synergy vectors 1, 2, 3, and so on, for all right leg synergies.

This script outputs a table with the dot products between all synergy vectors. A higher dot product corresponds to a more similar synergy activation.

6. Using the table created in `CompareSynergyVectors.m`, fill out the following table with the matching left leg synergy for each right leg synergy. Type N/A for boxes which you didn't use that many synergies.

| Right Leg Synergy | Left Leg Synergy | Dot Product Value |
| --- | --- | --- |
| 1 |  |  |
| 2 |  |  |
| 3 |  |  |
| 4 |  |  |
| 5 |  |  |
| 6 |  |  |

7. How similar would you say your left and right leg synergies are? Do you think it is reasonable to compare these matched synergies to each other? Why or why not?
8. For the solution **with** bilateral symmetry, fill out the following table comparing the left and right synergy activations to each other. The magnitude of a synergy activation is its maximum value. The number of peaks is the number of prominent maximums in the activation. The location of the peaks should be reported as a percentage of the gait cycle, and a comma separated list if there are multiple peaks.

|  | Right |  |  | Left |  |  |  |
| --- | --- | --- | --- | --- | --- | --- | --- |
| Synergy Number | Magnitude | Number of peaks | Location of peaks | Magnitude | Number of peaks | Location of peaks | Similar shape? (yes/no) |
| 1 |  |  |  |  |  |  |  |
| 2 |  |  |  |  |  |  |  |
| 3 |  |  |  |  |  |  |  |
| 4 |  |  |  |  |  |  |  |
| 5 |  |  |  |  |  |  |  |

**A note on similar shapes:** To decide if two synergies have a similar shape, you should be comparing the number of peaks and the locations of the peaks between the legs. Keep in mind that the left side will have a 50% gait cycle time shift from the right side. For example, if right leg synergy 1 has two peaks at 10% and 40% of the gait cycle, a “similar” left leg synergy 1 would have two peaks at 60% and 90% of the gait cycle. If both the right and left leg synergies have peaks at the same percentage in gait cycle, they are **not** similar to each other.

9. For the solution **with** bilateral symmetry, fill out the following table listing the primary muscles in each synergy. If a muscle has a synergy vector greater than 0.03 for a given synergy, write it in the table below.

| Synergy Number | Primary Muscles in Synergy Vector |
| --- | --- |
| 1 |  |
| 2 |  |
| 3 |  |
| 4 |  |
| 5 |  |

10. Using the given table detailing muscle groups, fill out the below table with what muscle groups are most represented in each synergy.

| **Synergy Number** | **Primary Muscle Groups in Synergy Vector** |
| --- | --- |
| 1 |  |
| 2 |  |
| 3 |  |
| 4 |  |
| 5 |  |

Primary muscle groups and their corresponding muscles are listed below:

| **Muscle Group** | **Muscles in Group** |
| --- | --- |
| Hip Adductors | Addbrev, Addlong, Addmag |
| Hip Abductors | Glmax, Glmed, Glmin, |
| Hip Rotators | Piri, Tfl, Gem, Quadfem |
| Hip Flexors | Iliacus, Psoas, Recfem, Sart |
| Hip Extensors | Bflh, Glmax, Semimem, Semiten |
| Knee Flexors | Bflh, Bfsh, Semimem, Semiten, Gaslat, Gasmed |
| Knee Extensors | Recfem, Vasint, Vaslat, Vasmed |
| Ankle Plantarflexors | Gaslat, Gasmed, Soleus |
| Ankle Dorsiflexors | Tibant, Edl, Ehl |
| Ankle Inverters/Everters | Edl, Ehl, Perbrev, Perlong, Tibpost, Fdl, Fhl |

11. Muscle synergies are often assigned a biomechanical function based on when in the gait cycle they are most active (Ting et al. 2015). Some common tasks assigned to synergies are:

a. Weight acceptance
b. Propulsion
c. Flexion
d. Leg swing

The figure below shows the approximate shape of these types of synergies.

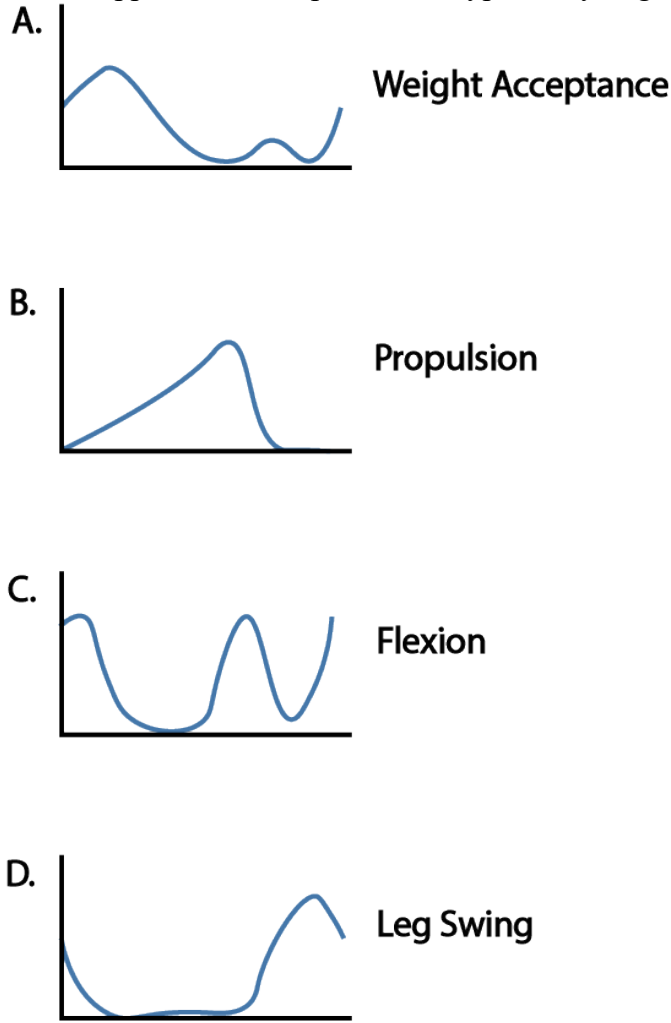

For the synergies with bilateral symmetry, fill out the following table with the function of each synergy. Reference the shapes of the synergy commands in the above figure. It is possible that multiple synergies serve the same function, or perhaps a mix of multiple functions.

| Synergy Number | Synergy Function |
| --- | --- |
| 1 |  |
| 2 |  |
| 3 |  |
| 4 |  |
| 5 |  |

12. With this subject, we observe *impaired* synergies, and *compensatory* synergies. Impaired synergies are on the affected side (right side in this case) and have a significantly lower activation than the matching synergy on the non-affected side. Compensatory synergies can be on either side, and are characterized by a synergy activation that is moderately larger than the matching synergy on the other leg.
a. Which right leg synergy (or synergies) are impaired? What function do these synergies serve? What muscle groups are in these synergies?
b. Which right or left leg synergies are compensatory? What functions do these synergies serve?

#### MODULE 3: SYNERGY-DRIVEN TRACKING OPTIMIZATION

In this module, you will use the NMSM Pipeline Tracking Optimization tool with your personalized skeletal geometry and neural control model to create a dynamically consistent synergy driven walking simulation that closely reproduces experimental joint motion, joint moment, ground reaction, and EMG data. **Tracking Optimization** results always provide the starting point for the NMSM Pipeline **Treatment Optimization** Process. In the subsequent module, you will perform a **Verification Optimization** (VO) starting from your TO results to confirm that your TO results are reliable, followed by a **Design Optimization** (DO) to design a synergy-based FES treatment for our stroke subject.

Similar to module 2, synergy-driven TO problems can take a long time to run, and so this project will focus more on analysis of synergy-driven TO results rather than on running multiple iterations of TO. You will only do one synergy-driven TO run, where the focus is primarily on getting the optimization to converge, not on getting an optimal result. A results directory with final results will be provided as a part of the assignment for analysis.

Synergy-driven TO problems are formulated similarly to torque-driven TO problems, but are generally much harder for the optimal control solver to find a solution. This difficulty arises because actuating a model with muscles is more complicated than discrete torque actuators on muscle joints. The additional problem complexity means that Synergy-driven TO is very sensitive to the input data it is given. If the prerequisite model personalization was not high quality, synergy-driven TO will often struggle to converge on a good solution. However, if effort is put out to get good quality and anatomically correct model personalization results, synergy-driven TO can converge quickly.

Synergy-driven TO makes use of input data from a variety of sources throughout the NMSM Pipeline. Firstly, you should always use a post-JMP OpenSim model. If your problem includes ground reactions, TO expects an osimx file containing a GCP contact surface set. Because synergy-driven TO uses muscle actuators, it requires both MTP results and NCP results. TO uses the calibrated muscle models from NCP and uses the muscle synergies calculated by NCP as an initial guess to calculate more dynamically consistent synergies. Finally, synergy-driven TO expects that your input data is formatted as prescribed by the preprocessing sub-tool. Therefore, you will be using the same preprocessing directory that you have used for the previous subprojects. The interaction between tool settings, data, and models required to perform this module is shown in the figure below:

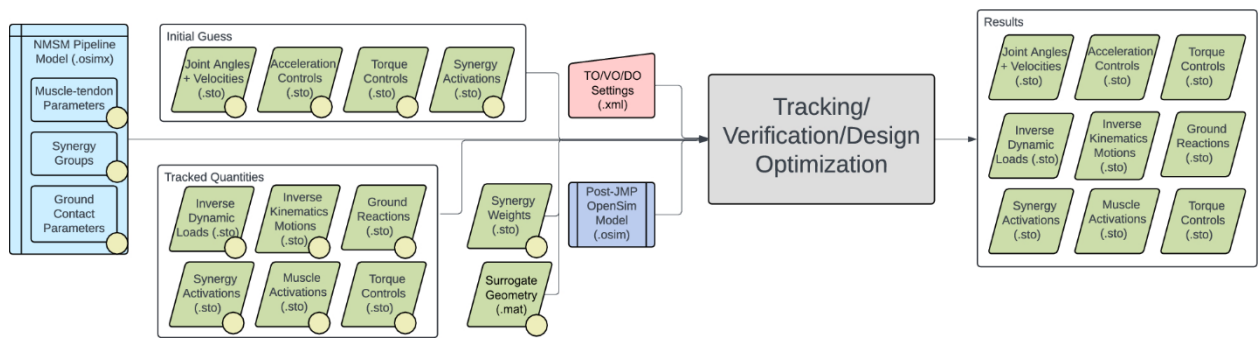

##### Module Task 1: Surrogate Muscle Geometry

An important step before all synergy-driven Tracking Optimization runs is to create a surrogate muscle geometry model. As we explored briefly in module 1, muscle geometry calculations are computationally expensive. Running the muscle analysis tool with all 86 muscles in the RCNL2025 model can take minutes for just one gait cycle. This was not problematic for MTP and NCP because the motion was not being changed by those tools, and so muscle analysis only needed to be run once. For Treatment Optimization however, the motion changes every iteration, and so muscle geometry needs to be recalculated repeatedly. This quickly becomes a problem for generating walking simulations.

The solution to this problem is to create a surrogate muscle geometry model that allows for muscle geometry calculations to be done much quicker. Rather than running a full muscle analysis calculation multiple times, we can fit a 6<sup>th</sup> order polynomial to the muscle geometry as a function of modeled joint angles. In short, we can run muscle analysis on a gait cycle and fit the muscle-tendon lengths and muscle-moment arms to the modeled joint angles. This polynomial model allows us to quickly and easily calculate muscle moment arms for a given model orientation.

The next problem that arises is selecting the proper motions to fit a surrogate model with. If we only use gait data to fit a surrogate model, then any new motions that deviate from the original motion will incur errors in muscle moment arms. On the other hand, we don't want to exhaustively sample every combination of model coordinates, because that will take far too much computation time to fit the surrogate model. The solution is to meet in the middle, and sample combinations of coordinates in the neighborhood around the original gait data. As such, the surrogate geometry uses a Latin hypercube sampling (LHS) algorithm to sample combinations of coordinates close to the original motion. LHS randomly searches combinations of coordinates using a minimal set of combinations that effectively covers the entire search space. We define the search space here to be the joint angles during gait motion $\pm 20^\circ$.

##### Step 1) Create your surrogate kinematics

1. Open the Matlab script `surrogateKinematicsScript.m`
2. Fill out the variables for `modelFileName`, and `referenceKinematicsFile`.
  a. `referenceKinematicsFile` should be your IK data in `preprocessing`.
3. Set `angularPadding` value to `20°`
4. Set `linearPadding` to `0.1m`.
5. Click Run.
6. Start OpenSim and load `UF_Subject_4_Scaled_JMP.osim`
7. Load the motion `surrogateData\IKData\gait_1.sto`
  a. Analyze what is happening in this motion. You will be asked about it at the end of the assignment.

This surrogate kinematics script samples joint angles in the area around the given gait data. Ideally, these sampled joint angles will encompass any new predicted motions and allow their moment arms to be accurate and quickly calculated.

##### Step 2) Run muscle analysis

1. Open the Analyze tool in OpenSim.
2. Set your input motion to `surrogateData\IKData\gait_1.sto`
  a. Do **not** filter this motion.

b. The motion looks very noisy but that is by design. The surrogate kinematics script adds “noise” to the base motion so that the sampling area for moment arms is larger.
3. Set your Prefix to **gait\_1**
4. Set your output directory to **surrogateData\MAData\gait\_1**
5. Add a muscle analysis set
6. Click run. This muscle analysis will take about 20 minutes.

You now have a set of muscle-tendon lengths and moment arms for every combination of joint angles produced in step 1. TO will automatically fit a polynomial to these values at the start of your run.

#### Module Task 2: Initial Tracking Optimization

This module task will focus on getting an initial guess synergy-driven Tracking Optimization. Because synergy-driven TO is such a complicated problem, one of the primary challenges in getting a good result is simply getting the optimal control problem to converge in the first place. A good method to get an initial TO to converge is to start the problem with very loose cost terms, and tight constraint terms. Once this loosely defined optimization converges, the next step is to progressively and systematically tighten cost terms and constraint terms until a satisfactory solution is obtained. This process allows us to diagnose if the constraint terms being used are compatible with each other, and systematically tightening the problem allows us to diagnose where the primary difficulties in convergence may come from.

##### Step 1) Create TO Settings File

1. Load your post-JMP model **UF\_Subject\_4\_Scaled\_JMP.osim** into the OpenSim GUI
2. Open the Tracking Optimization GUI
3. Set the *Input Osimx File* to be the Osimx file generated by your final NCP run
4. Set the *Tracked Quantities Directory* to **Preprocessing\preprocessed**
5. Set the *Initial Guess Directory* to your final NCP results directory
6. Set the *Trial Prefix* to **gait\_1**
7. Set the *Results Directory* to **TrackingOptimization\SynergyTOResultsV1**
8. Set the *Optimal Control Settings File* to **TrackingOptimization\gpopsSettings.xml**
9. Add all unlocked coordinates to States Coordinates List.
10. Go to the *RCNL Controllers* tab
11. Keep *Optimize Synergy Vectors* disabled
12. For the *Coordinates List*, select all unlocked lower limb coordinates
13. Set the *Surrogate Model Data Directory* to **TrackingOptimization\surrogateData**
14. Go to the *Cost/Constraints* tab
15. Add cost and constraint terms according to the following tables:

##### Cost Terms

| Name | Type | Components | Max Allowable Error |
| --- | --- | --- | --- |
| lower limb coordinate tracking loose | **generalized_coordinate_tracking** | **hip_flexion_r**<br>**knee_angle_r**<br>**ankle_angle_r**<br>**hip_flexion_l**<br>**knee_angle_l**<br>**ankle_angle_l** | 0.3 |
| Lower limb coordinate tracking tight | **generalized_coordinate_tracking** | **hip_adduction_r**<br>**hip_rotation_r**<br>**subtalar_angle_r**<br>**mtp_angle_r**<br>**hip_adduction_l**<br>**hip_rotation_l**<br>**subtalar_angle_l**<br>**mtp_angle_l** | 0.15 |
| Upper limb coordinate tracking loose | generalized_coordinate_tracking | lumbar_extension<br>lumbar_bending<br>lumbar_rotation<br>arm_flex_r arm_add_r<br>arm_rot_r elbow_flex_r<br>arm_flex_l arm_add_l<br>arm_rot_l elbow_flex_l | 0.15 |
| Pelvis translation tracking | generalized_coordinate_tracking | pelvis_tx pelvis_tz | 0.05 |
| Pelvis rotation tracking | generalized_coordinate_tracking | pelvis_tilt pelvis_list<br>pelvis_rotation | 0.15 |
| ID load tracking | inverse_dynamics_load_tracking | hip_flexion_r_moment<br>hip_adduction_r_moment<br>hip_rotation_r_moment<br>knee_angle_r_moment<br>ankle_angle_r_moment<br>subtalar_angle_r_moment<br>hip_flexion_l_moment<br>hip_adduction_l_moment<br>hip_rotation_l_moment<br>knee_angle_l_moment<br>ankle_angle_l_moment<br>subtalar_angle_l_moment | 25 |
| Speed tracking | generalized_speed_tracking | All unlocked coordinates | 5 |
| Vertical external force tracking | external_force_tracking | ground_force_1_vy<br>ground_force_2_vy | 100 |
| Horizontal external force tracking | external_force_tracking | ground_force_1_vx<br>ground_force_1_vz<br>ground_force_2_vx<br>ground_force_2_vz | 25 |
| External moment tracking y z | external_moment_tracking | ground_moment_1_my<br>ground_moment_1_mz<br>ground_moment_2_my<br>ground_moment_2_mz | 20 |
| External moment tracking x | external_moment_tracking | ground_moment_1_mx<br>ground_moment_2_mx | 20 |
| Muscle activation tracking | muscle_activation_tracking | *All model muscles* | 0.25 |
| Synergy activation tracking | controller_tracking | right_leg_1 right_leg_2<br>right_leg_3 right_leg_4<br>right_leg_5 left_leg_1<br>left_leg_2 left_leg_3<br>left_leg_4 left_leg_5 | 0.5 |

#### Constraint Terms

| Name | Type | Components | Max/Min Error |
| --- | --- | --- | --- |
| Kinetic consistency | kinetic_consistency | hip_flexion_r_moment<br>hip_adduction_r_moment<br>hip_rotation_r_moment<br>knee_angle_r_moment<br>ankle_angle_r_moment<br>subtalar_angle_r_moment<br>hip_flexion_l_moment<br>hip_adduction_l_moment<br>hip_rotation_l_moment<br>knee_angle_l_moment<br>ankle_angle_l_moment<br>subtalar_angle_l_moment | 0.1/-0.1 |
| Residual force reduction | root_segment_residual_load | pelvis_tx_force<br>pelvis_ty_force<br>pelvis_tz_force | 25/-25 |
| Residual moment reduction | root_segment_residual_load | pelvis_tilt_moment<br>pelvis_list_moment<br>pelvis_rotation_moment | 10/-10 |
| Rotational coordinate periodicity | generalized_coordinate_periodicity | pelvis_tilt pelvis_list<br>pelvis_rotation<br>hip_flexion_r<br>hip_adduction_r<br>hip_rotation_r<br>knee_angle_r<br>ankle_angle_r<br>subtalar_angle_r<br>mtp_angle_r<br>hip_flexion_l<br>hip_adduction_l<br>hip_rotation_l<br>knee_angle_l<br>ankle_angle_l<br>subtalar_angle_l<br>mtp_angle_l<br>lumbar_extension<br>lumbar_bending<br>lumbar_rotation<br>arm_flex_r<br>arm_add_r<br>arm_rot_r<br>elbow_flex_r<br>arm_flex_l<br>arm_add_l<br>arm_rot_l | 0.01/-0.01 |
|  |  | elbow_flex_l |  |
| Translational coordinate periodicity | generalized_coordinate_periodicity | pelvis_tx<br>pelvis_ty<br>pelvis_tz | 0.01/-0.01 |
| External force periodicity | external_force_periodicity | ground_force_2_vx<br>ground_force_2_vy<br>ground_force_2_vz<br>ground_force_1_vx<br>ground_force_1_vy<br>ground_force_1_vz | 5/-5 |
| External moment periodicity | external_moment_periodicity | ground_moment_2_mx<br>ground_moment_2_my<br>ground_moment_2_mz<br>ground_moment_1_mx<br>ground_moment_1_my<br>ground_moment_1_mz | 1/-1 |
| Rotational speed periodicity | generalized_speed_periodicity | pelvis_tilt pelvis_list<br>pelvis_rotation<br>hip_flexion_r<br>hip_adduction_r<br>hip_rotation_r<br>knee_angle_r<br>ankle_angle_r<br>subtalar_angle_r<br>mtp_angle_r<br>hip_flexion_l<br>hip_adduction_l<br>hip_rotation_l<br>knee_angle_l<br>ankle_angle_l<br>subtalar_angle_l<br>mtp_angle_l<br>lumbar_extension<br>lumbar_bending<br>lumbar_rotation<br>arm_flex_r<br>arm_add_r<br>arm_rot_r<br>elbow_flex_r<br>arm_flex_l<br>arm_add_l<br>arm_rot_l<br>elbow_flex_l | 0.1/-0.1 |
| Translational speed periodicity | generalized_speed_periodicity | pelvis_tx<br>pelvis_ty<br>pelvis_tz | 0.01/-0.01 |

16. Inside gpopsSettings.xml, change `<setup_nlp_max_iterations>` to 250
17. Inside your settings file, inside `<RCNLSynergyController>`, copy and paste:

```
<load_surrogate_model>true</load_surrogate_model>
```

Fitting the surrogate model can take a long time for our model, so this line loads a preexisting surrogate model into the TO run to save time.

18. Open the Matlab script `TrackingOptimization\runTO.m` and click Run
19. This TO run will take approximately an hour to run on the Duncan Hall Computers

##### Step 1a) Analyze Convergence

This step should be done while your TO is running.

After clicking Run on Matlab, you will see a similar output to the following appear after about 10 minutes:

| iter | objective | inf pr | inf du | lg(mu) | \|\|d\|\| | lg(rg) | alpha du | alpha pr | ls |
| --- | --- | --- | --- | --- | --- | --- | --- | --- | --- |
| 0 | 9.1398349e-02 | 2.10e+03 | 6.14e-03 | 0.0 | 0.00e+00 | - | 0.00e+00 | 0.00e+00 | 0 |
| 1 | 9.1034505e-02 | 2.07e+03 | 4.22e+00 | -2.3 | 1.35e+00 | - | 9.98e-03 | 2.11e-02f | 1 |
| 2 | 9.1321351e-02 | 1.99e+03 | 2.52e+01 | -2.3 | 1.11e+00 | - | 1.21e-02 | 3.54e-02f | 1 |
| 3 | 9.3515725e-02 | 1.82e+03 | 1.19e+02 | -2.3 | 1.94e+00 | - | 2.23e-02 | 6.91e-02h | 1 |

The main important columns in this output are `iter`, `objective`, `inf_pr`, and `inf_du`. The `objective` column is the value for the cost function, where a lower value is a better adherence to the cost function. `inf_pr` and `inf_du` are the primal and dual infeasibility, respectively, and correspond to constraint satisfaction. Specifically, `inf_pr` corresponds to how well constraints are satisfied, and `inf_du` corresponds to how well the cost function and constraint function agree with each other. If `inf_pr` is low (constraints are satisfied), but `inf_du` is high, that roughly means that the cost function could be lower than it currently is. In this case, the constraints might be satisfied, but the solution is not “optimal” because the cost function is still too high. The optimization is configured to converge when both `inf_pr` < 9.99e-5 and `inf_du` < 9.99e-1.

As mentioned earlier in the assignment, a key challenge in synergy-driven TO is simply getting the optimization to converge. The optimization you are running should take around 200 iterations to converge. During this time, if `inf_pr` > 1.00e4 or `inf_du` > 1e6 for more than 10 iterations, it is unlikely that your optimization will converge. To save time, you should stop the optimization and troubleshoot your settings file. Another sign that your optimization will likely not converge is if the letter “r” appears next to your iteration counter. This sign means that the optimizer entered “restoration” mode. Put simply, the optimizer found that the current search space was not feasible (meaning the constraints cannot be satisfied). While entering restoration mode does not always mean the optimization will not converge, it is typically good practice to cancel the optimization if it occurs because it is likely that the problem as a whole is not feasible.

Some likely problems that can cause an optimization to not converge are:

1) Data quality issues: This will likely not be a problem for this project but is important if you are doing your own research.
2) Conflicting constraint terms: If you have constraint terms with conditions that conflict with each other, the problem as a whole will be infeasible and `inf_pr` and `inf_du` will quickly increase.

3) Cost terms conflict with constraint terms: This often happens if cost terms are too tight and don't give the optimizer freedom to satisfy constraints. Alternatively, if you include inappropriate cost terms, constraints will also be difficult to match. An example of this is tracking pelvis\_ty in the cost function. This cost term will not work well with residual reduction, because tracking pelvis\_ty forces the model into the ground which can increase the vertical residual force.
4) Other infeasible constraints: Often constraints are simply not able to be satisfied. For example, in the kicking tutorial, we increase the toe marker velocity by using a constraint term to force the X velocity of the toe marker to be 14.3-14.4 m/s. If however, you include the Y and Z velocities in that constraint terms, the problem will not converge because those velocities will not reasonably reach 14.3-14.4 m/s

Check your settings file for these issues, and if you still are not sure why your optimization is not converging, email the instructor for help.

#### **Step 2) Analyze Full TO Results**

The previous section walked you through how to create an initial TO settings file. That TO should have converged, thereby demonstrating problem feasibility, but the settings still needed to be iterated on to get the best solution. Next, you can analyze a final TO run from the end of that iteration process.

In this run, cost and constraint terms were systematically tightened to yield an optimal solution. Another key change was that synergy vectors were optimized as well. We found that allowing small changes in synergy vectors is important for getting a good TO solution. To go along with changing the synergy vectors, we also track the NCP synergy vectors to make sure they don't change too much.

In the TO run you did in step 1, synergy vectors were not optimized, meaning we effectively assumed that the synergy vectors created by NCP were perfect. While the synergy vectors created by NCP should be very close to the true synergy vectors, giving TO some freedom to change them often helps get a better solution. One drawback to allowing synergy vectors to change is that it slows down the optimization significantly.

To view the settings file used to get these results, you may open **SynergyTOResultsFull\TOSettings.xml**. You can plot these TO results compared to your initial TO results by uncommenting the code on line 7 inside **runTO.m**.

#### Deliverables:

Inside one PDF file, submit all plots outputted by your TO run, and answer the following questions.

1. In 1-2 sentences, describe the goal of Tracking Optimization
2. Why is synergy-driven tracking optimization generally harder than torque-driven tracking optimization? Give 2 reasons.
3. What is the main reason we want to use a surrogate muscle geometry model?
4. Briefly describe the process for creating a surrogate muscle geometry model.
5. Describe the motion that is outputted from `surrogateKinematicsScript.m`

Regarding the TO run you did on your own (step 1):

1. Why do you think some coordinate tracking terms have different max allowable errors than others?
2. **Briefly** describe the tracking quality for the following quantities. Mention the overall tracking quality, which components had better or worse tracking than others, and if you think the max allowable errors should be increased for the relevant cost term:
  a. Joint Angles
  b. Joint Velocities
  c. Joint Loads
  d. External Loads
  e. Muscle Activations
  f. Synergy Activations
3. If you were to iterate on this TO solution, what cost/constraint terms would you prioritize changing.
  a. **Exploration opportunities:** Feel free to change the max allowable errors and re-run TO. How did the results change? Were there any unexpected changes?

Regarding the full TO results given with the project (step 2):

1. Which constraint terms were changed between your initial TO run and the final TO run? Why were the other constraint terms kept the same?
2. Why do you think it is important to change synergy vectors in TO?
3. Did the synergies change very much during the TO run? Do the trends you observed in the synergies from module 2 still hold true?
4. It is important for TO results to visualize the TO results motion compared to the experimental motion. Follow the following steps to visualize your motion in OpenSim:
  a. Open up 2 instances of `UF_Subject_4_Scaled_JMP.osim` and make them different colors
    i. Right click on the *Bodies* tab under the model, and click Display>Colors
  b. Ensure both models have a zero model offset
    i. Right click on the model, and click Display>Model Offset
  c. Right click on each model, and click *Load Motion*
  d. Load `Preprocessing\preprocessed\IKData\IKDataFixedTime.sto` to one model
    i. `IKDataFixedTime.sto` is the same data as `gait_1.sto`, but the time column is offset to start at $t=0$.

e. Load **TrackingOptimization\SynergyTOResultsFull\IKData\gait\_1.sto** to the other model.
f. Sync both IK motions.
  i. Under the *Motions* tab for each model, highlight the motions (called “DataType=double”), right click them, and select *Sync Motions*
  ii. To select multiple motions at once on Windows, hold down ctrl and click on each motion.
g. Play the motions at 0.1x speed.
5. Qualitatively describe how well the TO simulated tracks the experimental motion. Are there any significant differences between the motions? Why do you think that might be?

#### MODULE 4: SYNERGY-DRIVEN VERIFICATION AND DESIGN OPTIMIZATION

In this module, you will use the Design Optimization (DO) tool with your previously generated Tracking Optimization (TO) walking simulation results to design a synergy-based FES treatment protocol for a stroke subject. You will first use Verification Optimization (VO) to sanity check your TO solution, and then you will implement cost terms in DO to modify the subject's gait in the desired way.

Because strokes involve neural impairments, your modeled treatments will focus on modifying the synergy controls produced by TO. This contrasts with the previous project with torque-driven Treatment Optimization, where the changes in DO were primarily in the subject's kinematics. This different treatment methodology means that the results might be a little bit less intuitive and take more effort to analyze.

##### Module Task 1) Verification Optimization

For this module task, you will use the NMSM Pipeline **Verification Optimization (VO)** tool to verify that the muscle synergy controls found by your Tracking Optimization (TO) produce the same walking motion and ground reactions as the TO did, but without tracking those quantities. VO will seek to minimize changes in your TO muscle synergy controls so that a dynamically consistent, periodic walking motion is produced. Results generated by your TO will provide the starting point for your VO.

VO serves as a “sanity check” for your TO results before moving further into the Treatment Optimization Process. It is possible that your TO run could have converged to a good-looking solution but had problems that will make a Design Optimization (DO) difficult. VO allows you to test that the input data that you give to DO is consistent with itself. Additionally, VO may serve as a “dry run” for your DO run. In other words, your VO run will be identical to your DO run, but without any of the design elements of the DO run. For example, in this project, we will design a new walking motion that changes synergy activations and ground reactions. As such, the VO run will look identical to the DO run, with all relevant constraints that will be used in DO, but without the cost terms that change the synergy activations and ground reactions.

A synergy-driven VO should converge in around 50 iterations, and the VO solution should look close to identical to the TO solution. In the case that your VO does not converge quickly, it is likely caused by one or more of the following:

1. You added constraint terms to your VO that are inconsistent with the solution. This often leads to the optimizer going into restoration mode.
2. The cost terms used in your VO somehow conflict with each other or other constraint terms
3. Your TO solution itself has a problem that needs to be fixed.

#### Step 1) Create your VO settings file

18. Create a copy of **TrackingOptimization\TOSettings.xml** into **VerificationOptimization**.
19. Rename the copied file **VOSettingsTemplate.xml**
20. Open **VOSettings.xml** in a text editor and change the XML headings at the top and bottom of the file from **<TrackingOptimizationTool>** to **<VerificationOptimizationTool>**
21. Disable **<optimize\_synergy\_vectors>**
22. Load your post-JMP model **UF\_Subject\_4\_Scaled\_JMP.osim** into the OpenSim GUI
23. Open the Verification Optimization GUI
24. Load your settings file **VOSettings.xml** into the Verification Optimization GUI.
25. Set the *Input Osimx File* to **NeuralControlPersonalization\NCPResultsBilateral\UF\_Subject\_4\_Scaled\_JMP\_gcp\_mtp\_mtp\_ncp.osimx**
  a. If the text box is red when you load the GUI in, you must re-select this file or the GUI will not be able to find it. The GUI does some light parsing through this file to locate ground reaction force and moment names.
26. Set the *Tracked Quantities and Initial Guess Directories* to **TrackingOptimization\TOResults**.
27. Set the *Results Directory* to **VerificationOptimization\VOResults**
28. Set the *Surrogate Model Data Directory* to **TrackingOptimization\surrogateData**
29. Click to **Cost/Constraints**
30. Delete all cost terms except for:
  a. Upper Limb Coordinate Tracking
  b. Pelvis Translation Tracking
  c. Pelvis Rotation Tracking
31. Add the following cost terms:

| Name | Type | Components | Max Allowable Error |
| --- | --- | --- | --- |
| Synergy Controller Tracking | **controller_tracking** | **right_leg_1**<br>**right_leg_2**<br>**right_leg_3**<br>**right_leg_4**<br>**right_leg_5**<br>**left_leg_1**<br>**left_leg_2**<br>**left_leg_3**<br>**left_leg_4**<br>**left_leg_5** | 1 |
| Toes Coordinate Tracking | **generalized_coordinate_tracking** | **mtp_angle_r**<br>**mtp_angle_l** | 0.025 |

32. Add the following constraint terms:

| Name | Type | Components | Max/Min Error |
| --- | --- | --- | --- |
| Initial Coordinate Deviation | **initial_generalized_coordinate_deviation** | **All coordinates in the States Coordinate List** | 0.01/-0.01 |
| Initial Speed Deviation | `initial_generalized_speed_deviation` | All coordinates in the States Coordinate List | 0.1/-0.1 |
| Ground Reaction Forces 1 Swing Phase | `external_force_value` | `ground_force_1_vx`<br>`ground_force_1_vy`<br>`ground_force_1_vz` | 1/-1 |
| Ground Reaction Forces 2 Swing Phase | `external_force_value` | `ground_force_2_vx`<br>`ground_force_2_vy`<br>`ground_force_2_vz` | 1/-1 |
| Ground Reaction Moments 1 Swing Phase | `external_moment_value` | `ground_moment_1_mx`<br>`ground_moment_1_my`<br>`ground_moment_1_mz` | 1/-1 |
| Ground Reaction Moments 2 Swing Phase | `external_moment_value` | `ground_moment_2_mx`<br>`ground_moment_2_my`<br>`ground_moment_2_mz` | 1/-1 |

33. Save this settings file as `VerificationOptimization\VOSettings.xml`
34. Open `VOSettings.xml` in a text editor.
35. Inside your Ground Reaction Forces 1 Swing Phase and Ground Reaction Moments 1 Swing Phase constraint terms, copy and paste the following:
  `<time_ranges>0.17 0.48</time_ranges>`
36. Inside your Ground Reaction Forces 2 Swing Phase and Ground Reaction Moments 2 Swing Phase constraint terms, copy and paste the following:
  `<time_ranges>0.7 1</time_ranges>`
37. These constraint terms enforce that the ground reactions must be zero during swing phase. The time ranges are in units of normalized time (0 to 1), and make the constraint only active for a specified time window.
  a. We generally need these added constraints in DO but not in TO. This is because the ground reactions and joint positions/velocities are actively tracked in TO. They are not, however, tracked in DO, and so the DO solution can “cheat” by changing initial conditions or allowing ground reactions during swing phase.
38. Run the Matlab script `VerificationOptimization\runVO.m`
39. This VO run will take approximately 50 iterations to converge
40. If the optimization enters restoration mode as explained in the previous subproject, cancel the optimization and email the instructor.

As explained earlier, the purpose of VO is to be sanity check for your TO results and DO problem formulation. Your VO results should be closely identical to your TO results. If you get VO results that are different from your TO results, you have a problem. An example of a problematic VO run is inside `VOResultsFailed`. You can run the code on line 9 to explore the results. You will be asked about them in the deliverables section.

#### Module Task 2) Plan the Treatment

As we analyzed during NCP in the second module, the synergies in this run all have distinct functions throughout the gait cycle. These functions and the muscles in each synergy are summarized in the table below:

| Synergy Number | Synergy Function | Primary Muscle Groups in Synergy Vector |
| --- | --- | --- |
| 1 | Weight Acceptance | Hip Flexors, Hip Extensors |
| 2 | Propulsion | Ankle Plantarflexors |
| 3 | Propulsion/Weight Acceptance | Ankle Plantarflexors, Hip Abductors |
| 4 | Leg Swing | Hip Extensors, Hip Flexors |
| 5 | Flexion | Hip Flexors |

We explain this subject's impairment by characterizing synergies as healthy, impaired, or compensatory based on their relative magnitudes between legs. The magnitudes of the synergies are listed in the table below:

| Synergy # | Right Leg Activation Magnitude | Left Leg Activation Magnitude |
| --- | --- | --- |
| 1 | 0.30 | 0.21 |
| 2 | 0.18 | 0.22 |
| 3 | 0.18 | 0.3 |
| 4 | 0.12 | 0.17 |
| 5 | 0.22 | 0.22 |

The primary impairment for this subject occurs in right leg synergy 3, where the magnitude of synergy activation 3 is much higher on the non-paretic (left) side than on the paretic (right) side. In addition to the primary impairment, there are several compensatory synergies across both legs, where a synergy activation is marginally larger on one leg compared to the other. Synergy 1 is compensatory on the right leg. Synergies 2 and 4 are compensatory on the right leg. Synergy 5 is healthy.

These synergy asymmetries manifest themselves in the ground reaction forces. Specifically, the braking and propulsive impulses are highly asymmetric between the legs. The paretic side (right) has a lower propulsive and higher braking impulse compared to the non-paretic side. The specific numbers are shown in the table below.

|  | Propulsive Impulse | Braking Impulse |
| --- | --- | --- |
| Right Leg | 5.36 | 14.87 |
| Left Leg | 18.25 | 5.26 |

Your goal for this section is to plan a treatment to simulate for this subject. Your goal is to design a treatment that eliminates the asymmetries in synergy activations and analyze how the propulsive and braking impulses respond to that treatment. For simplicity, this treatment will only involve scaling the synergy activation magnitudes, and as such, the shapes will not change. You may equalize the synergy activations using one of the following methods:

1. **Meet in the middle** – Calculate the average magnitude of both synergy activations, and scale the larger synergy activation down to the average, and the larger synergy activation up to the average.
2. **Increase the smaller synergy** – Scale the smaller synergy activation up to the magnitude of the larger synergy activation
3. **Decrease the larger synergy** – Scale the larger synergy activation down to the magnitude of the smaller synergy activation
4. **Any combination of the above.**

Your job for this task is to choose a method of equalizing synergy activations between the legs and calculate the scale factors required for your method. Fill out the following table with the scale factors you are applying to each synergy activation. The scale factors should be calculated with simple multiplications on the max magnitude of the synergy activations. If one synergy has a magnitude of 0.5 and you want it to increase to 0.75, then the scale factor is $0.75/0.5=1.5$. <u>You will not be graded on whether you chose the “correct” method of equalizing the synergy activations. You will however be expected to justify *why* you chose your method of scaling synergy activations.</u>

The goal of our synergy FES protocol is to make the subject more symmetric by increasing or decreasing the magnitude of synergy activations. With FES treatments, it is important to consider the subject’s limitations, and how FES can be used to change their synergies. Importantly, FES can only ever add to muscle activations. FES cannot reduce muscle activations. However, when equalizing synergy activations, we might want to reduce the magnitude of certain compensatory synergies. When planning a treatment of this type, we are making an implicit assumption that we are using FES to increase impaired synergies, and the subject will respond on their own by equalizing compensatory synergies without needing to directly stimulate those synergies. By extension, we are also assuming that the subject will be able to change their other synergies that are not being directly stimulated, which might not be true. This is a possible limitation that can only be truly studied by implementing our desired treatment on the original subject.

| Synergy | Right Leg<br>Scale Factor | Left Leg<br>Scale Factor |
| --- | --- | --- |
| 1 |  |  |
| 2 |  |  |
| 3 |  |  |
| 4 |  |  |
| 5 |  |  |

##### Module Task 3) Design Optimization

Now that you have a planned treatment design that equalizes the synergy activations, you can create a DO settings file that implements your treatment design. This DO will have two components – First we will scale the synergy activations as you designed in task 2, and then we will add cost terms that equalize the propulsive and braking impulses. will be done in two stages – First, we will just scale the synergy activations as prescribed in task 2. Next, we will add cost terms for the propulsive and braking impulses that track the scaled synergy activations.

1. Create a copy of **VerificationOptimization\VOSettings.xml** into **DesignOptimization**.
2. Rename this copied file to **DOSettingsTemplate.xml**.
3. Open **DOSettingsTemplate.xml** in a text editor and change the XML headings at the top and bottom of the file from **<VerificationOptimizationTool>** to **<DesignOptimizationTool>**
4. Load your post-JMP model **UF\_Subject\_4\_Scaled\_JMP.osim** into the OpenSim GUI
5. Open the Design Optimization GUI
6. Load your settings file **DOSettingsTemplate.xml** into the Design Optimization GUI.
7. Set the *Input Osimx File* to **NeuralControlPersonalization\NCPResultsBilateral\UF\_Subject\_4\_Scaled\_JMP\_gcp\_mtp\_mtp\_ncp.osimx**
  a. If the text box is red when you load the GUI in, you must re-select this file or the GUI will not be able to find it. The GUI does some light parsing through this file to locate ground reaction force and moment names.
8. Set the *Tracked Quantities and Initial Guess Directories* to **VerificationOptimization\VOResults**.
9. Set the *Results Directory* to **DesignOptimization\DOResults**
10. Set the *Surrogate Model Data Directory* to **TrackingOptimization\surrogateData**
11. Save this settings file as **DOSettings.xml**
12. Open **DOSettings.xml** in a text editor.
13. Inside **<RCNLCostTermSet>**, set **Synergy Controller Tracking** to **false**
14. Copy and paste the following cost terms into your **<RCNLCostTermSet>**

```
<RCNLCostTerm name="Synergy activation tracking right 1">
  <is_enabled>true</is_enabled>
  <type>controller_tracking</type>
  <controller_list>right_leg_1</controller_list>
  <scale_factor></scale_factor>
  <max_allowable_error>1</max_allowable_error>
</RCNLCostTerm>
<RCNLCostTerm name="Synergy activation tracking left 1">
  <is_enabled>true</is_enabled>
  <type>controller_tracking</type>
  <controller_list>left_leg_1</controller_list>
  <scale_factor></scale_factor>
  <max_allowable_error>1</max_allowable_error>
</RCNLCostTerm>
<RCNLCostTerm name="Synergy activation tracking right 2">
  <is_enabled>true</is_enabled>
  <type>controller_tracking</type>
  <controller_list>right_leg_2</controller_list>
  <scale_factor></scale_factor>
  <max_allowable_error>1</max_allowable_error>
</RCNLCostTerm>
<RCNLCostTerm name="Synergy activation tracking left 2">
  <is_enabled>true</is_enabled>
  <type>controller_tracking</type>
```

```
    <controller_list>left_leg_2</controller_list>
    <scale_factor></scale_factor>
    <max_allowable_error>1</max_allowable_error>
</RCNLCostTerm>
<RCNLCostTerm name="Synergy activation tracking right 3">
    <is_enabled>true</is_enabled>
    <type>controller_tracking</type>
    <controller_list>right_leg_3</controller_list>
    <scale_factor></scale_factor>
    <max_allowable_error>1</max_allowable_error>
</RCNLCostTerm>
<RCNLCostTerm name="Synergy activation tracking left 3">
    <is_enabled>true</is_enabled>
    <type>controller_tracking</type>
    <controller_list>left_leg_3</controller_list>
    <scale_factor></scale_factor>
    <max_allowable_error>1</max_allowable_error>
</RCNLCostTerm>
<RCNLCostTerm name="Synergy activation tracking right 4">
    <is_enabled>true</is_enabled>
    <type>controller_tracking</type>
    <controller_list>right_leg_4</controller_list>
    <scale_factor></scale_factor>
    <max_allowable_error>1</max_allowable_error>
</RCNLCostTerm>
<RCNLCostTerm name="Synergy activation tracking left 4">
    <is_enabled>true</is_enabled>
    <type>controller_tracking</type>
    <controller_list>left_leg_4</controller_list>
    <scale_factor></scale_factor>
    <max_allowable_error>1</max_allowable_error>
</RCNLCostTerm>
<RCNLCostTerm name="Synergy activation tracking right 5">
    <is_enabled>true</is_enabled>
    <type>controller_tracking</type>
    <controller_list>right_leg_5</controller_list>
    <scale_factor></scale_factor>
    <max_allowable_error>1</max_allowable_error>
</RCNLCostTerm>
<RCNLCostTerm name="Synergy activation tracking left 5">
    <is_enabled>true</is_enabled>
    <type>controller_tracking</type>
    <controller_list>left_leg_5</controller_list>
    <scale_factor></scale_factor>
    <max_allowable_error>1</max_allowable_error>
```

```
</RCNLCostTerm>
```

15. Inside each cost term, fill out `<scale_factor>` with the scale factors you calculated in task 2.

16. Inside of your `<RCNLCostTermSet>`, copy and paste the following cost terms.

```
<RCNLCostTerm name="Propulsive impulse">
  <is_enabled>true</is_enabled>
  <type>propulsive_impulse_goal</type>
  <hindfoot_body_list>calcn_r calcn_l</hindfoot_body_list>
  <error_center>10.7</error_center>
  <max_allowable_error>4.3</max_allowable_error>
</RCNLCostTerm>
<RCNLCostTerm name="Braking impulse">
  <is_enabled>true</is_enabled>
  <type>braking_impulse_goal</type>
  <hindfoot_body_list>calcn_r calcn_l</hindfoot_body_list>
  <error_center>10.7</error_center>
  <max_allowable_error>4.3</max_allowable_error>
</RCNLCostTerm>
```

a. This cost term sets a design goal for the optimization to equalize the propulsive and braking impulses between the feet. The cost term imposes that the braking and propulsive impulses for both the left and the right feet must be 10.1 Ns. The value of 10.1 was obtained by averaging the propulsive impulses between the legs before the treatment, and then assuming the braking impulses should also be equal to that number.
17. Disable all controller tracking cost terms added in step 14.
18. Re-enable your original controller tracking cost term that tracks all synergy activations with no scale factors. This should be the same controller tracking term that was used in VO.
19. Run `DOSettings.xml`
20. After running DO, open the file `calcPropAndBrakeImpulses.xml` and change the initial guess directory to be the results directory from your stage 1 DO run. Run line 11 in `RunDO.m` and record the calculated propulsive and braking impulses.
21. Edit `calcPropAndBrakeImpulses.xml` and change the initial guess directory to be the results directory from your stage 2 DO run. Run line 11 in `RunDO.m` and record the calculated propulsive and braking impulses.

#### Deliverables:

Inside one PDF file, submit all plots outputted by your VO and DO runs, and answer the following questions.

1. One purpose of VO is to diagnose potential problems with a TO solution before getting caught up in a DO run. Diagnosing problems with a TO solution is rarely straightforward, however. Inside the VO results directory named **VOResultsFailed**, you can see VO results that tracked the synergy activations nearly perfectly, but did not reproduce the same joint angles, ID loads, and ground reactions. The diagnosis for this problem was that the kinetic consistency term in TO was too loose and needed to be tightened. Changing kinetic consistency in TO from 0.1 to 0.01 fixed the problem. **Explain why too loose of a kinetic consistency in TO might lead to the result seen in VOResultsFailed.** Remember that a kinetic consistency constraint enforces that the joint moments produced by muscles must be equal to the ID loads to within a given tolerance.
2. Fill out the table with your synergy activation scale factors:

| Synergy | Right Leg<br>Scale Factor | Left Leg<br>Scale Factor |
| --- | --- | --- |
| 1 |  |  |
| 2 |  |  |
| 3 |  |  |
| 4 |  |  |
| 5 |  |  |

3. Explain and justify the method you used to decide on synergy activation scale factors.
4. Fill out the table with the propulsive and braking impulses from your TO and DO runs.

|  |  | Propulsive<br>Impulse | Braking<br>Impulse |
| --- | --- | --- | --- |
| TO Results | Right Leg |  |  |
|  | Left Leg |  |  |
| DO Results | Right Leg |  |  |
|  | Left Leg |  |  |

5. Do you think your treatment design was a success? Why or why not? If not, what do you think you could do to make it better?
6. If you were to do another DO iteration to get a better treatment design, what would you change in your problem formulation?
7. Given your treatment design, which synergies would you stimulate with FES, and which synergies would you assume that the subject can change on their own in response to the FES treatment?
8. Do you think that your treatment design can be readily translated into a clinical synergy FES stimulation pattern? Why or why not? If not, what could you change to more accurately simulate FES for this subject?
9. What are the primary limitations of this computational treatment design?

### Supplementary Material C: Scaled Generic Model Comparison

#### Introduction

Another important issue is whether the various NMSM Pipeline Model Personalization tools substantially improve how well a musculoskeletal model can reproduce experimental data collected from a specific subject. These data include marker motion, joint motion, ground reaction, and joint moment data. Since the current standard in the field is a scaled generic musculoskeletal model, this document quantifies how much better musculoskeletal models personalized with the NMSM Pipeline can reproduce experimental measurements compared to standard scaled generic models.

#### Comparisons for Tutorial 1

For the Joint Model Personalization (JMP) module, our model scaling process involves having users perform unique steps beyond standard model scaling practices. First, users use a unique method of scaling the feet that guarantees they are flat on the ground when markers are attached to them. This step is critical for Ground Contact Model Personalization (GCP) to work well. Users also personalize the knee adduction angle during model scaling so that shank markers are attached more accurately. These unique model scaling steps resulted in a better “pre-JMP” model than a standard scaled generic model. For comparison to Figure 3 in the main text, the Figure C1 below shows marker errors for a standard scaled generic model without our extra model scaling steps. These marker tracking errors are substantially larger than those produced by our post-JMP model.

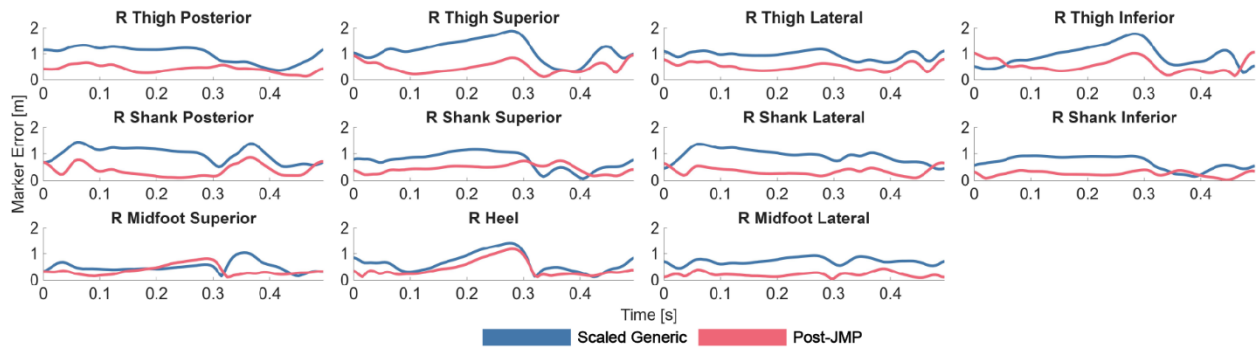

**Figure C1:** Representative marker tracking errors between a scaled generic model and our post-JMP model. The maximum and average marker errors for the scaled generic model during gait are 4.2 cm and 1.4 cm, respectively. Both of these errors are larger than the value of 2.1 cm and 0.92 cm produced by the pre-JMP model used in the tutorial. The corresponding maximum and average marker errors for the post-JMP model are 1.6 cm and 0.57 cm, respectively.

The goal of JMP is to yield more accurate joint angles from inverse kinematics and joint moments from inverse dynamics. Below is a comparison of lower body joint angles (Figure C2) and joint moments (Figure C3) produced by the scaled generic model above, the pre-JMP model presented in the main text, and the post-JMP model presented in the main text. As shown in Figure C2, root-mean-square errors (RMSEs) in joint angles between the post-JMP model and the scaled generic model are as large as $11.8^\circ$ for hip rotation and between $4.0$ and $5.9^\circ$ for ankle, subtalar, and toes angles. Errors of these magnitudes will adversely affect the ability of Muscle-tendon Model Personalization and Ground Contact Model Personalization to achieve good calibration results.

This magnitude of errors for ankle, subtalar, and toes joints will significantly hinder the ability of Ground Contact Model Personalization to achieve good calibration results. As shown in Figure C3, RMSEs in joint moments between the same two models are as large as $8.9$ Nm for knee angle moment and $5.9$ Nm for hip flexion moment. Errors of these magnitudes will adversely affect the ability of Muscle-tendon Model Personalization to achieve good calibration results.

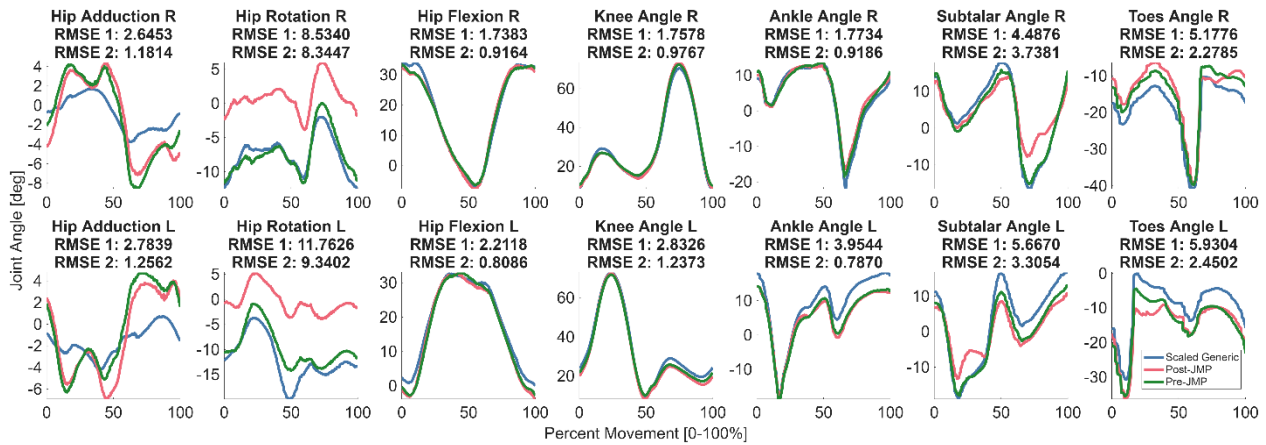

**Figure C2:** Inverse kinematics joint angles for the standard scaled generic model (blue), the pre-JMP model (green), and the post-JMP model (red). RMSE 1 indicates the difference between the post-JMP model and the scaled generic model, while RMSE 2 indicates the difference between the post-JMP model and the pre-JMP model. The RMSE values reported are with respect to the post-JMP solution, therefore a larger error indicates that Joint Model Personalization had a bigger effect on that joint.

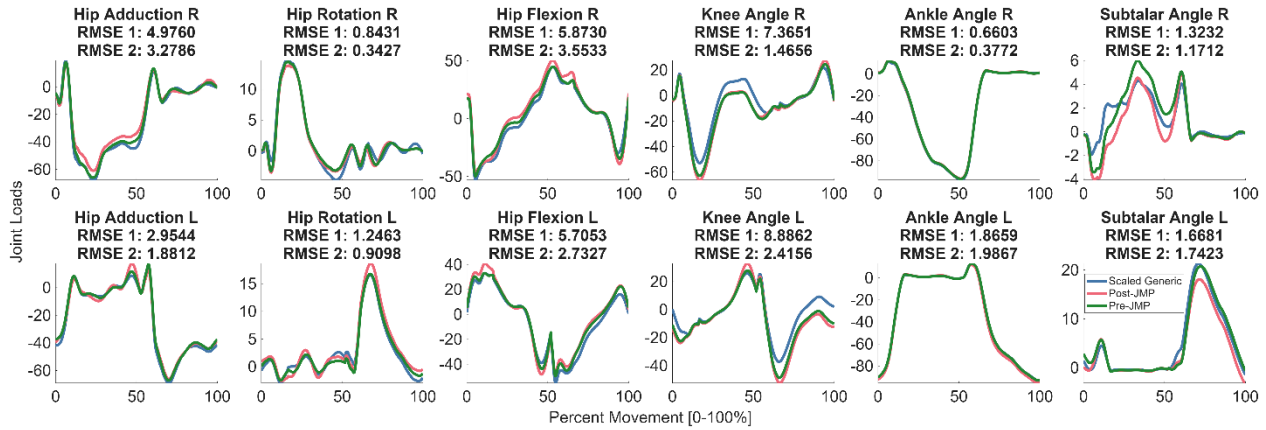

**Figure C3:** Inverse dynamics joint moments for the standard scaled generic model (blue), the pre-JMP model (green), and the post-JMP model (red). RMSE 1 indicates the difference between the post-JMP model and the scaled generic model, while RMSE 2 indicates the difference between the post-JMP model and the pre-JMP model.

For the GCP module, calibration is important to accurately match all 6 components of the ground reaction data (note that how well a foot-ground contact model reproduces experimental ground reaction moment data is almost never reported). Figure C4 shows results for a generic GCP model with only the resting spring length calibrated. Tracking quality for all ground reactions is significantly worse than that of a calibrated model. Because these results are so low quality, it is unreasonable to use them in a Tracking Optimization.

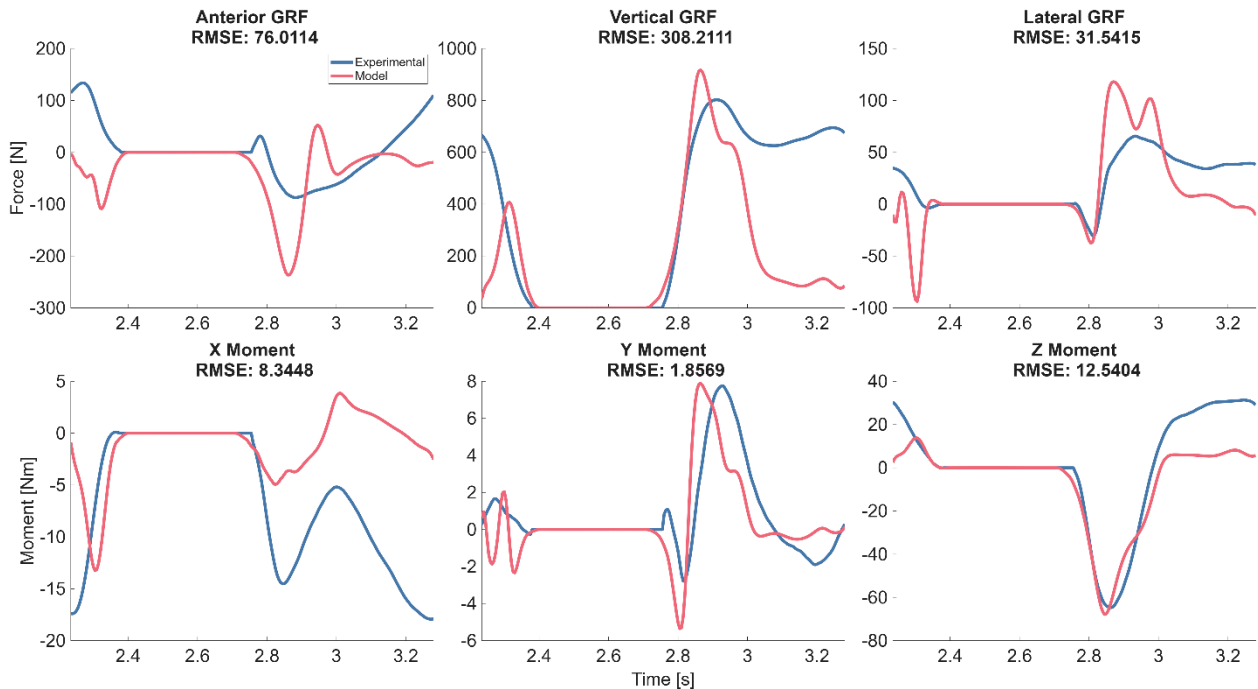

**Figure C4:** Uncalibrated GCP results. The blue curves represent experimental ground reactions, and the red curves represent ground reactions produced by an uncalibrated model.

#### Comparisons for Tutorial 2

The second tutorial uses a post-JMP model as the starting point of the model personalization process. It was shown above that a post-JMP model produces different inverse dynamics joint moments than those produced by a scaled generic model. These results are consistent with those reported by Reinbolt *et al.* (2007).

The primary personalization step in tutorial 2 is Muscle-tendon Personalization (MTP). Figure C5 shows uncalibrated MTP results with a scaled generic model. All model parameters use their initial guesses taken from the scaled generic model. This result is similarly low quality as the GCP result because MTP calibrates several important parameters that are not often calibrated. Importantly, MTP calibrates EMG scale factors to give an estimate of percent of total activation without using a maximum voluntary contraction experiment.

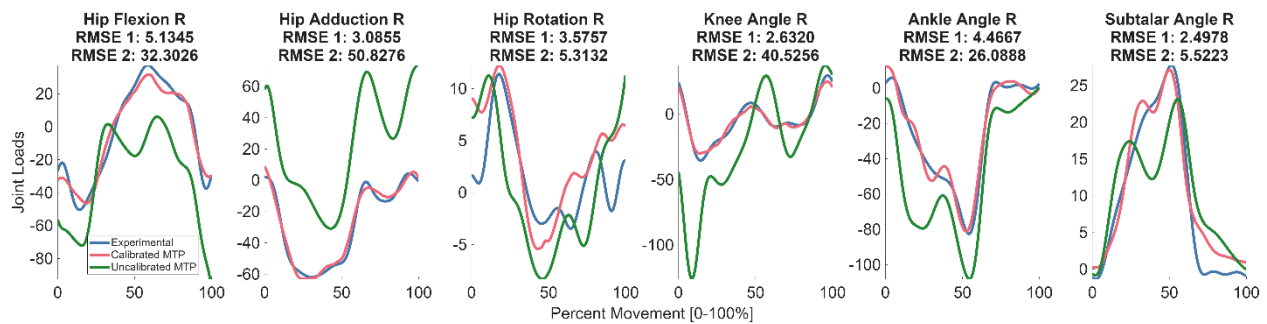

**Figure C5:** Joint moment matching achieved by a scaled generic model when used in the MTP module without any calibration of model parameter values. The blue curves represent experimental joint moments, the red curves represent a calibrated MTP model, and the green curves represent the uncalibrated MTP model. The RMSE 1 values measure the error between the experimental moments and the calibrated MTP moments. RMSE 2 values measure the error between the experimental moments and the uncalibrated MTP moments. While the MTP moments somewhat resemble the shape of the experimental joint moments, the magnitudes of the MTP joint moments are very inaccurate.

Because these MTP results are so inaccurate, it does not make sense to run Neural Control Model Personalization or any part of the Treatment Optimization process using them.

### Supplementary Material D: Reproducibility Evaluation

#### Introduction

An important issue for this article is the extent to which the presented research is truly reproducible. This question can be answered from two perspectives. The first perspective is that of *mechanistic reproduction*. For this perspective, a user starts with a scaled generic OpenSim model and all necessary experimental data, reads the tutorial instructions, and use the provided XML settings files created by the authors to run each NMSM Pipeline tool. If that approach is followed, the user will be able to recreate all steps performed by the authors to generate the same final results produced by the authors. Thus, from this perspective, the presented research is 100% reproducible. To our knowledge, no previously published neuromusculoskeletal modeling study has demonstrated 100% mechanistic reproduction, providing readers with everything needed to go from initial model and data to final model and results.

The second perspective is that of *organic reproduction*. For this perspective, a user starts with a scaled generic OpenSim model and all necessary experimental data, reads the tutorial instructions, and creates their own XML settings files to run each NMSM Pipeline tool. If that approach is followed, the user may make some different decisions along the way, or may be unable to complete some sections of a tutorial. The main distinction between mechanistic and organic reproducibility is human error. Users can arrive at different results because of reasons such as our instructions not being clear enough, or the user clicking on the wrong option. Thus, from this perspective, the presented research will be less than 100% reproducible. How much less provides a helpful indicator of how well the tutorials are written, how much flexibility is provided in some of the tutorial decisions, and how difficult it is for a user to learn and understand all tutorial steps.

Given that the presented research is 100% reproducible from the standpoint of *mechanistic reproduction*, the remainder of this document explores the extent to which the presented research is reproducible from the standpoint of *organic reproduction*.

#### Methods

The organic reproducibility of the research presented in this article was tested by implementing the tutorials as course projects in a combined undergraduate/graduate mechanical engineering course at Rice University. The course had no required prerequisite courses in programming or biomechanics, and few students had prior experience using OpenSim or the NMSM Pipeline. When developing the tutorials, the authors assumed that students had no background in coding, numerical methods, or biomechanics. Students were allowed to work in self-selected groups of two if desired.

Each tutorial was implemented as a single course project, with each module taking a single week for students to complete with two exceptions: the Joint Model Personalization module took two weeks because students had to use OpenSim to scale the model before using Joint Model Personalization, and the Torque-driven Tracking Optimization module took two weeks due to the higher computational runtime required by this module. Students were instructed to submit specific deliverables for each module, including plots showing their outputs and numerical values for errors when applicable.

The in-class lectures were structured around basic musculoskeletal modeling concepts and these tutorials, with each module having two corresponding lectures: a theoretical lecture covering how the relevant tools and modeling techniques work, and a hands-on lecture that worked through simplified tutorials with students to demonstrate how to use the relevant OpenSim and NMSM Pipeline tools. The theoretical lectures were designed to teach the “what” and the “why” of neuromusculoskeletal modeling. These lectures focused on the history of modeling techniques, how the current tools improve on older tools, and why the modeling techniques being discussed are important to the field of neuromusculoskeletal modeling. The hands-on lectures were designed to teach the “how” of neuromusculoskeletal modeling and interactively worked through basic tutorials for OpenSim and NMSM Pipeline tools. Students were encouraged to work interactively through the tutorials during these lectures. All lectures were recorded and uploaded to the course website for students to review later. All modules were assigned to students on the day of the hands-on lecture. To ensure that students struggling with previous modules could complete future modules, premade results from previous modules were released to students each time a new module was assigned.

Several resources were given outside of class to support students working through the tutorials on their own time. First, students were given 24/7 access to an on-campus computer lab with all necessary software pre-installed on all computers. The Windows computers were well equipped with 20 core CPUs and 32 GB of RAM and so could run all NMSM Pipeline tools reasonably fast. Second, students had access to weekly office hours where the course instructor and teaching assistant were available to answer questions. Third, students were provided with a frequently asked questions (FAQ) page on the course website. Each time a student emailed the instructor or teaching assistant with a question, the question and answer were anonymized and displayed on the FAQ page. Students were encouraged to check this page whenever they had a problem.

We evaluated module reproducibility by reviewing student submissions and placing them into one of three categories: completely reproduced, closely reproduced, or not reproduced. The reproducibility category into which a submission was placed was determined qualitatively by assessing key features of reported error metrics or output plots. The key features that were assessed were chosen based on their high sensitivity to the rest of the solution. For Joint Model Personalization, the key features assessed were maximum marker tracking error and marker error plots from the simultaneous run. For Ground Contact Model Personalization, the key feature assessed was the ground reaction x moment, which is exceptionally sensitive to both ground contact model parameters and foot kinematics. For torque-driven Tracking Optimization, the key feature assessed was also the ground reaction x moment. For the torque-driven Design Optimization, the key feature assessed was the peak adduction moment in each knee since that quantity defined the desired outcome for the first project. For Muscle-tendon Model Personalization, the key features assessed were the shapes and RMSE values of modeled joint moments and muscle activations. For Neural Control Model Personalization, the key features assessed were the shapes and RMSE values of modeled joint moments and muscle synergy activations. For synergy-driven Tracking Optimization, the key feature assessed was the ground reaction x moment. For synergy-driven Design Optimization, the key features assessed were the braking and propulsive impulses since those quantities defined desired outcomes of the second project. Descriptions for how the key features were used to place a submission into its reproducibility category are provided in Table D1.

**Table D1:**

| **NMSM Pipeline Tool** | **Completely Reproduced** | **Closely Reproduced** | **Not Reproduced** |
| --- | --- | --- | --- |
| Joint Model Personalization | Used the correct markers for the simultaneous run and had the correct max marker error. | Used the correct markers for the simultaneous run but had max marker error off by $\pm 0.5\text{cm}$, but less than $\pm 1.5\text{cm}$. | Did not use the correct markers, marker file, or got very little reduction in marker error. |
| Ground Contact Model Personalization | Solution converged to a set of ground contact parameters that produced a ground reaction X moment identical to that presented. | Solutions were close but featured a noticeable different shape of the ground reaction X moment, but all other ground reactions were nearly identical. | The optimization entirely failed or converged to a solution with significantly different ground reactions for all 6 DOF. |
| Torque-driven Tracking Optimization | Converged solution produced a ground reaction X moment | Solutions were close but featured a noticeably different shape of the | Tracking Optimization did not converge within the specified number of |
|  | nearly identical to that presented. | ground reaction X moment. | iterations or converged to a significantly different solution. |
| Torque-driven Design Optimization | Solution converged to a motion that achieved the desired reduction in peak knee adduction moment and produced the same knee adduction moment as those presented. | Solution converged to a motion that achieved the desired reduction in peak knee adduction moment but produced a slightly different knee adduction moment shape. | Design Optimization did not converge within the specified number of iterations or converged to a significantly different solution. |
| Muscle-tendon Model Personalization | Student successfully used Synergy Extrapolation and Muscle-tendon length Initialization and achieved the desired moment tracking and muscle activation profiles. | Student successfully used all tools but did not exactly copy all parameters, leading to a solution close to that presented but not identical. | Student did not use Synergy Extrapolation or Muscle-tendon length initialization correctly, leading to a solution that significantly deviated from that presented. |
| Neural Control Model Personalization | Student chose the same number of synergies and achieved the same joint moment tracking and a total muscle VAF > 95%. | Student chose a different number of synergies, but their NCP run gave the intended result for that number of synergies. | Student did not achieve a valid NCP result for varying reasons. These students all had very low total muscle VAFs. |
| Synergy-driven Tracking Optimization | Converged solution produced a ground reaction X moment nearly identical to that presented. | Solutions were close but featured a noticeably different shape of the ground reaction X moment. | Tracking Optimization did not converge within the specified number of iterations. |
| Synergy-driven Design Optimization | Converged solution produced symmetric propulsive and braking impulses within 0.5 Ns of those presented. | Converged solution produced symmetric propulsive and braking impulses to within 1 Ns of those presented. | Verification or Design Optimization did not converge within the specified number of iterations. |

#### Results

Results are presented for all modules in Figures D1-D8. Figures D1-D4 present results for the first tutorial, while Figures D5-D8 present results for the second tutorial. The number of student groups is listed in the title of each figure. The number of student groups changes between figures due to students either changing their groups or not properly submitting deliverables, in which case the group was assigned to the “Not Reproduced” category.

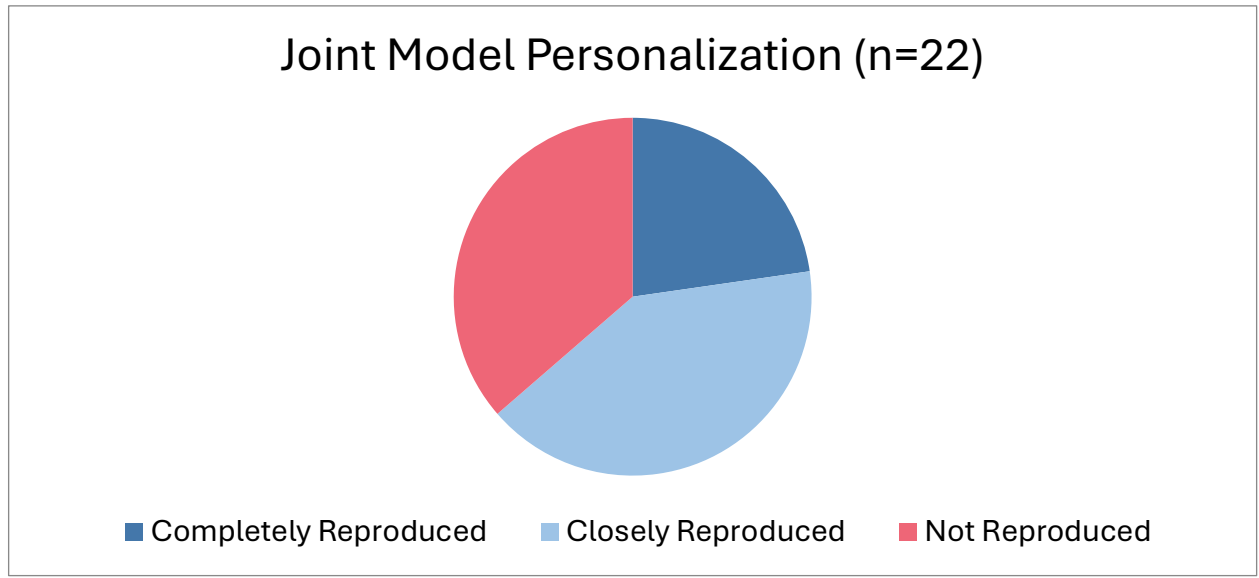

**Figure D1:** Five student groups were able to completely reproduce the authors' Joint Model Personalization results, 9 student groups were able to closely reproduce the authors' results with small differences, and 8 student groups did not reproduce the authors' results.

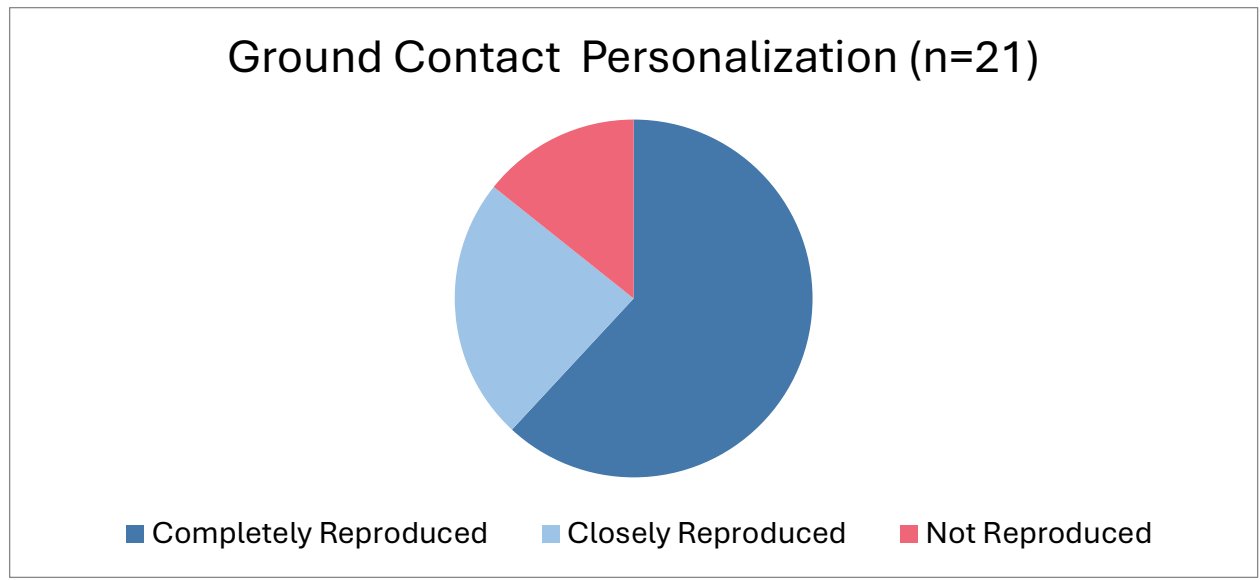

**Figure D2:** Thirteen student groups were able to completely reproduce the authors' Ground Contact Model Personalization results, 5 student groups were able to closely reproduce the authors' results with small differences, and 3 student groups did not reproduce the authors' results.

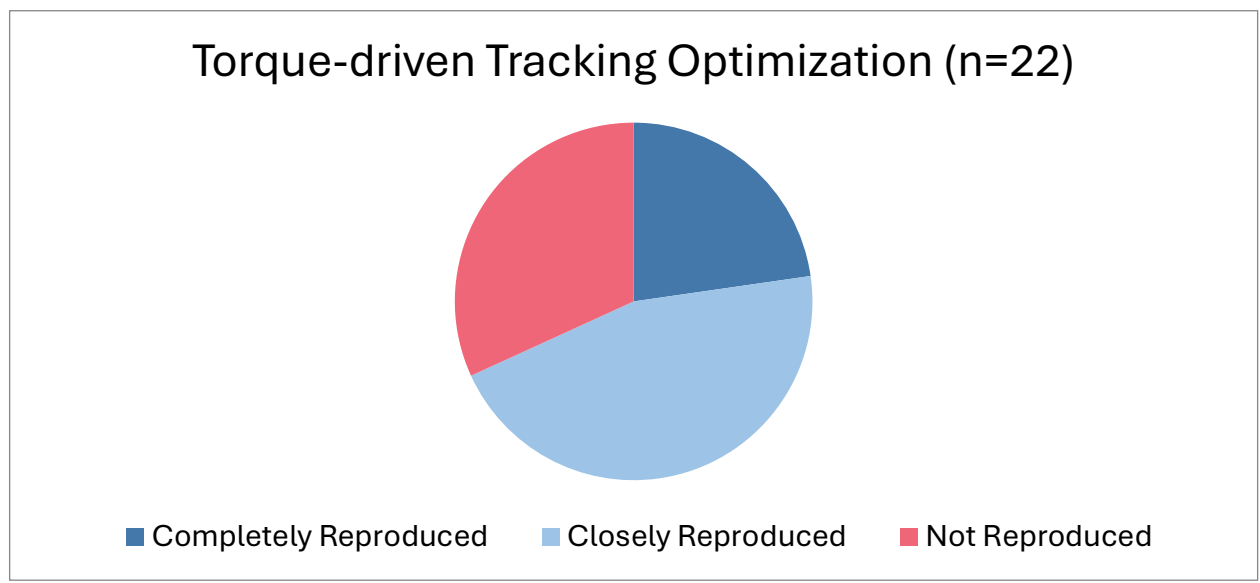

**Figure D3:** Five student groups were able to completely reproduce the authors' torque-driven Tracking Optimization results, 10 student groups were able to closely reproduce the authors' results with small differences, and 7 student groups did not reproduce the authors' results.

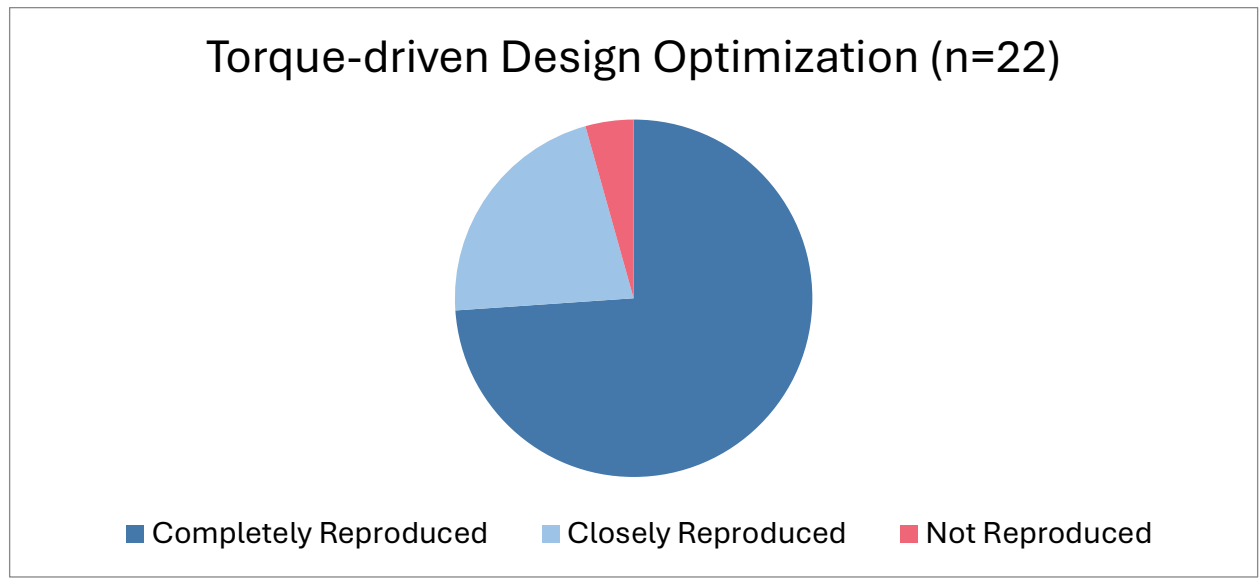

**Figure D4:** Seventeen student groups were able to completely reproduce the intended torque-driven Design Optimization results, 5 student groups were able to closely reproduce the authors' result with small differences, and 1 student group did not reproduce the authors' results.

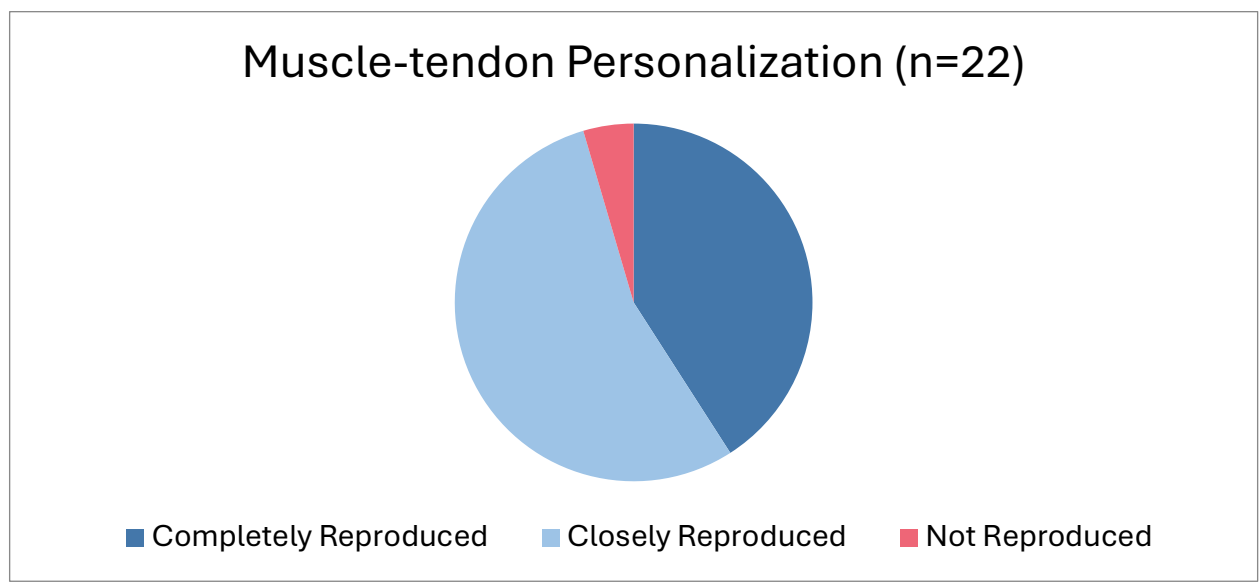

**Figure D5:** Nine student groups were able to completely reproduce the authors' Muscle-tendon Model Personalization results, 11 student groups were able to closely reproduce the authors' results with small differences, and 1 student group did not reproduce the authors' results.

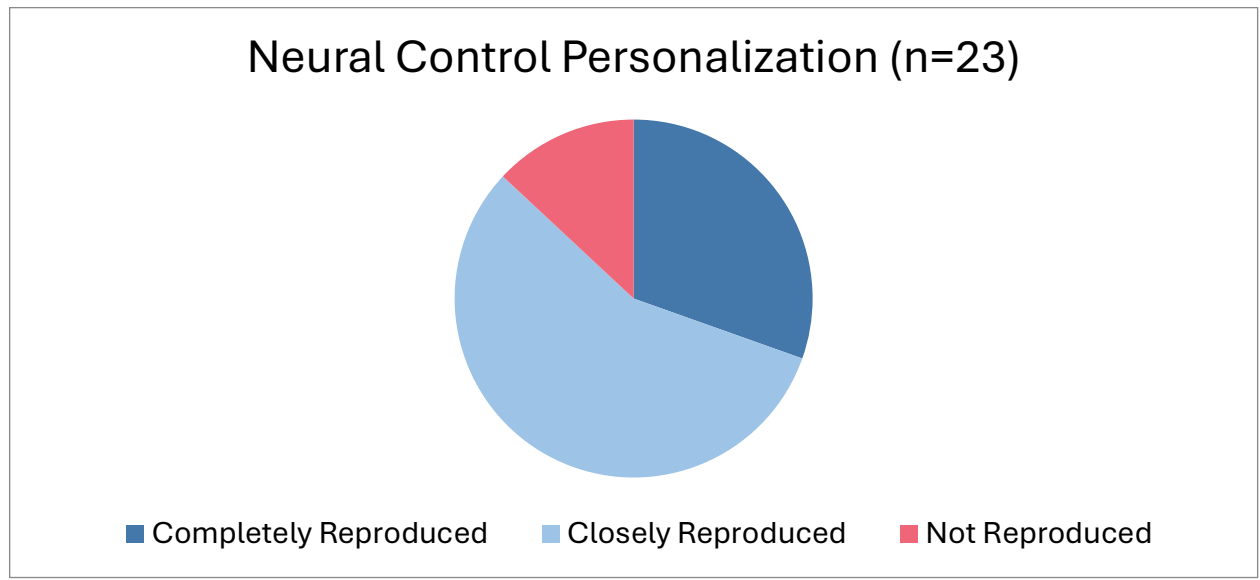

**Figure D6:** Seven student groups were able to completely reproduce the authors' Neural Control Model Personalization results, 13 student groups were able to closely reproduce the authors' results with small differences, and 3 student groups did not reproduce the authors' results.

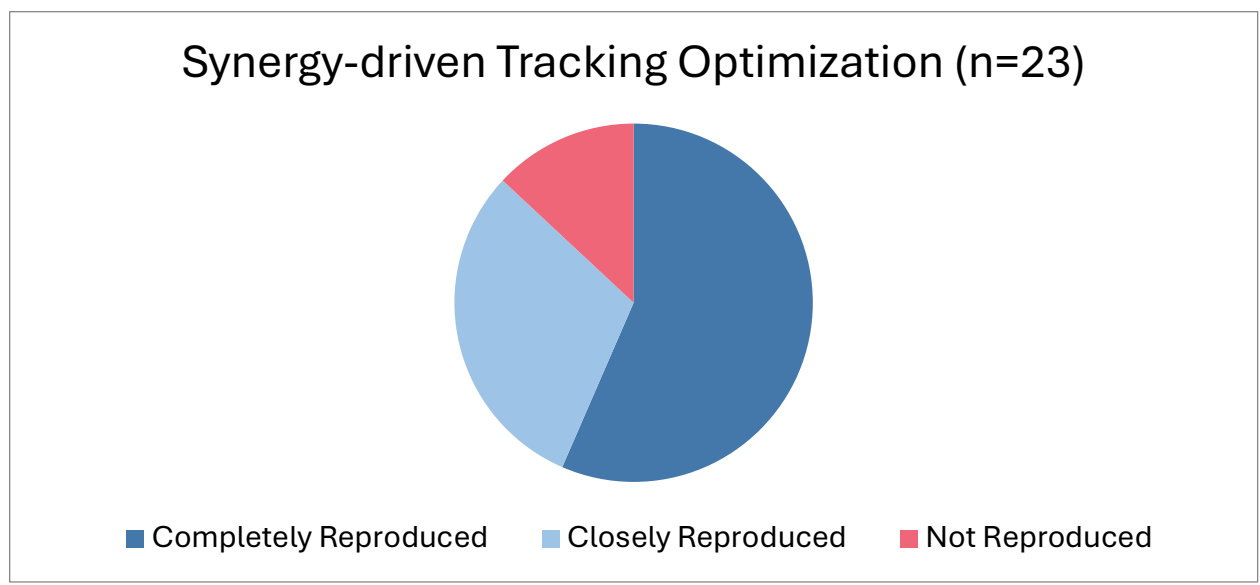

**Figure D7:** Thirteen student groups were able to completely reproduce the authors' Synergy-driven Tracking Optimization results, 7 student groups were able to closely reproduce the authors' results with small differences, and 3 student groups did not reproduce the authors' results.

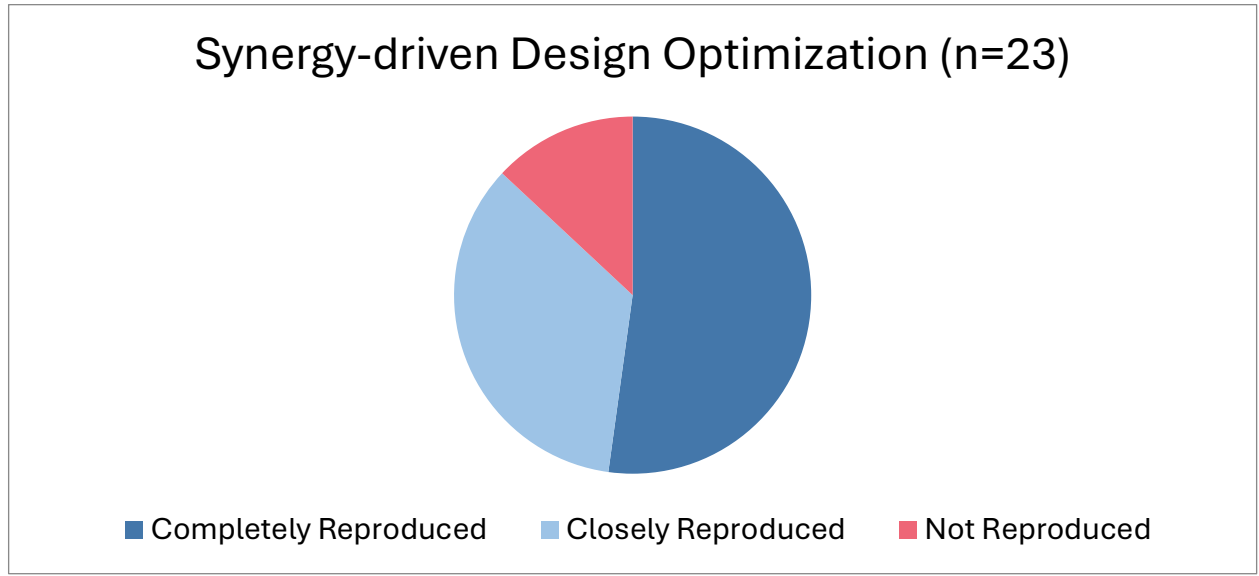

**Figure D8:** Twelve student groups were able to completely reproduce the authors' Synergy-driven Design Optimization results, 8 student groups were able to closely reproduce the authors' results with small differences, and 3 student groups did not reproduce the authors' result.

#### Discussion

Across all modules, the majority of student submissions either completely or closely reproduced the authors' results. When a submission fell into the category of *closely reproduced*, it was always due to small errors or deviations in the problem formulations, but the results still closely resembled the final results presented in this article. When a submission fell into the category of *not reproduced*, it was primarily because the student produced a significantly different solution from the results presented in this article. For these cases, it was not entirely clear where student made errors that caused their solutions to deviate. In the following paragraphs, the reproducibility of each module within each tutorial is discussed in more detail.

For the first tutorial, student groups performed Joint Model Personalization, Ground Contact Model Personalization, torque-driven Tracking Optimization, Verification Optimization, and Design Optimization. Joint Model Personalization featured the largest number of students who could not reproduce the authors' results, and this module featured the lowest number of students who perfectly reproduced the final results. One likely reason for this lower reproducibility is that students were given more freedom for how to approach the Joint Model Personalization process. Students were not explicitly instructed on which model markers they should use in their Joint Model Personalization tasks, nor were they explicitly instructed on which marker file they should use. This lack of explicit instruction led to a large distribution of potential solutions that students could achieve. Of the 8 student groups that did not reproduce the authors' results, 4 of them did not choose the same model markers or the same marker file. Another likely reason for lower reproducibility with this module is that it includes using the Model Scale tool in OpenSim, which is known to be challenging to use for beginning users. The 4 student groups that did not reproduce the authors' results and many of the student groups that closely reproduced the authors' results had difficulties with model scaling before Joint Model Personalization was performed, resulting in different marker errors. A final likely reason for lower reproducibility is that this module was the first project assignment that students were given. It may have taken students longer for the first project to understand fully how project tasks and deliverables worked. Overall, reproducibility for the Joint Model Personalization module would have been improved if students had started with a pre-scaled generic OpenSim model and if marker choices had been less open-ended.

Ground Contact Model Personalization had a majority of student groups reproduce the authors' results. Of the student groups that closely reproduced the intended result, most followed all instructions exactly but forgot to unlock the toes joint for inverse kinematics. The students that did not reproduce the authors' results struggled in various ways that are difficult to diagnose purely from viewing plots. Overall, the Ground Contact Model Personalization module was exceptionally reproducible.

Torque-driven Tracking Optimization featured similar levels of reproducibility as Joint Model Personalization. Seven students did not reproduce the authors' results because their optimizations did not converge. The reasons for nonconvergence varied between student groups but were primarily related to students not using the correct input data. Of the 10 student groups that closely reproduced the final result, 7 used an incorrect ground reactions data file and converged to a very good solution, but it had slightly different ground reaction moments compared to the authors' solution. The remaining 3 student groups obtained high quality results but with different x moments than the authors' results.

Torque-driven Design Optimization was exceptionally reproducible with 74% of student groups perfectly reproducing the authors' results for their chosen treatment design. There were no clear trends among the student groups that closely reproduced the authors' results. The reason for higher reproducibility of this module compared to the Tracking Optimization module is that the students were given correct Tracking Optimization results to use as a starting point for Verification and Design Optimization.

For the second tutorial, student groups performed Muscle-tendon Model Personalization, Neural Control Model Personalization, synergy-driven Tracking Optimization, Verification Optimization, and Design Optimization. Muscle-tendon Model Personalization featured 41% of students completely reproducing the authors' solution and 55% of students closely reproducing the authors' solution. For all solutions that closely reproduced the authors' solution, the key discrepancies were small differences in muscle activations. These differences were likely caused by student groups not using the exact cost term maximum allowable errors and error centers that were listed in the tutorial document. While all cost term parameters were listed in the tutorial document, it is understandable that students could miss a single cost term value and get a different solution. Only one student group got results that were significantly different from the authors' results, suggesting that the Muscle-tendon Model Personalization module is highly reproducible.

The Neural Control Model Personalization module had 55% of student groups in the *closely reproduced* category because students were given freedom for choosing how to approach the problem. Students were encouraged to explore solutions with different numbers of synergies. Student groups that completely reproduced the authors' results all decided to submit a solution with four synergies, while those that closely reproduced the authors' results used five synergies – another reasonable choice. Because of the freedom given to students in selecting the number of synergies, the distinction between *mostly reproduced* and *completely reproduced* is misleading for this module.

The synergy-driven Tracking Optimization module had 57% of student groups in the *completely reproduced* category. Among the student groups that were in the *closely reproduced* category, it was not clear where alternative decisions were made, but all of their solutions converged well to an acceptable solution. The student groups in the *not reproduced* category were all assigned to that category because they did not submit plots with their submission, making it impossible to judge to what extent they reproduced the authors' results. However, one of the primary challenges with synergy-driven Tracking Optimization is simply getting the optimization to converge to any solution. Thus, it is likely that these student groups were unable to achieve optimization convergence.

The synergy-driven Design Optimization module had 52% of students in the *completely reproduced* category. Like the Neural Control Model Personalization module, student groups were given significant freedom with how they approached this module. Consequently, reproducibility was judged based on whether they achieved the target values for braking and propulsive impulses. Most student groups in the *closely reproduced* category struggled with generating Verification Optimization results, but they still achieved the intended changes to propulsive and braking impulses. Student groups in the *not reproduced* category either did not submit plots or their Design Optimization runs never converged. Reproducibility of Verification Optimization was not analyzed because it is inherently tied to Design Optimization – If students did not reproduce Verification Optimization, they did not reproduce Design Optimization either. Overall, 87% of students were able to use Design Optimization to achieve the desired correction to the subject's propulsive and braking impulses.

Overall, 64% of student groups were able to completely or closely reproduce the authors' results by following the written instructions in these two tutorials. Although we made a distinction between *completely reproduced* and *closely reproduced*, all solutions in the *closely reproduced* category were only slightly different from those in the *completely*

*reproduced* category. In many cases, a single parameter change was the difference between these two categories. The two modules with the lowest reproducibility were Joint Model Personalization and torque-driven Tracking Optimization. Students seemed to struggle more with these modules primarily because of flexibility provided by the tutorials regarding which input data files should be used. The next time the course is taught, we plan to update the instructions for these modules to state explicitly which input files should be used. With that one small change, we anticipate that these modules will have similar levels of reproducibility to the other modules.

In conclusion, the research presented in this article was demonstrated to be organically reproducible based on an evaluation of data collected from student groups in a combined undergraduate/graduate course taught at Rice University. One limitation of this evaluation is that students in the course had access to the instructor and teaching assistant through email and office hours. Although users at other institutions who work through these tutorials will not have the same access, the NMSM Pipeline has a Simtk.org user forum where users can and do post questions and receive help when working through NMSM Pipeline tutorials.
